## Supplementary figures and images for "Improving the coverage of credible sets in Bayesian genetic fine-mapping"

### rs151234.png

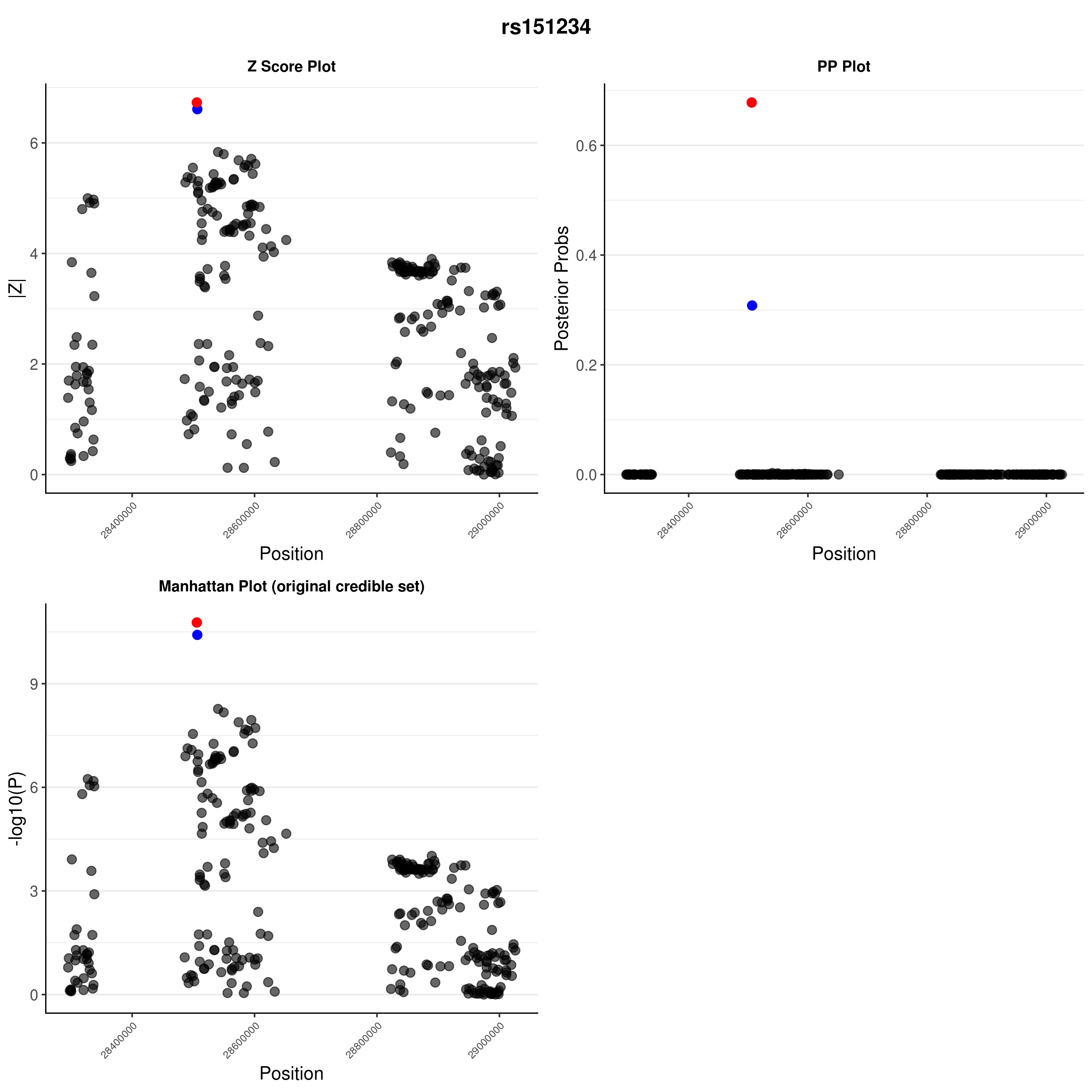

### rs151234.png

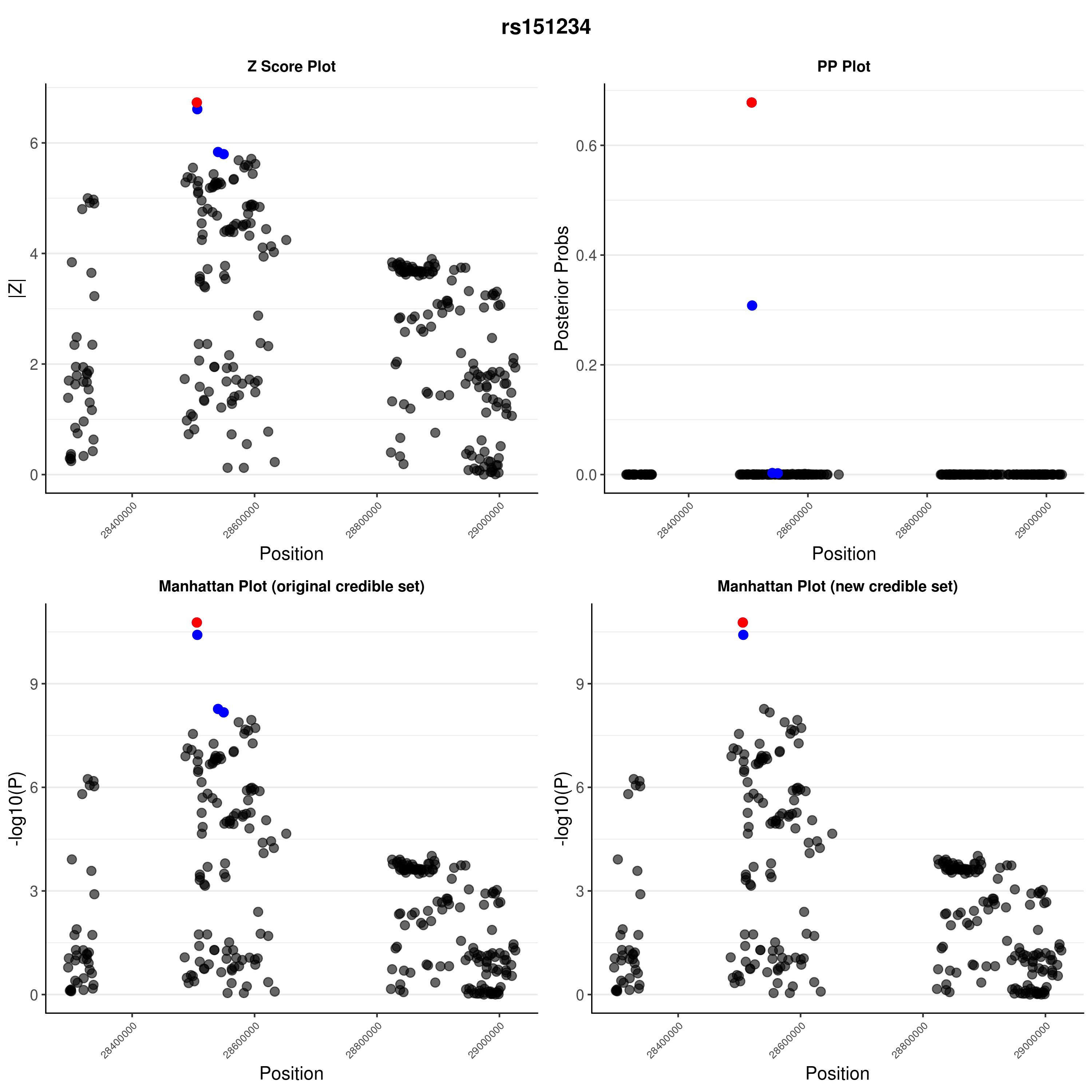

### rs193778.png

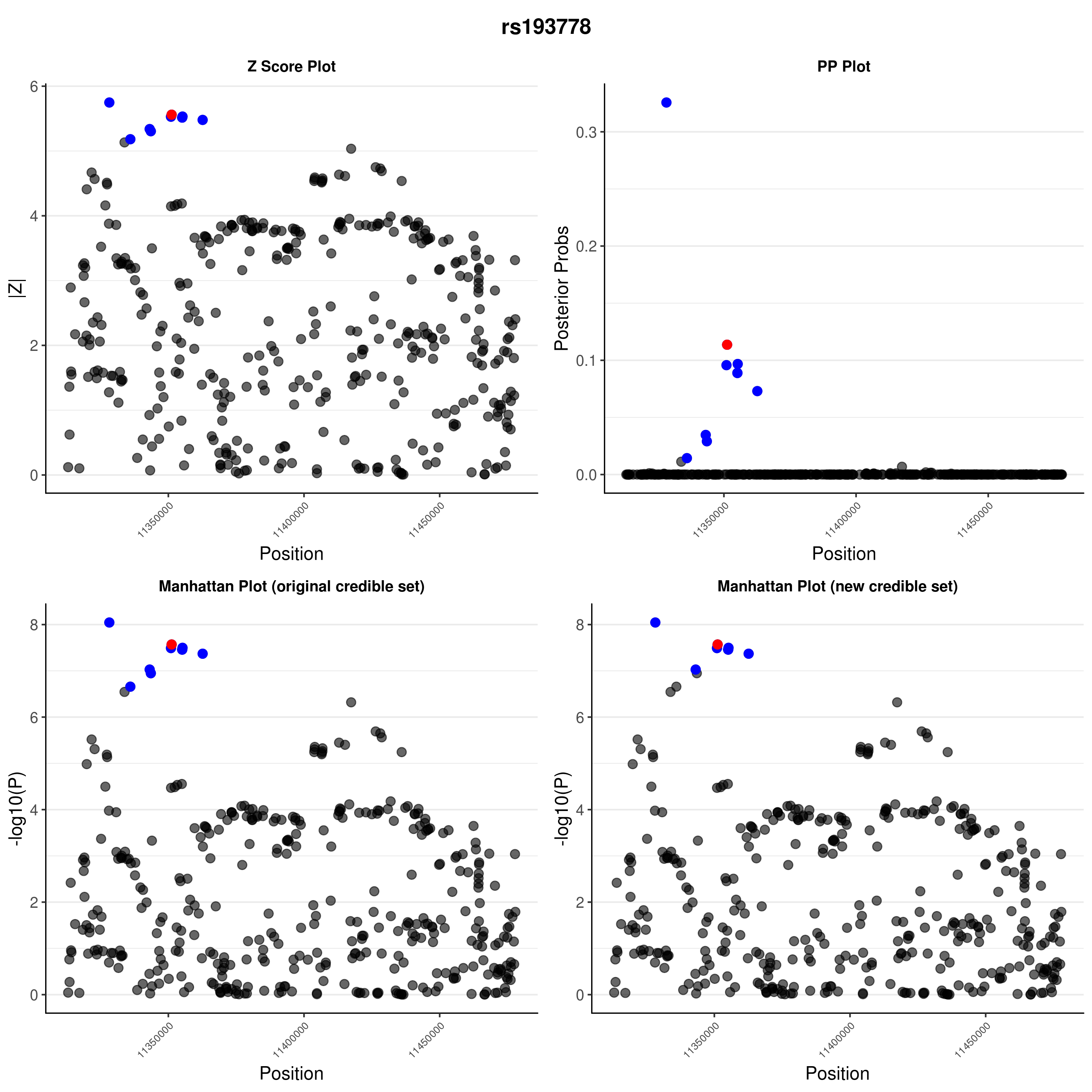

### rs193778.png

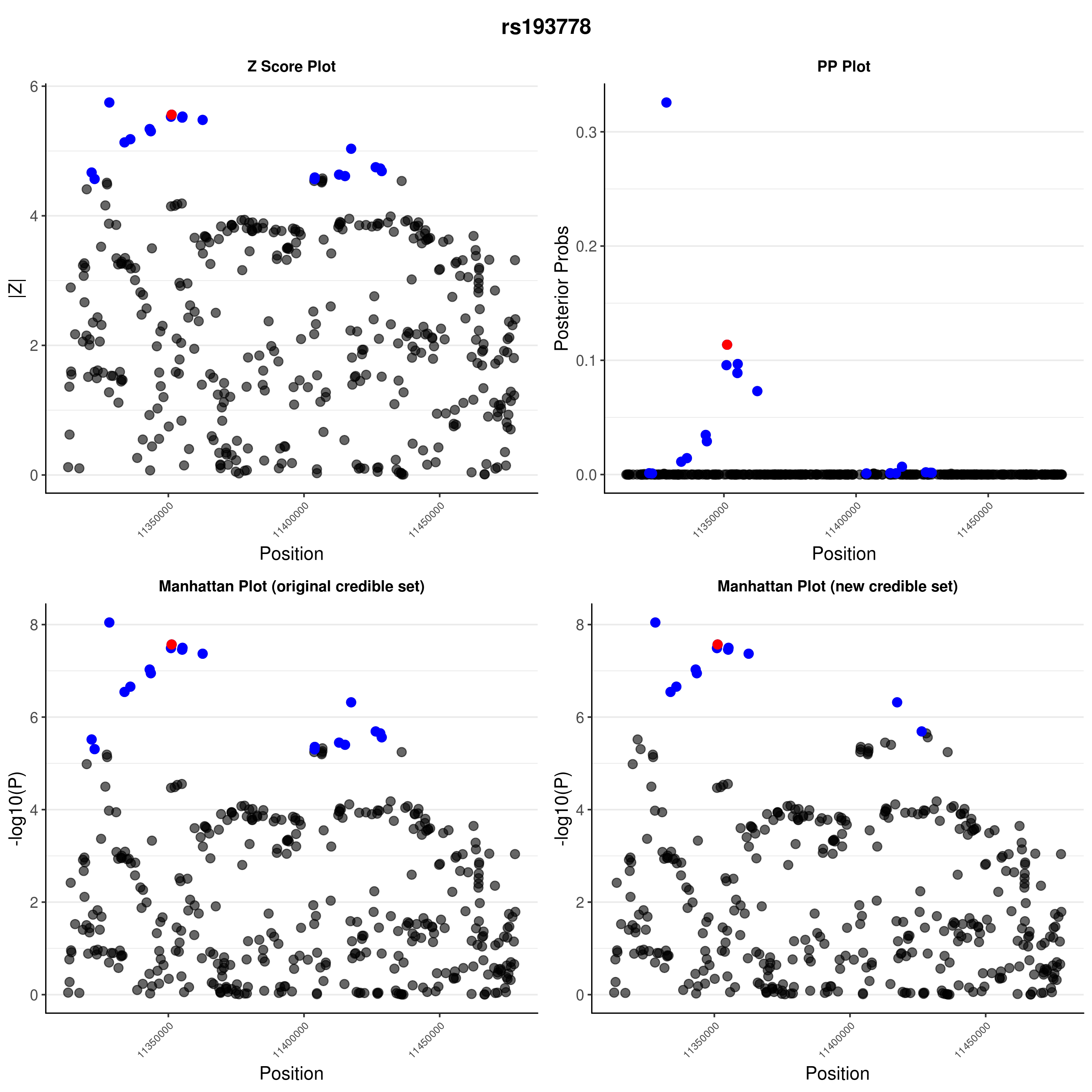

### rs229533.png

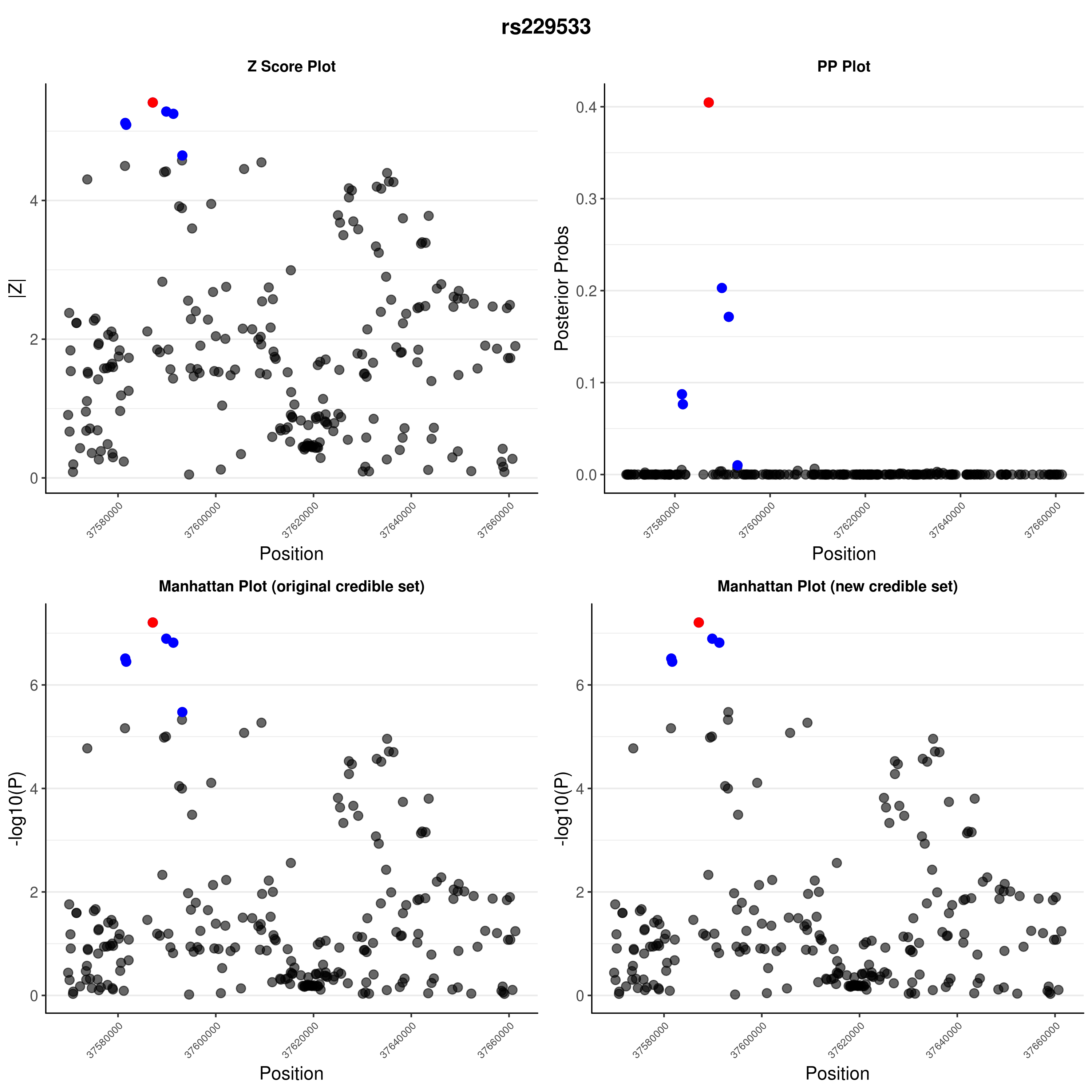

### rs229533.png

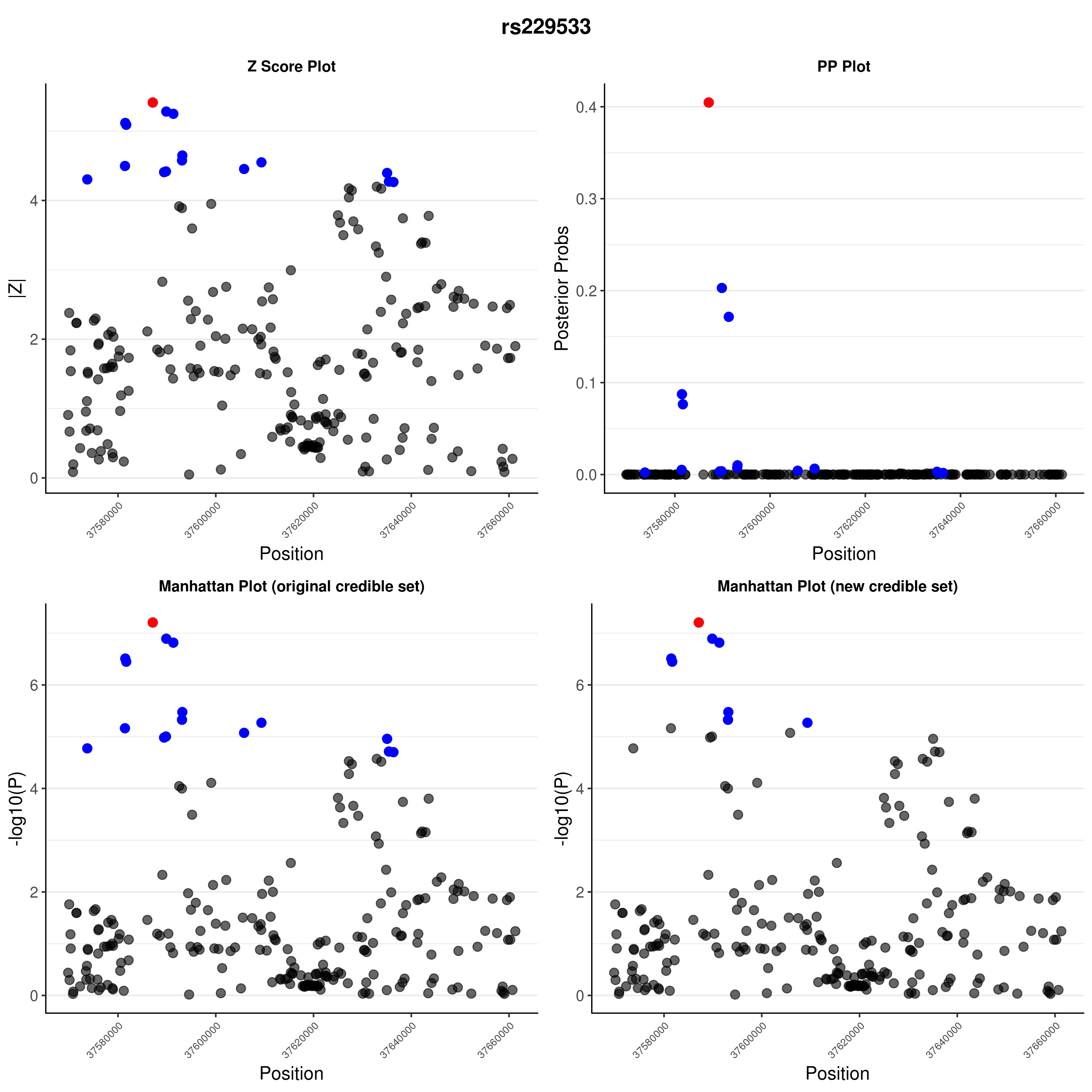

### rs402072.png

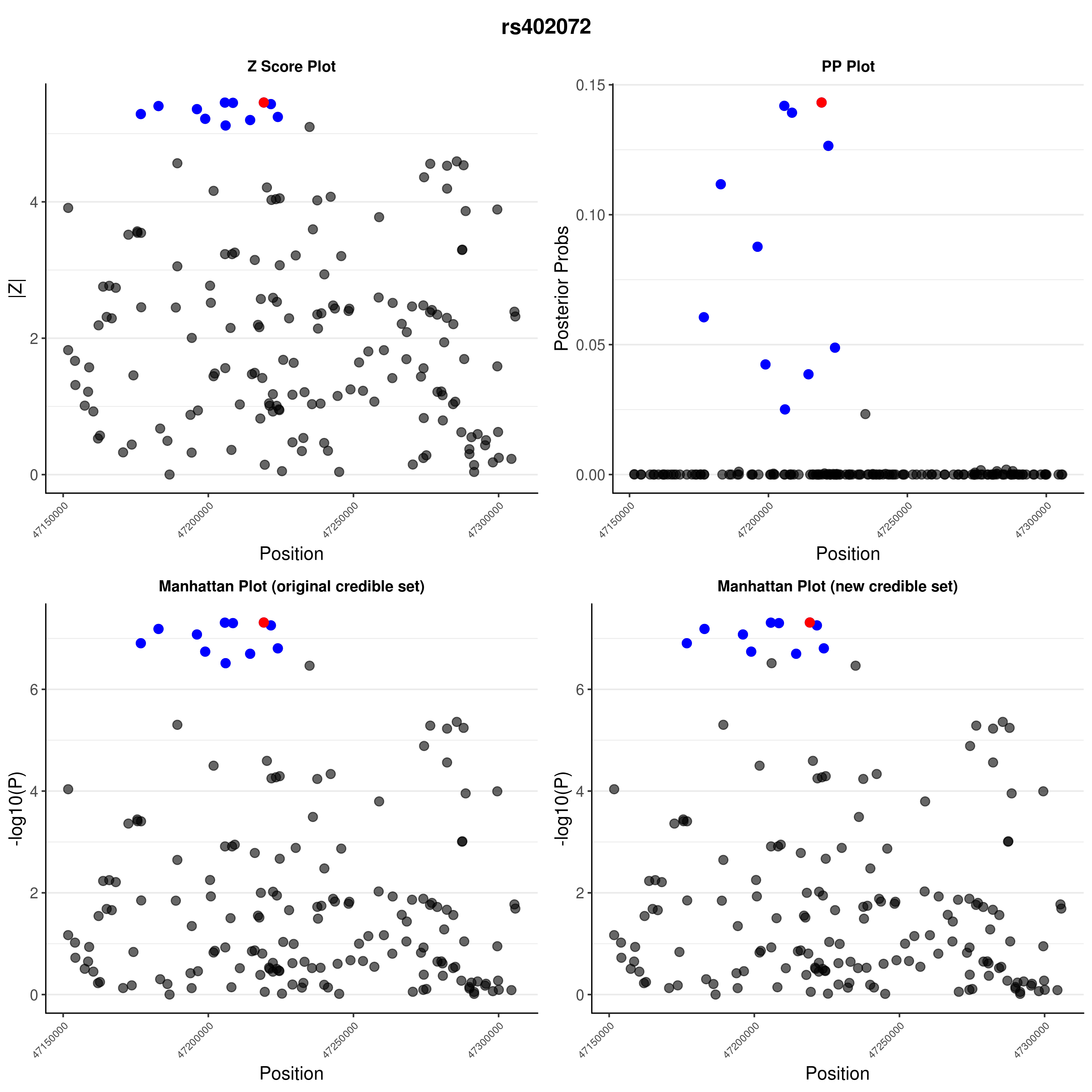

### rs402072.png

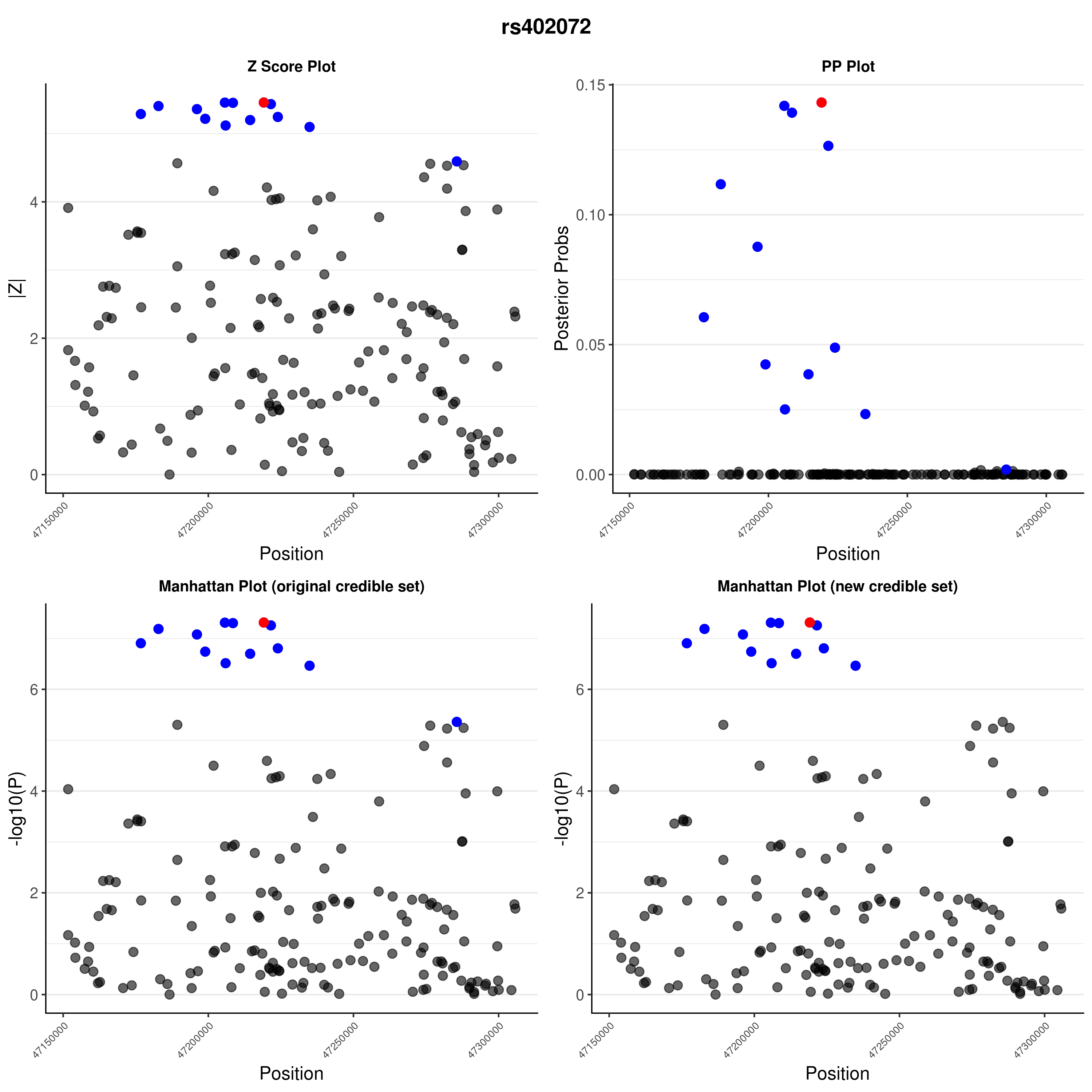

### rs516246.png

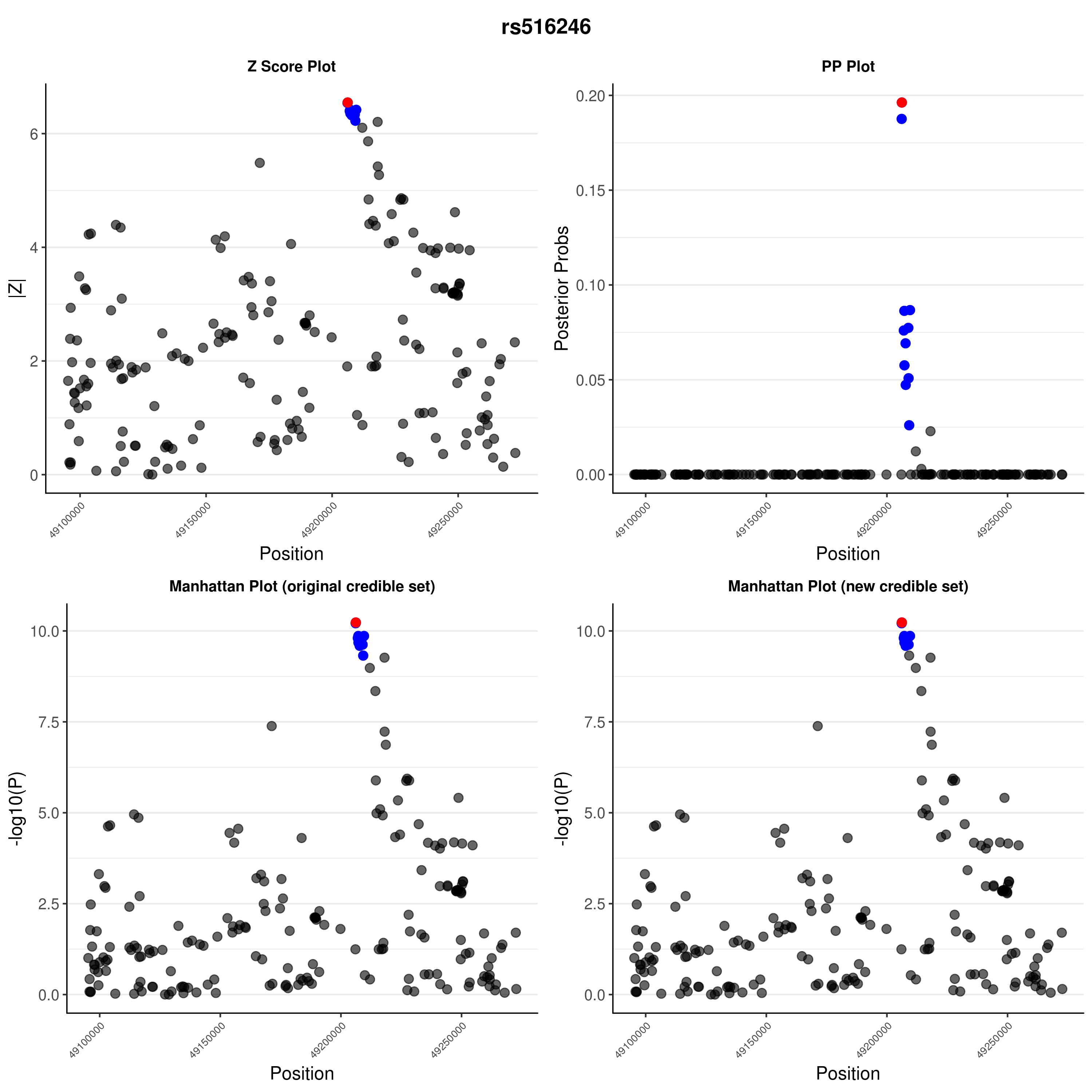

### rs516246.png

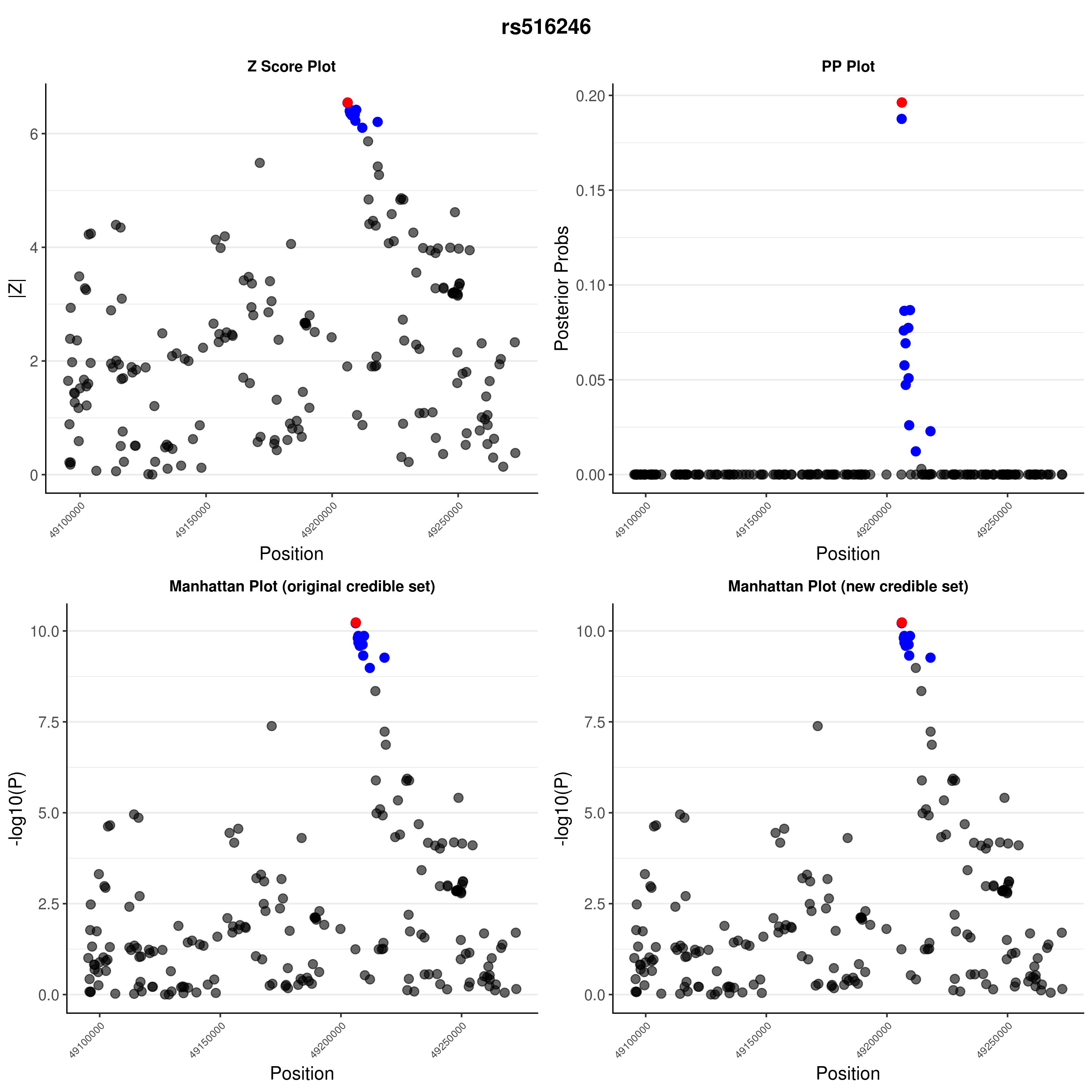

### rs653178.png

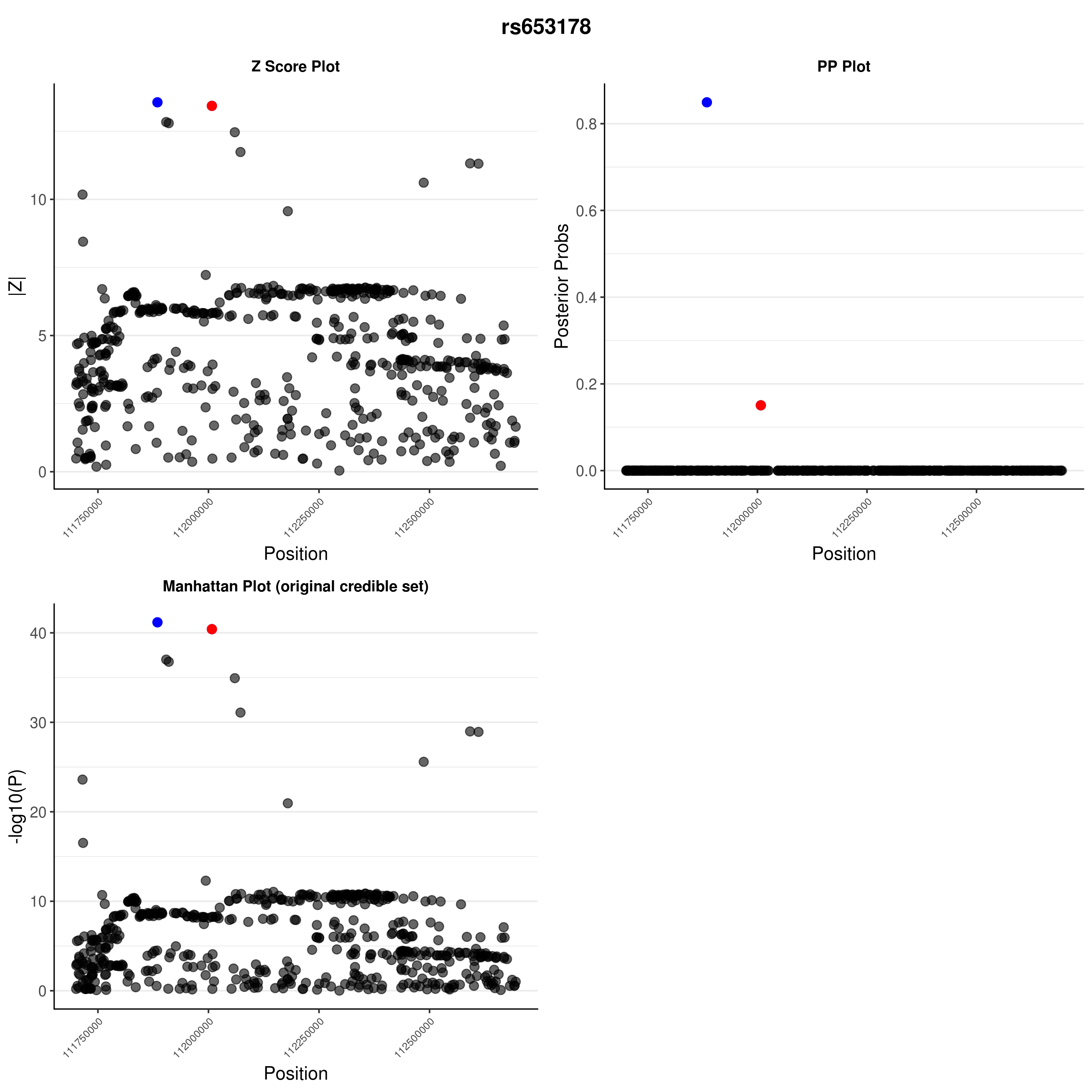

### rs705705.png

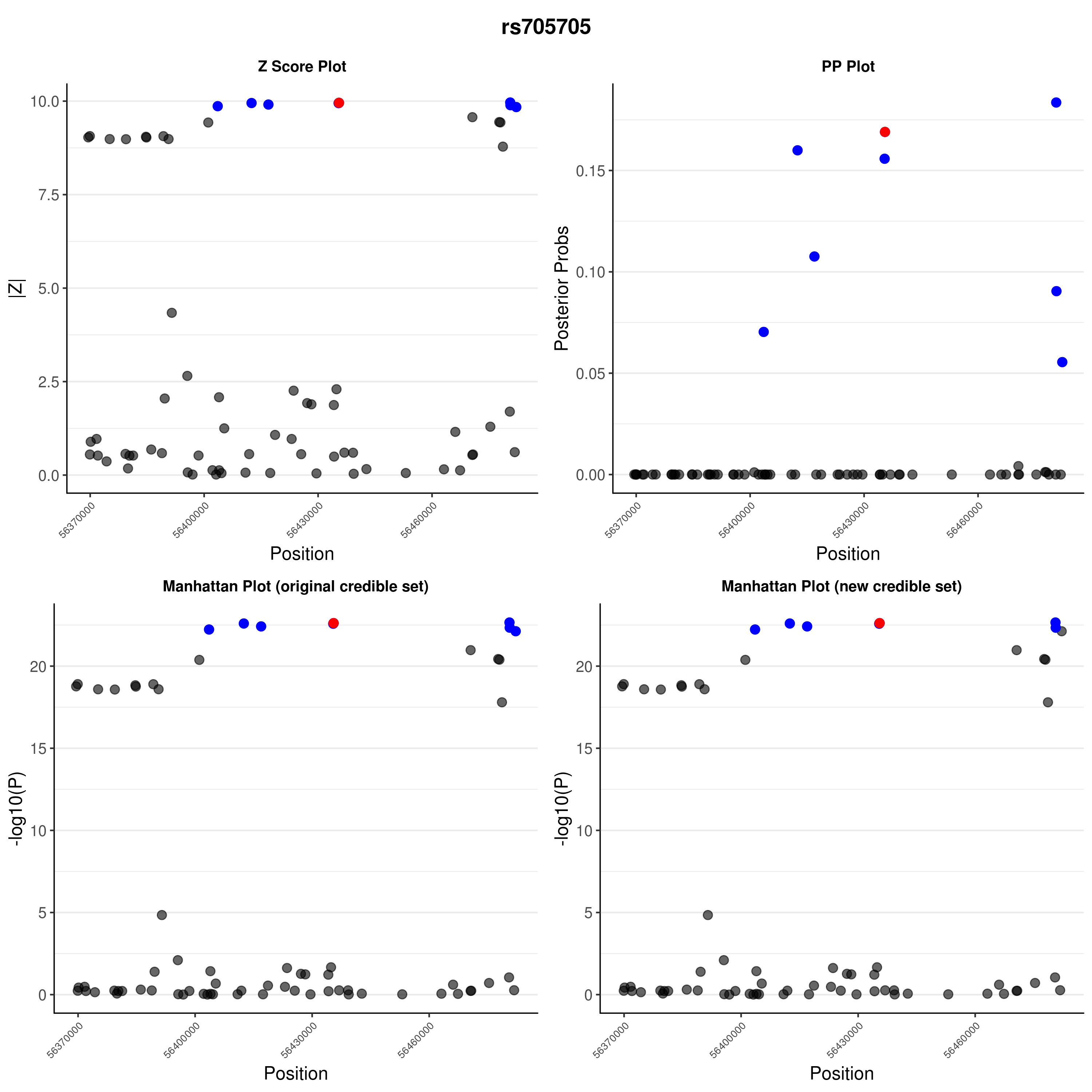

### rs705705.png

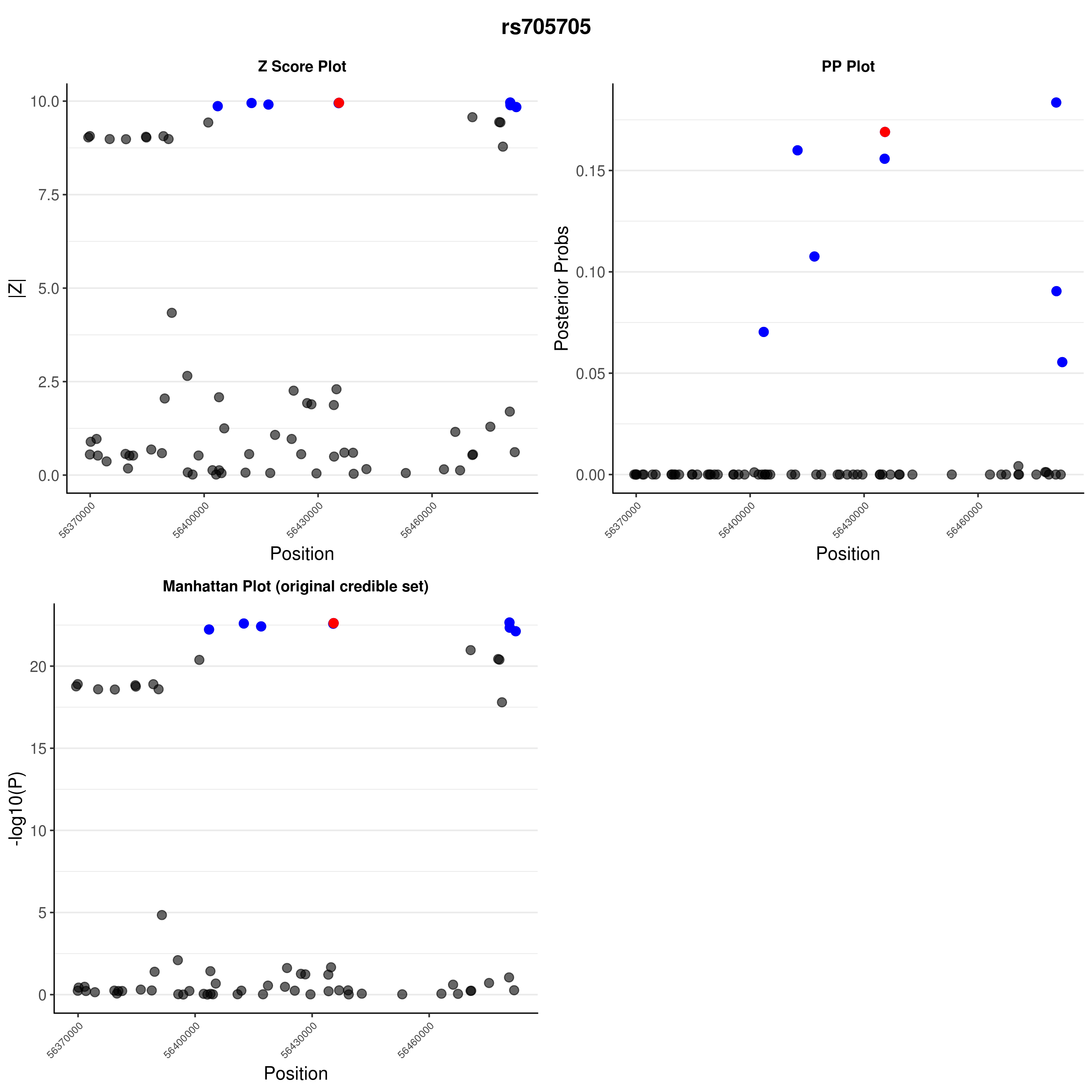

### rs757411.png

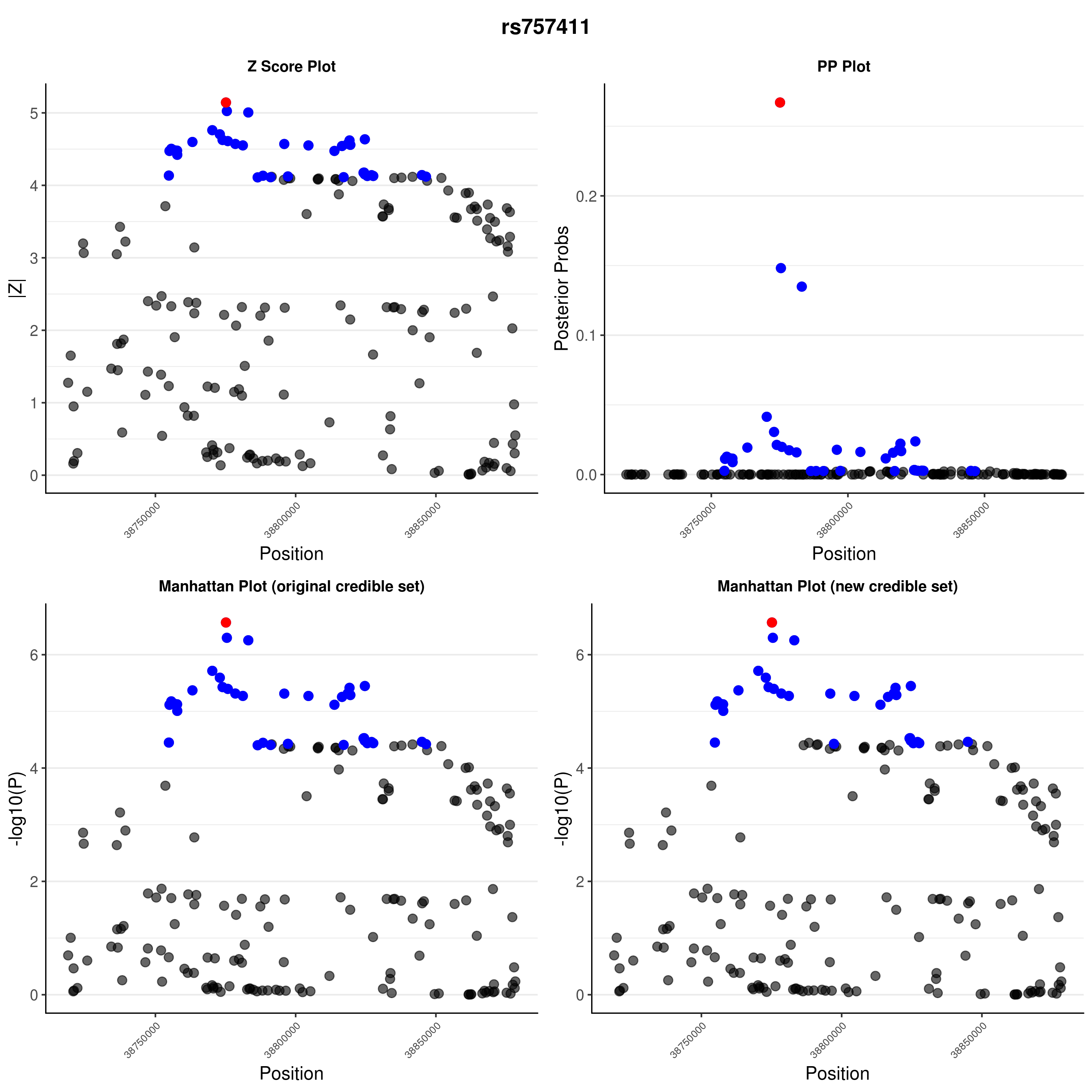

### rs757411.png

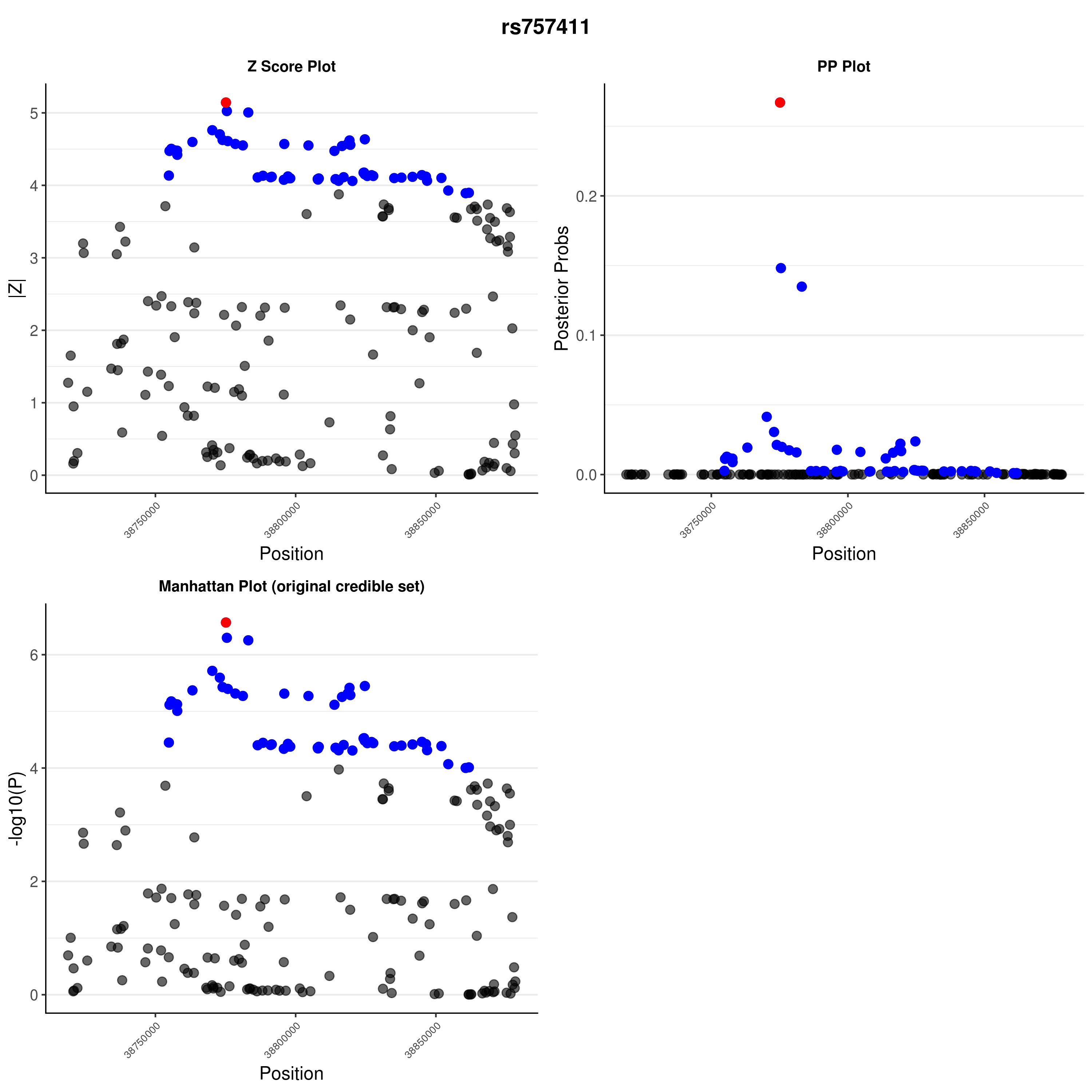

### rs917911.png

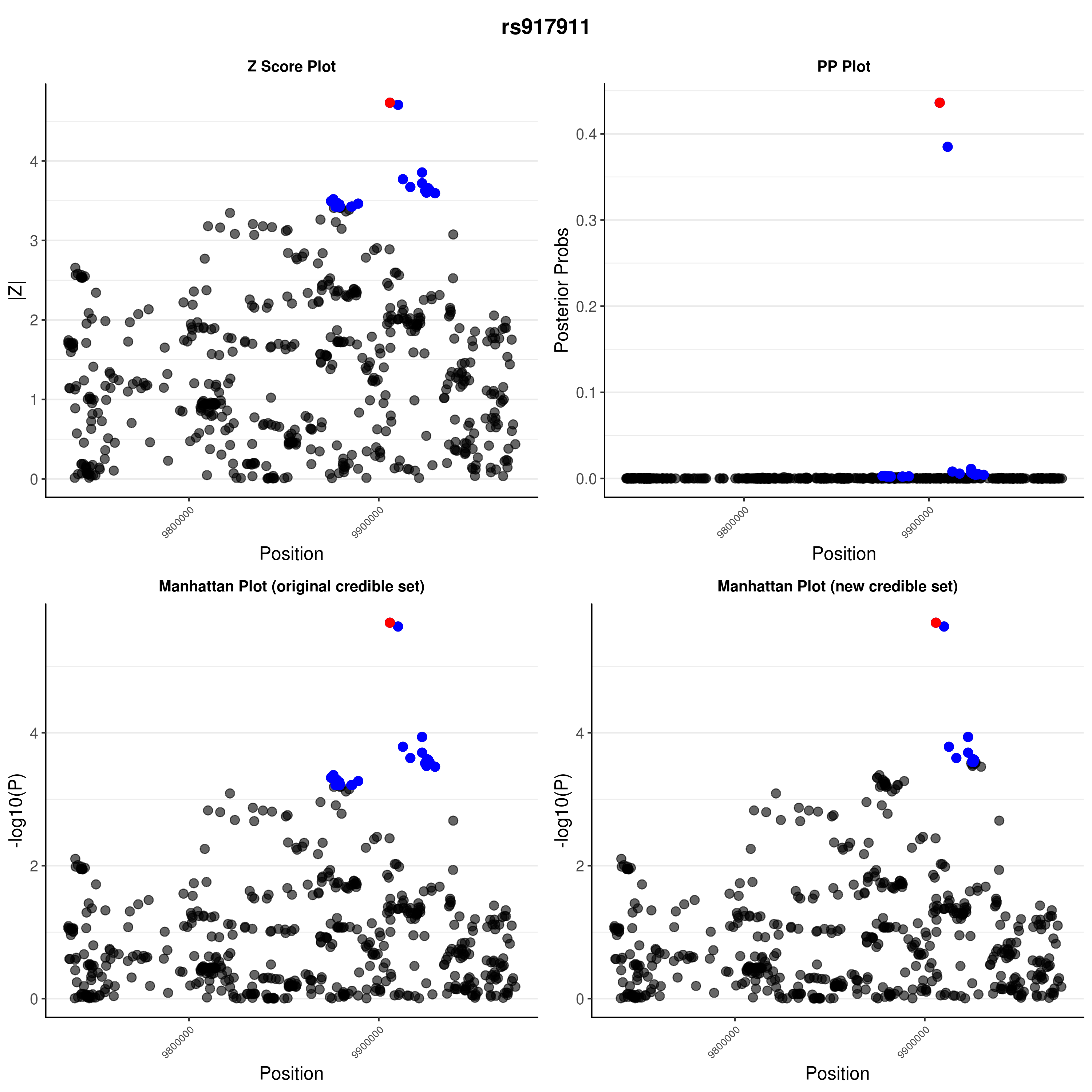

### rs917911.png

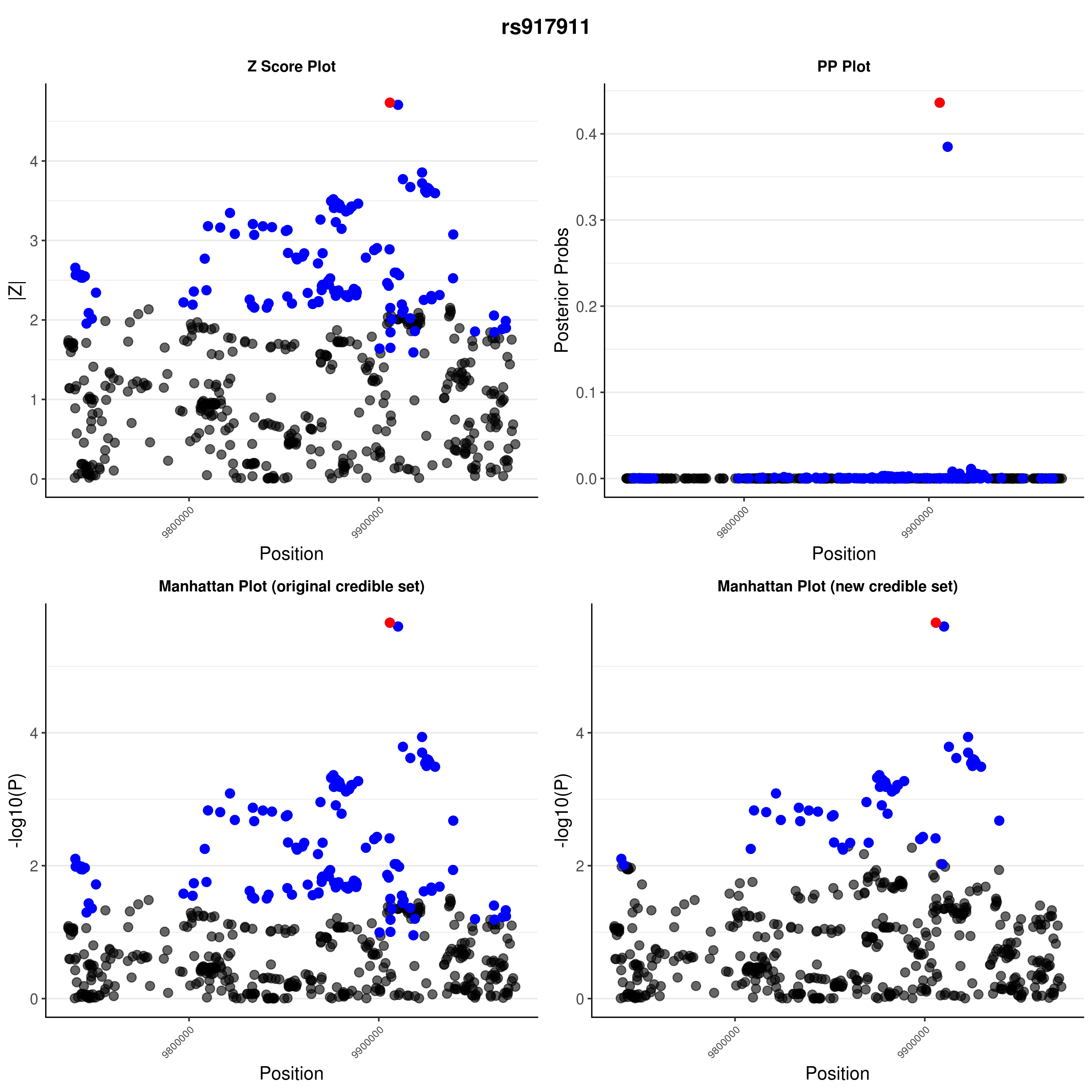

### rs1456988.png

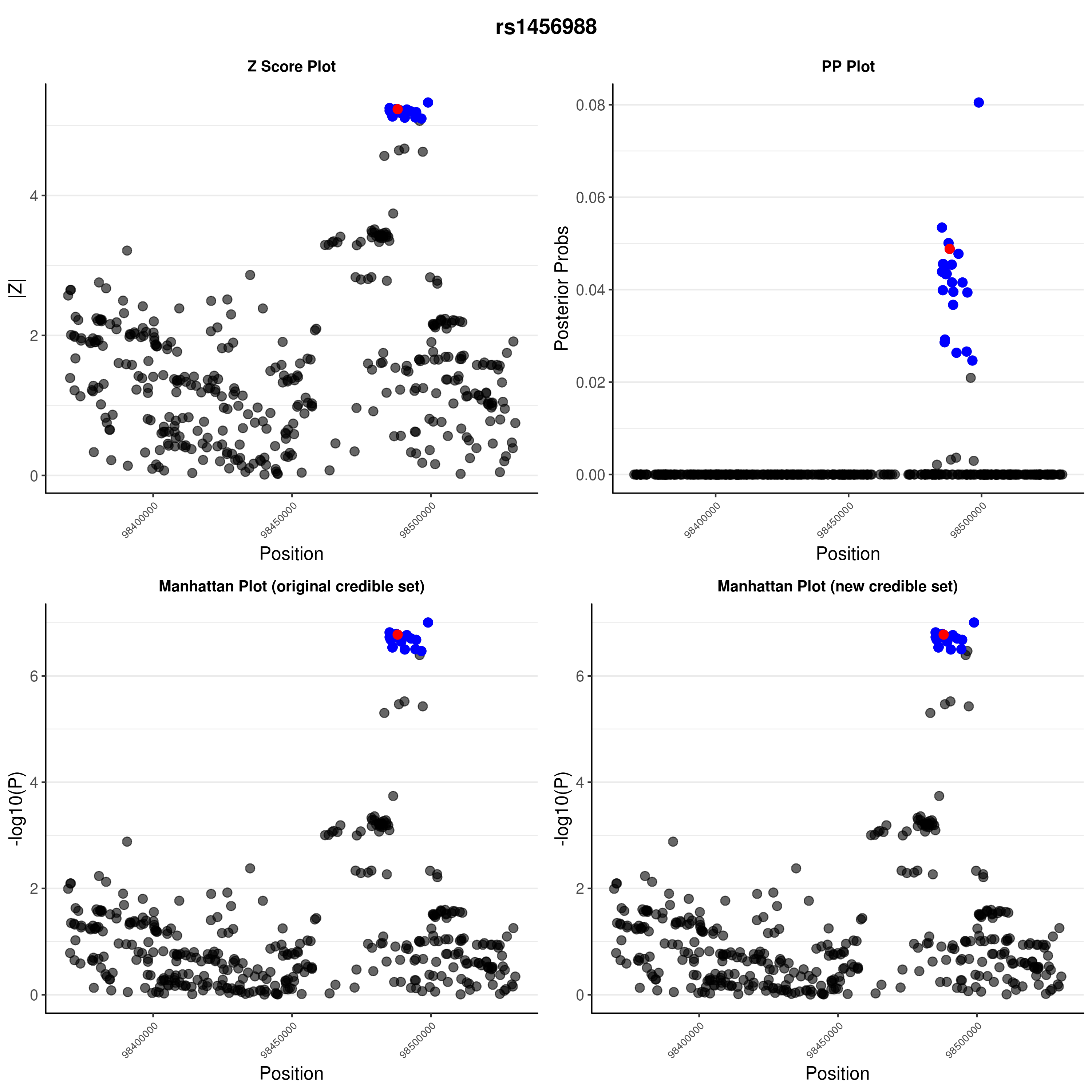

### rs1456988.png

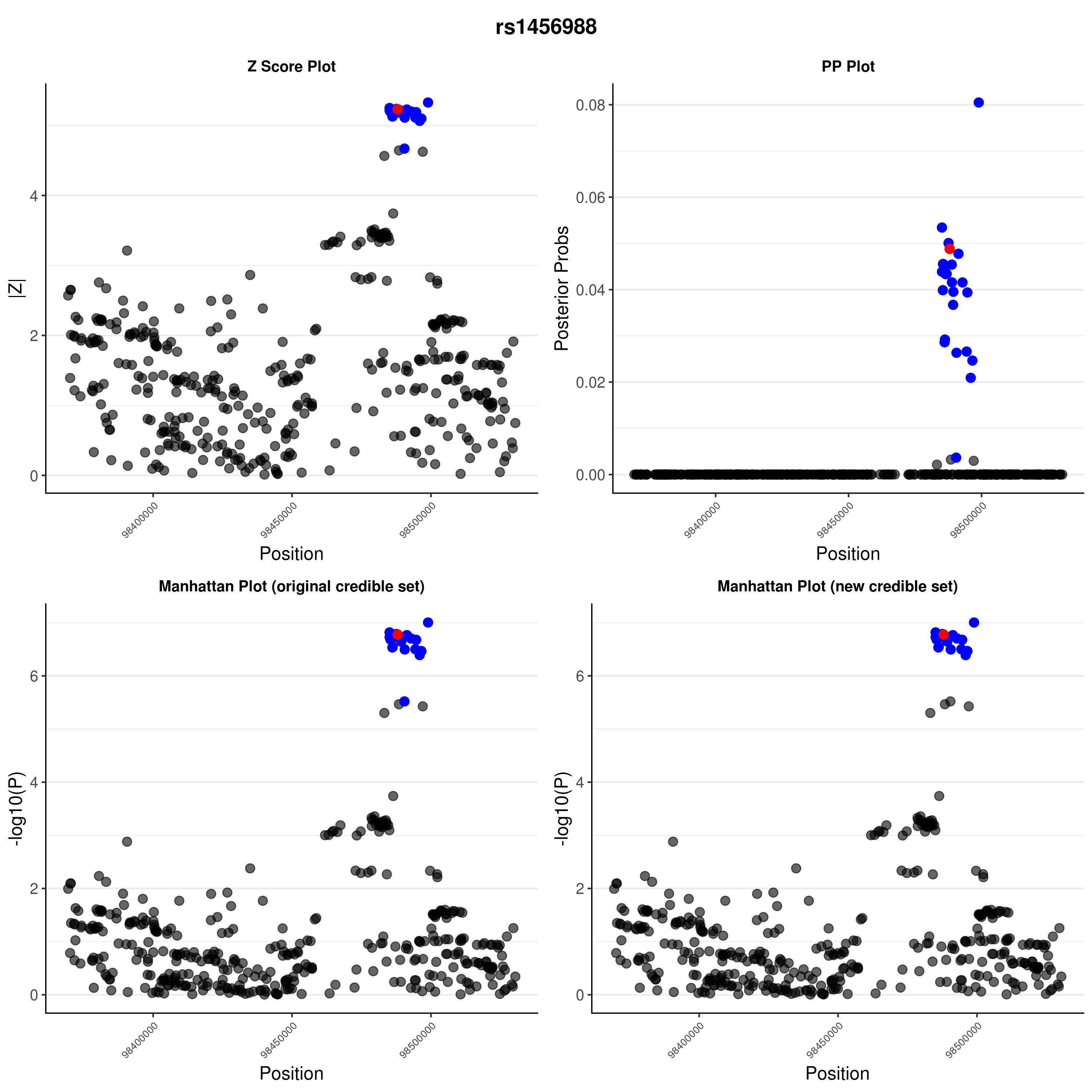

### rs1538171.png

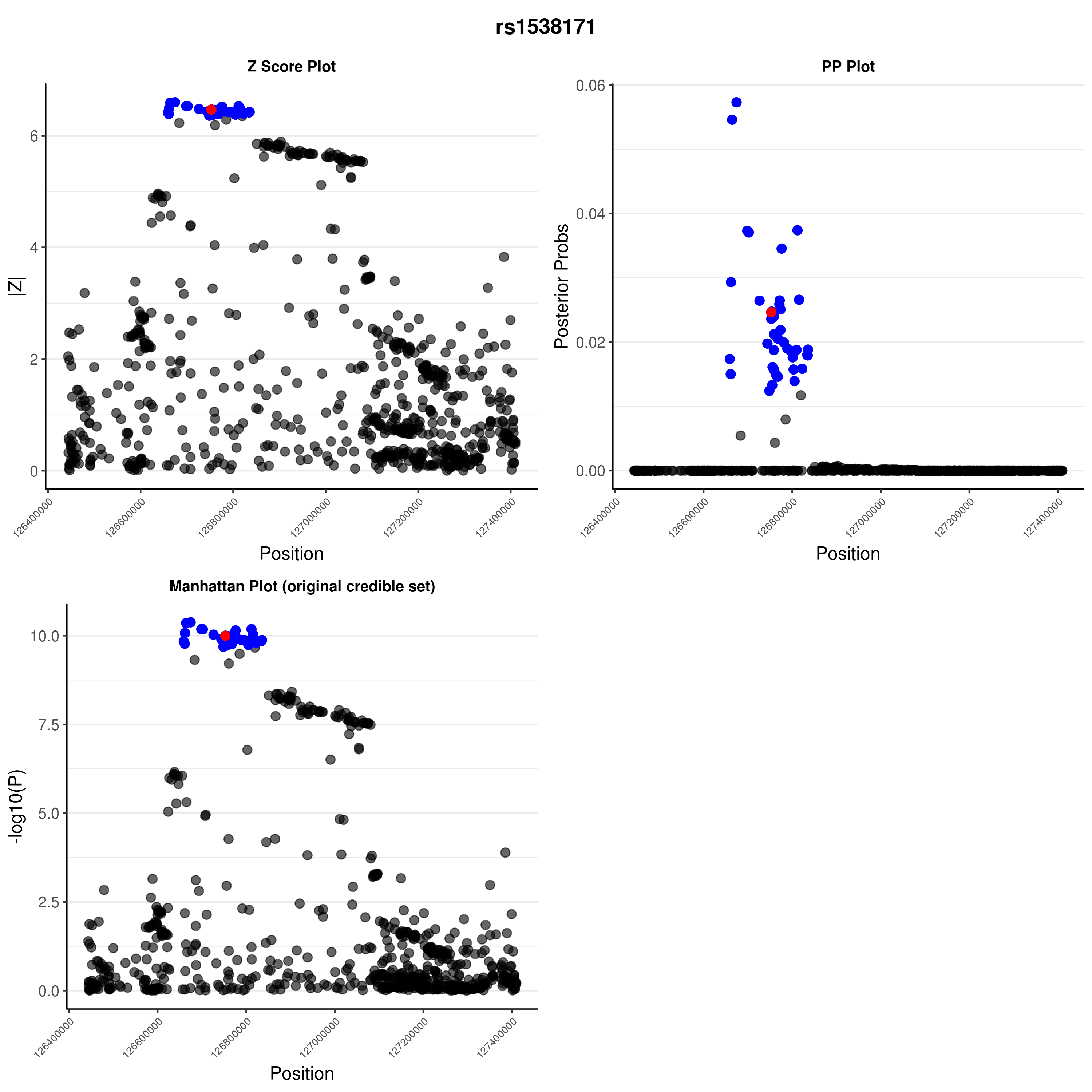

### rs1538171.png

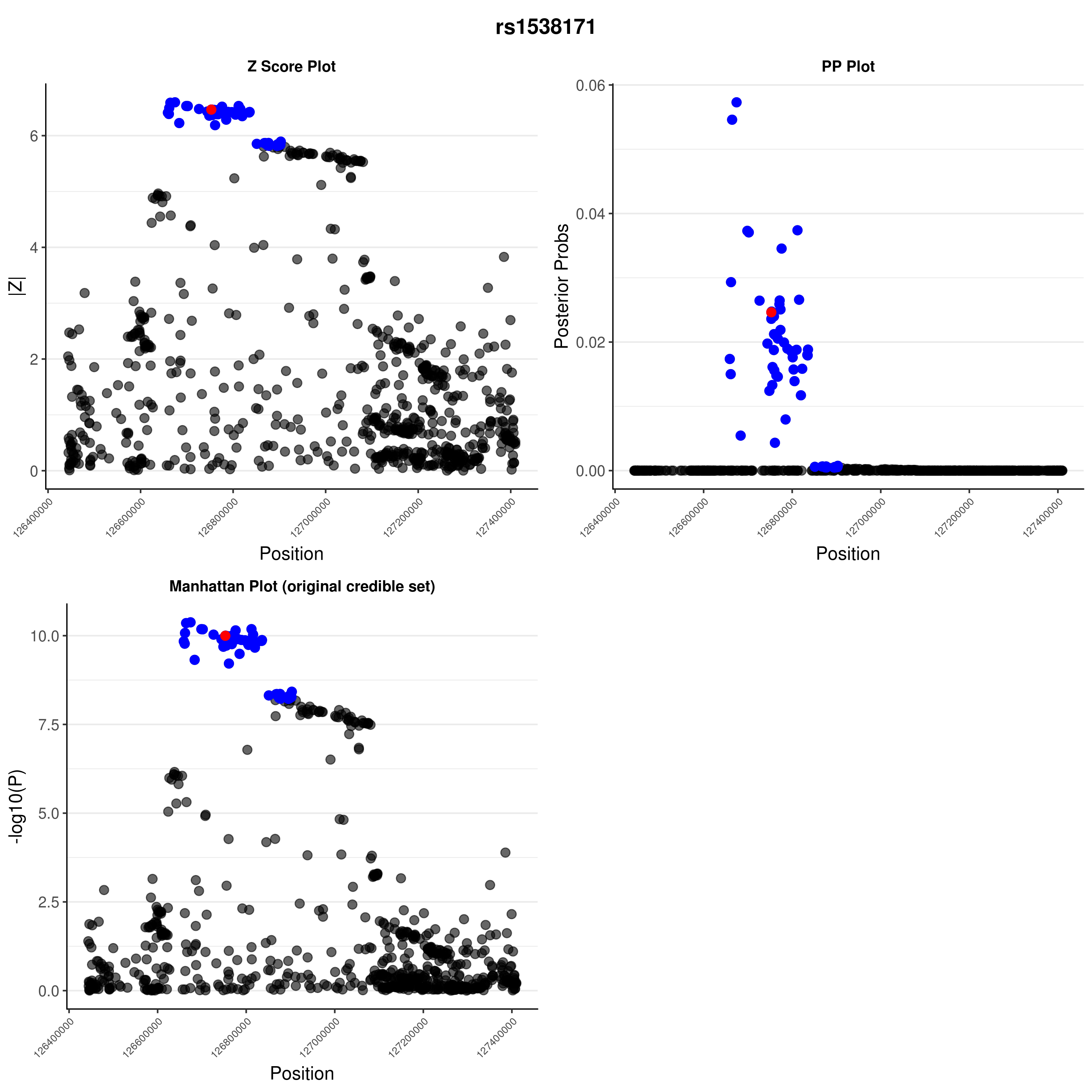

### rs1615504.png

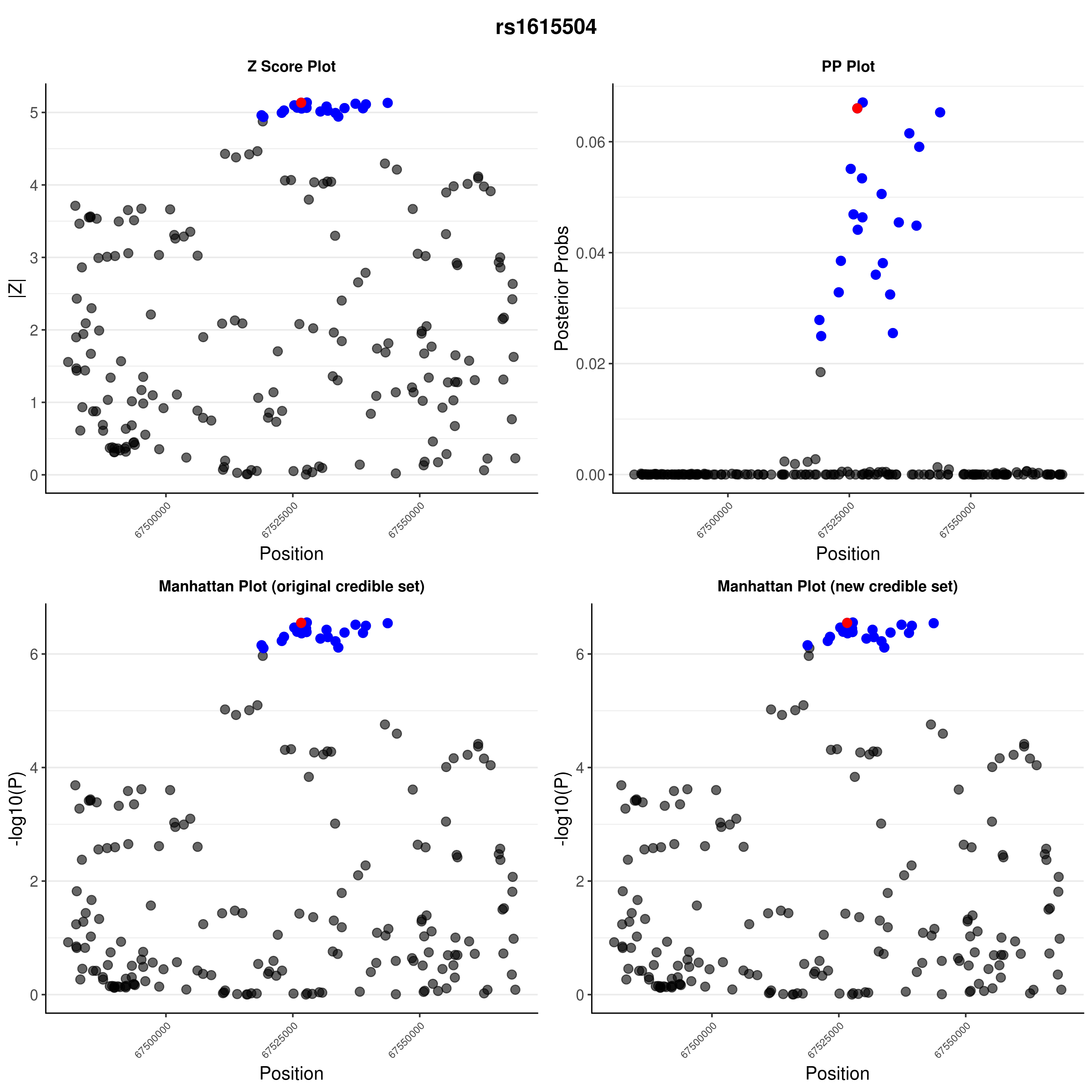

### rs1615504.png

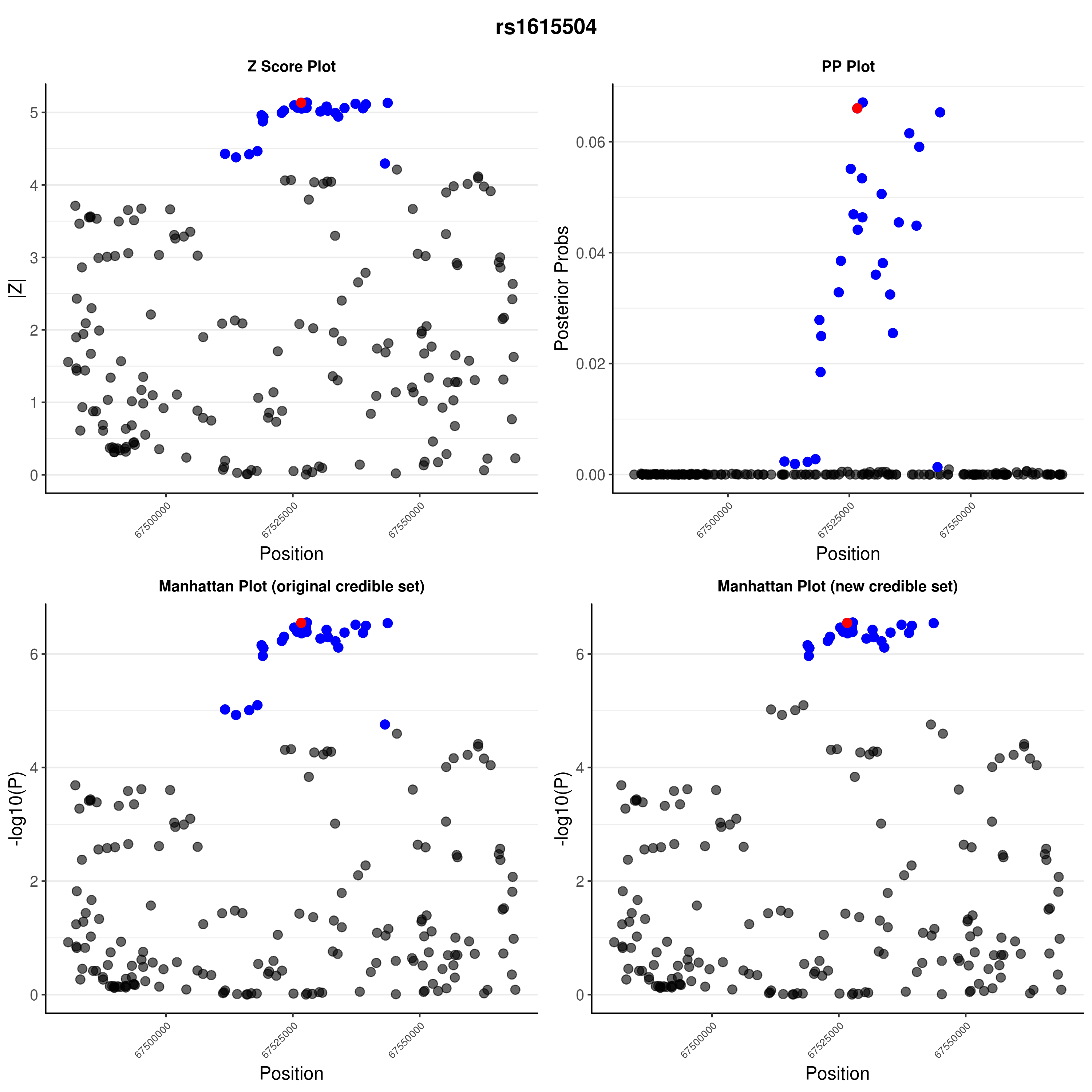

### rs1893217.png

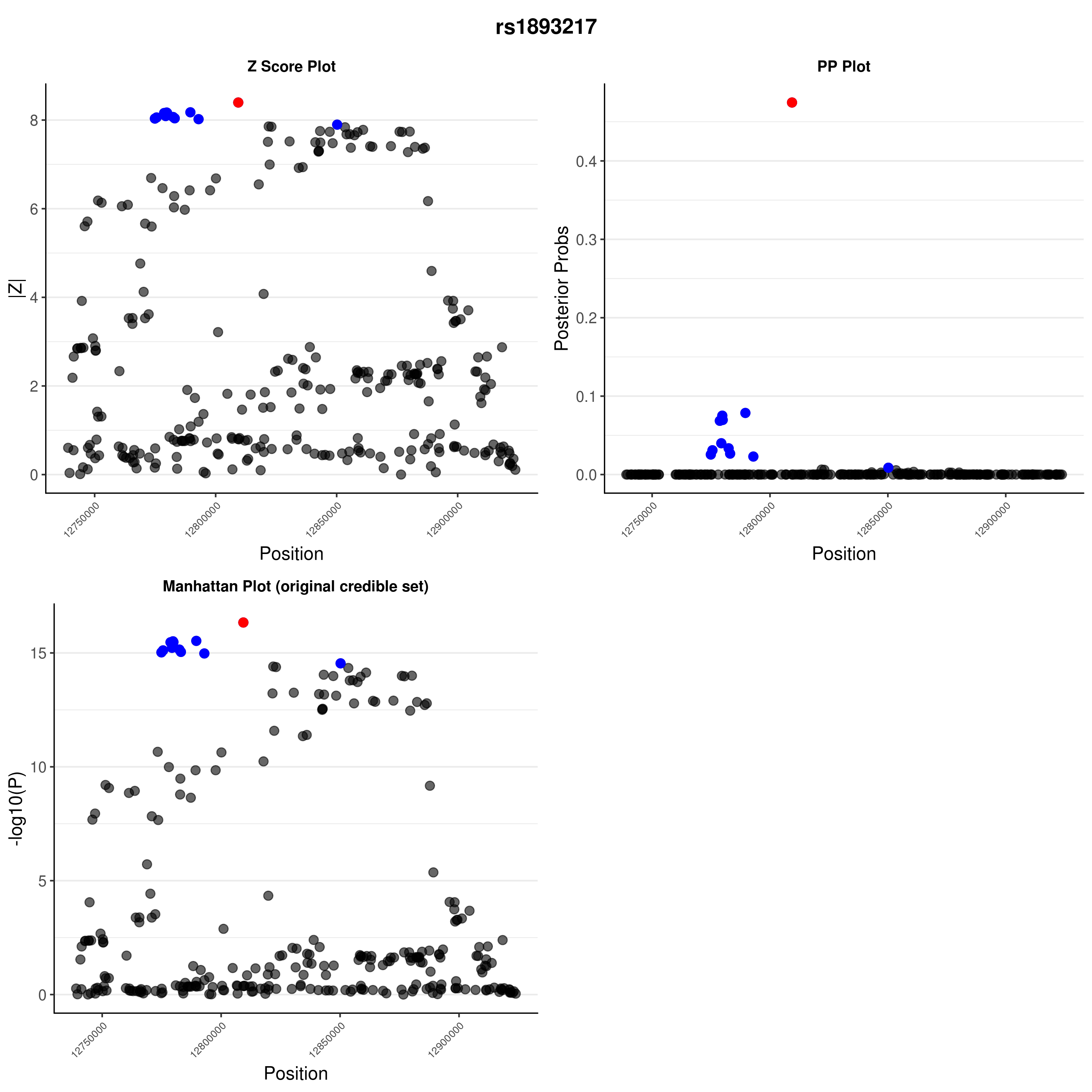

### rs1893217.png

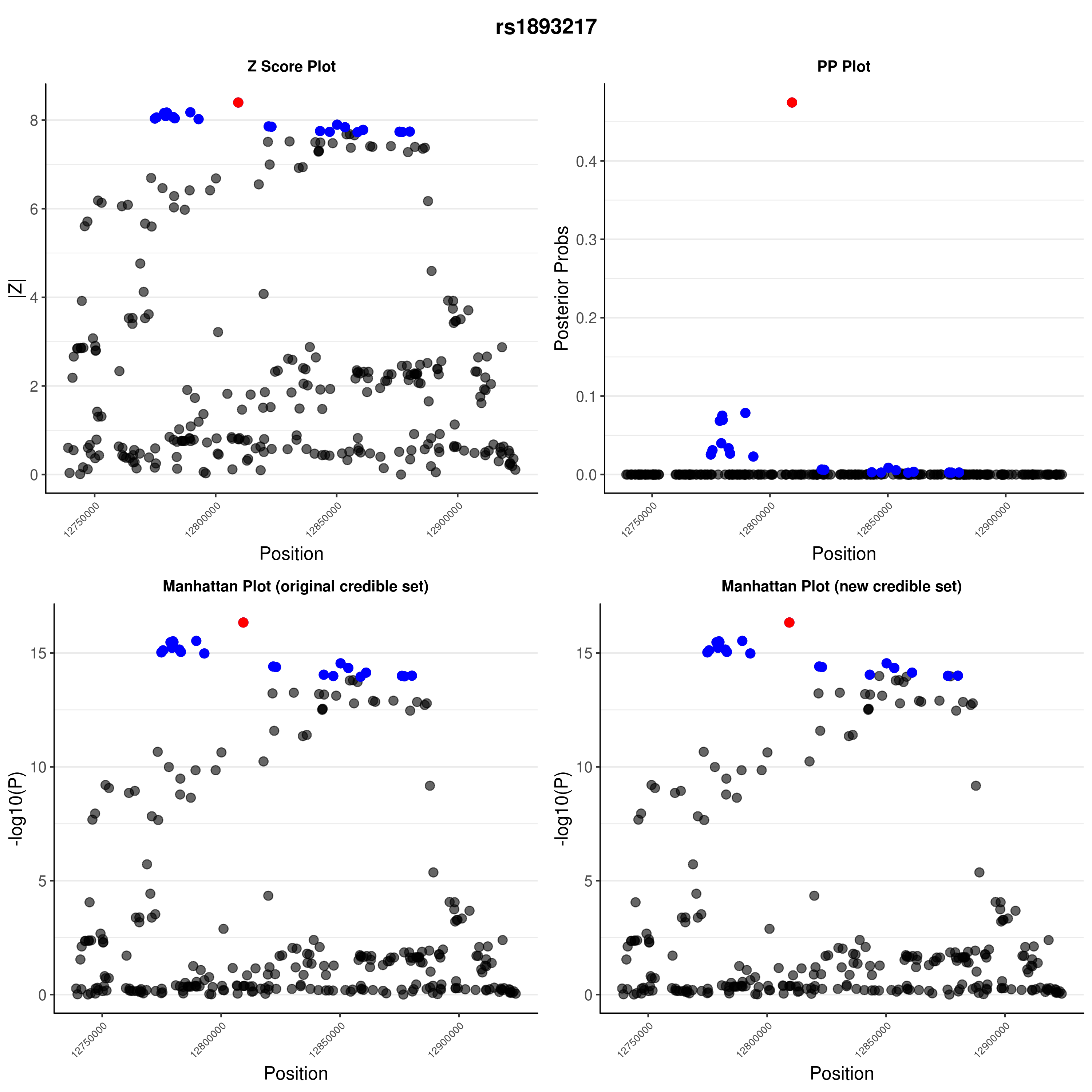

### rs2111485.png

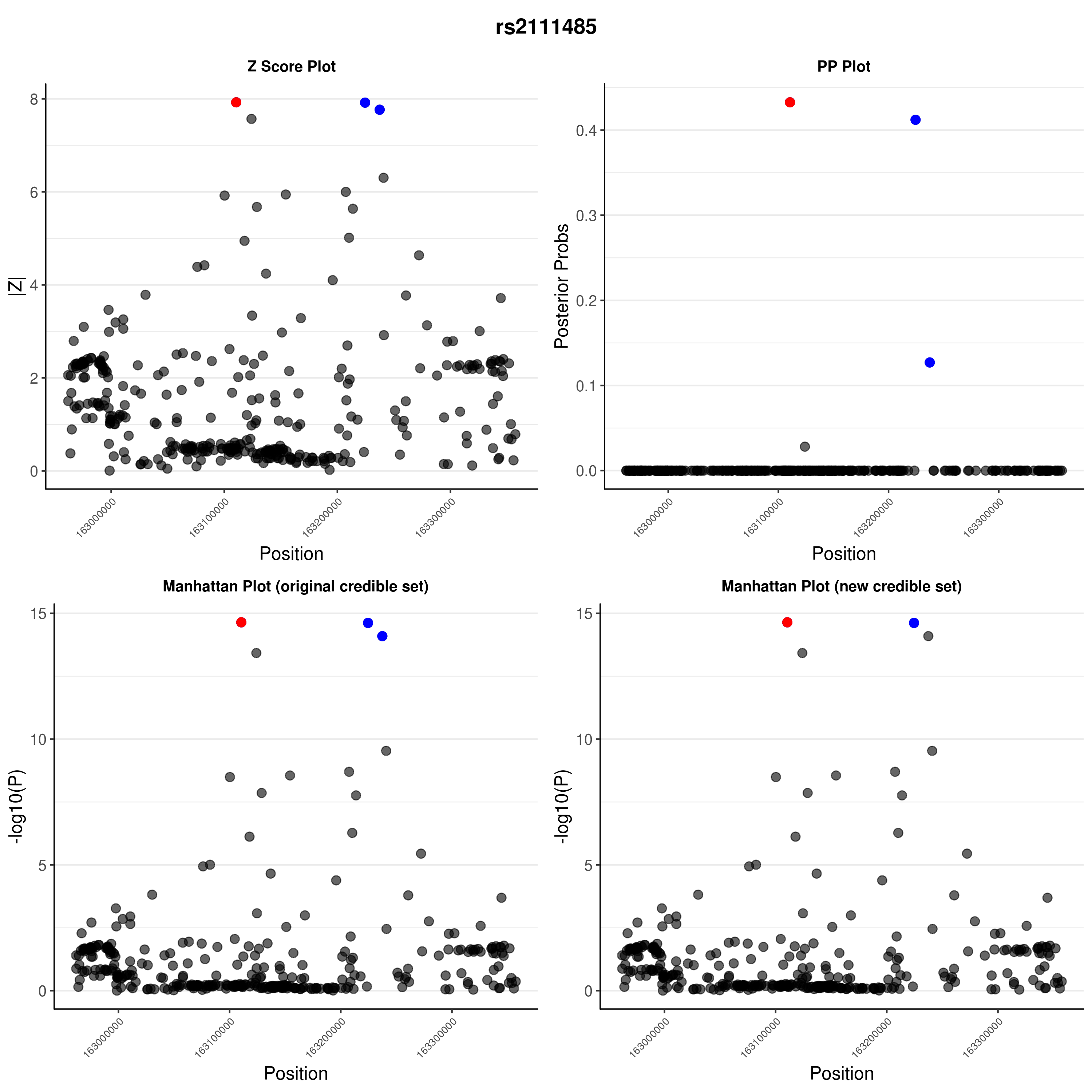

### rs2111485.png

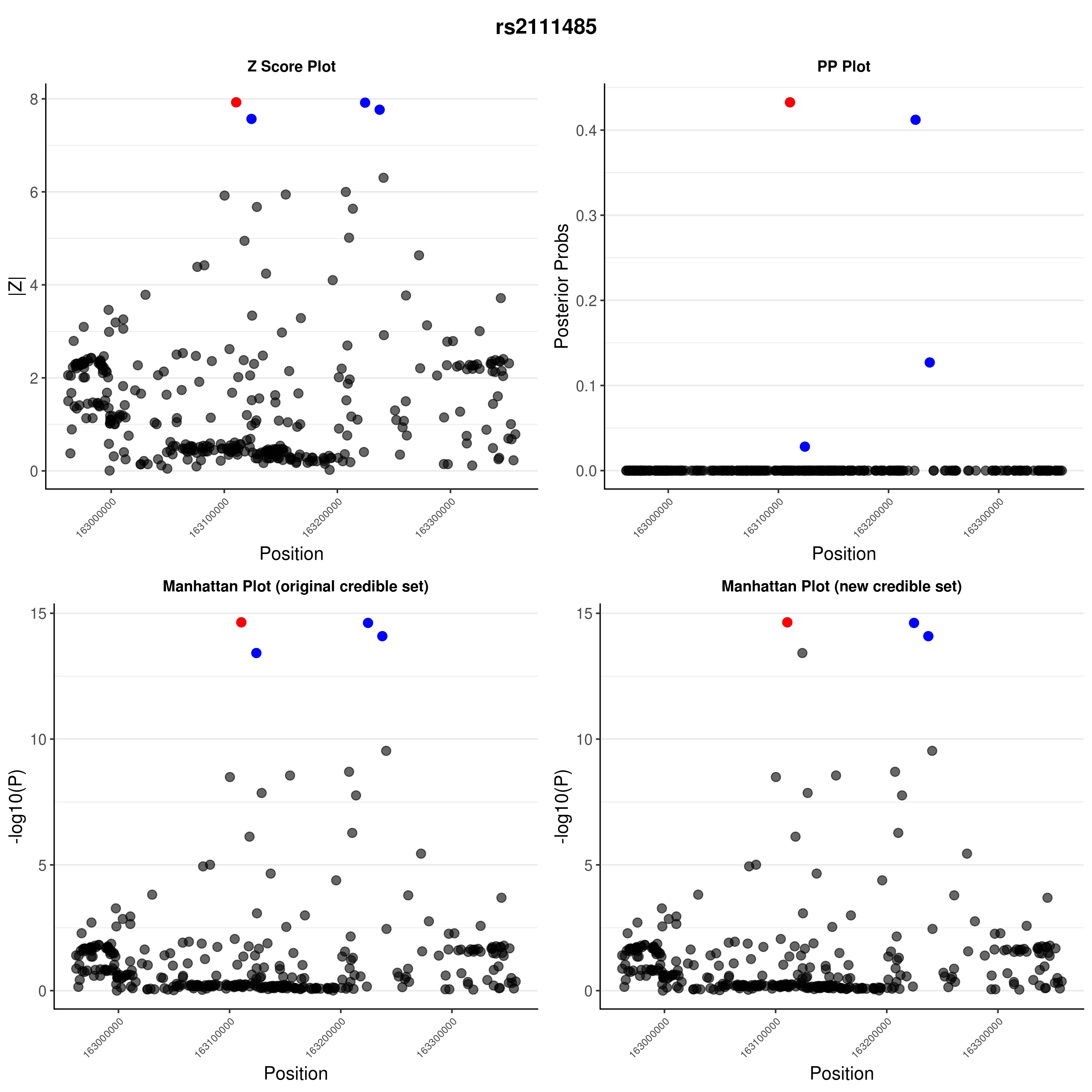

### rs2476601.png

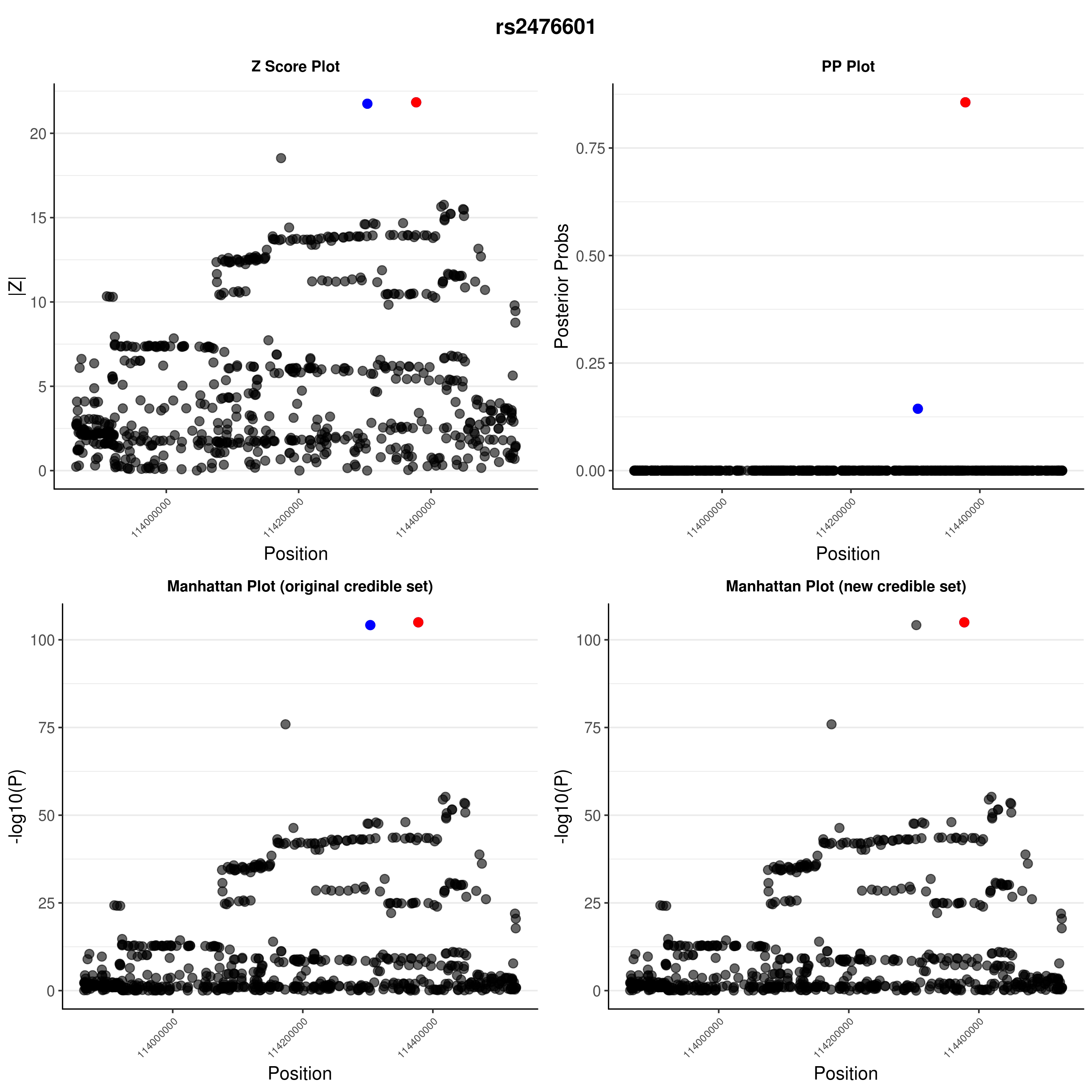

### rs2476601.png

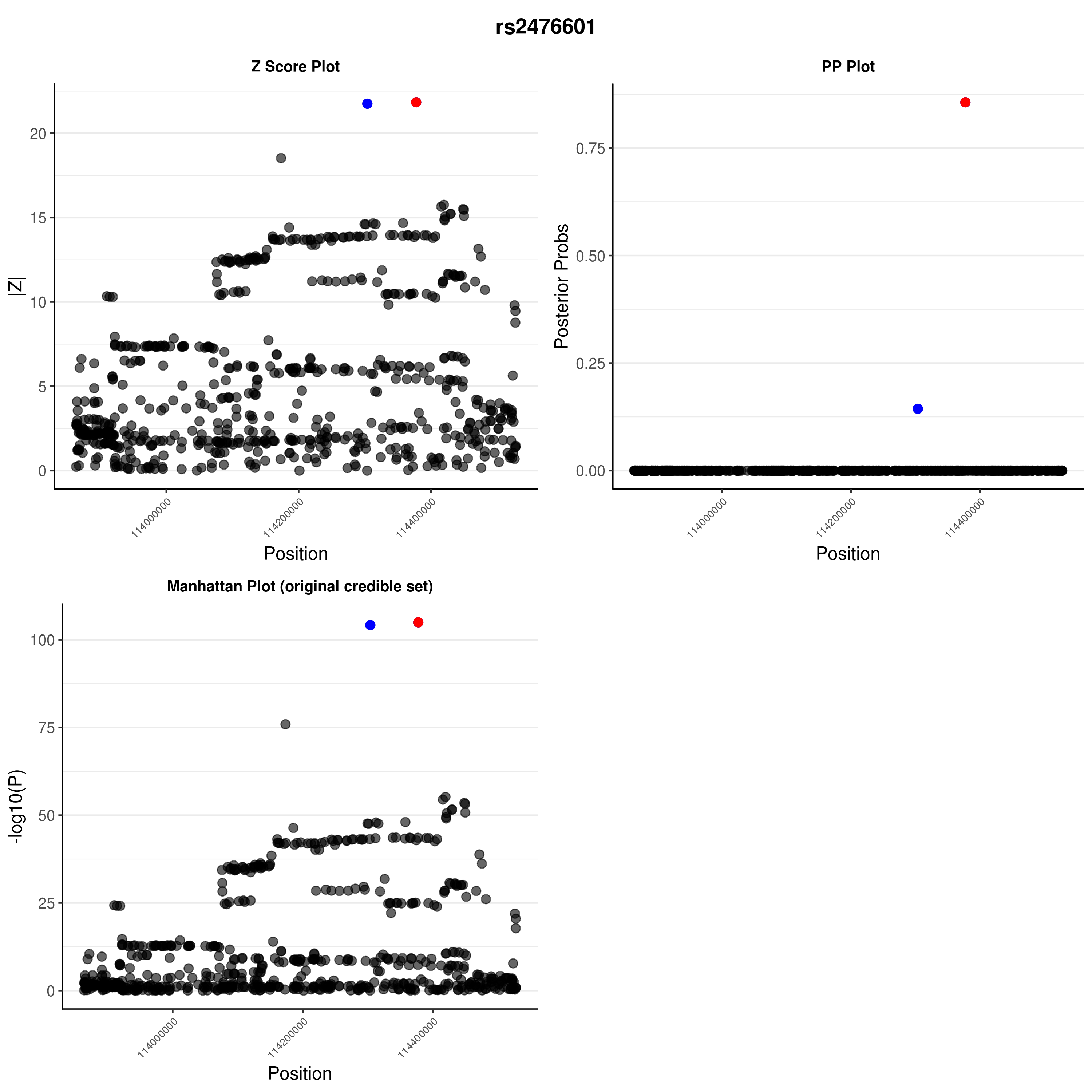

### rs3024505.png

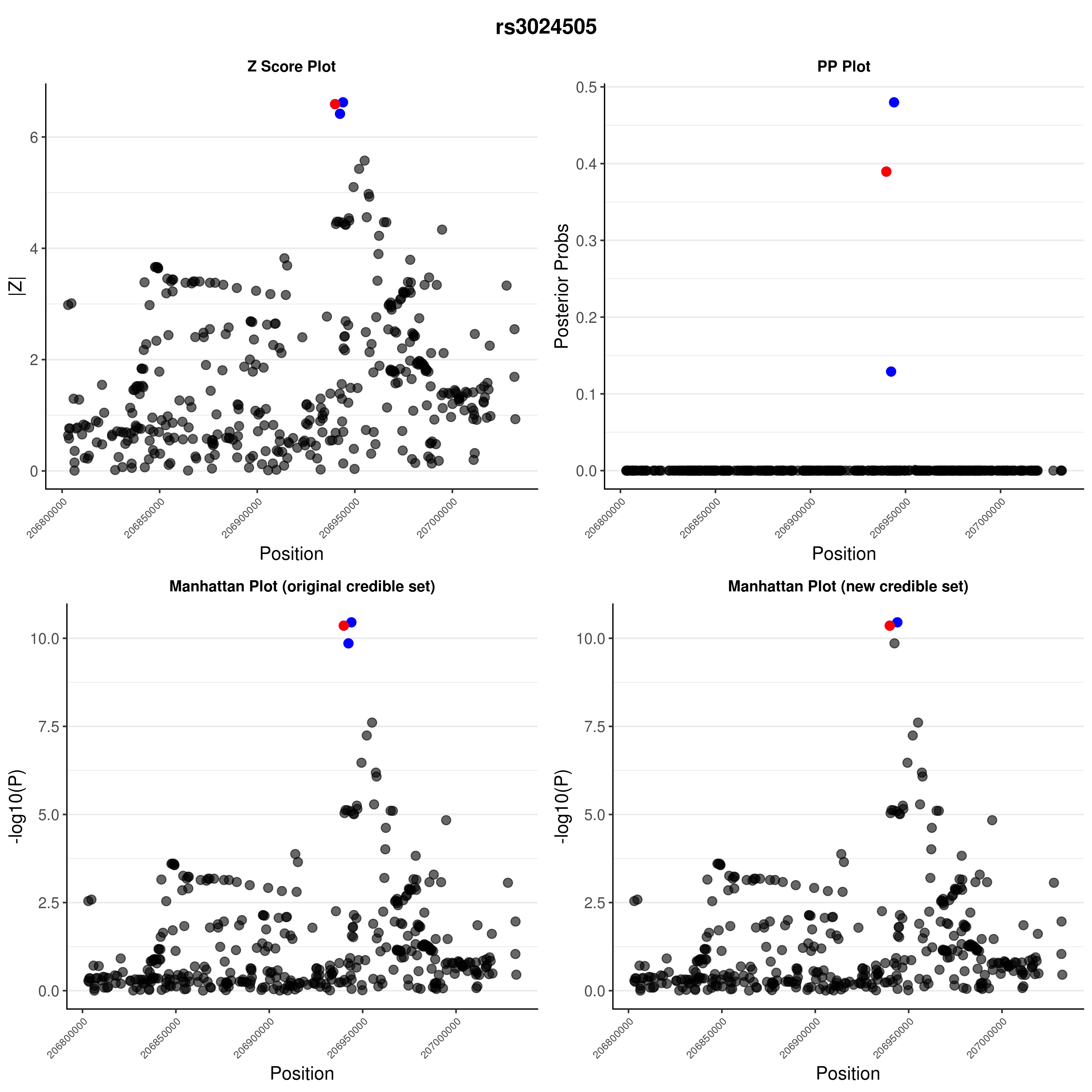

### rs3024505.png

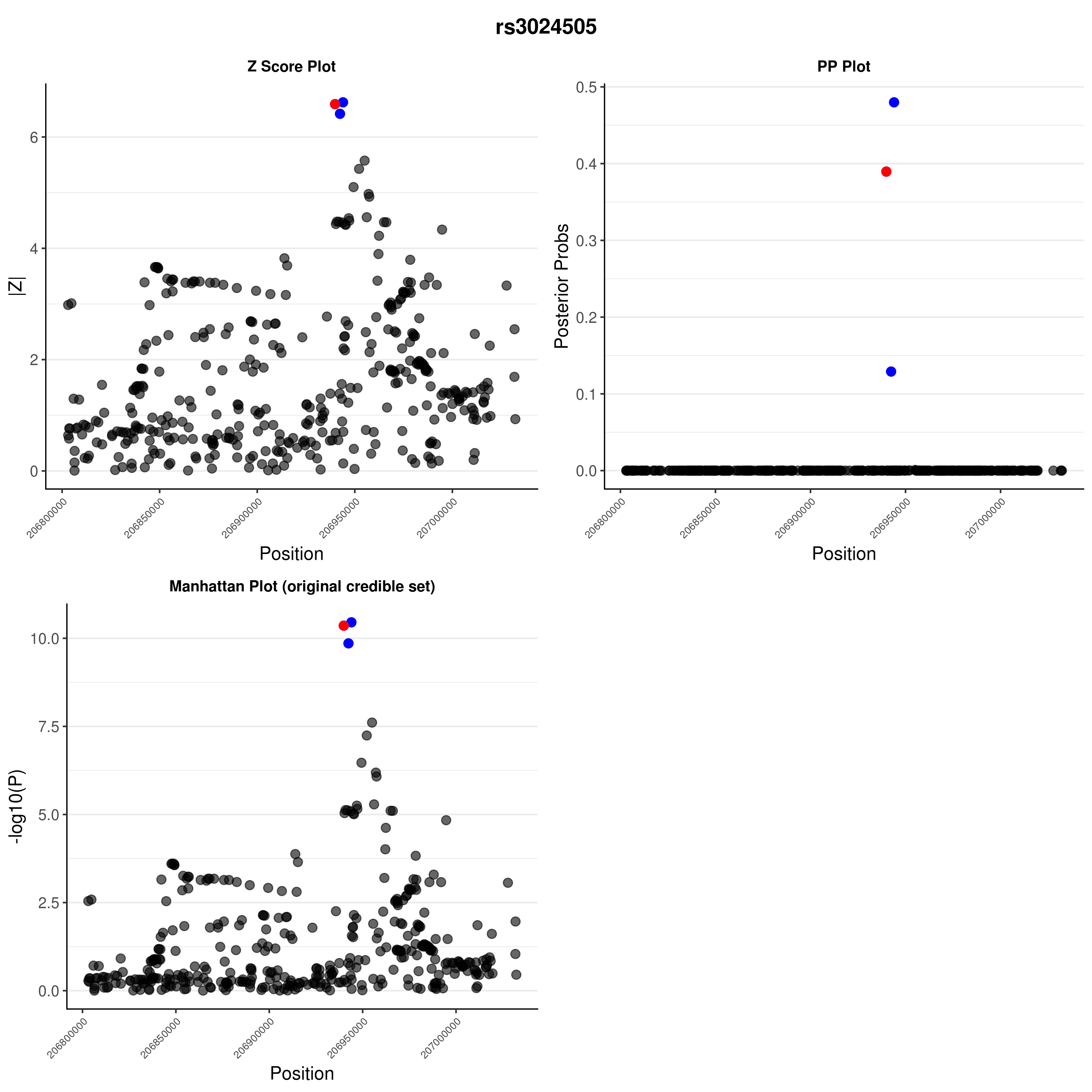

### rs3087243.png

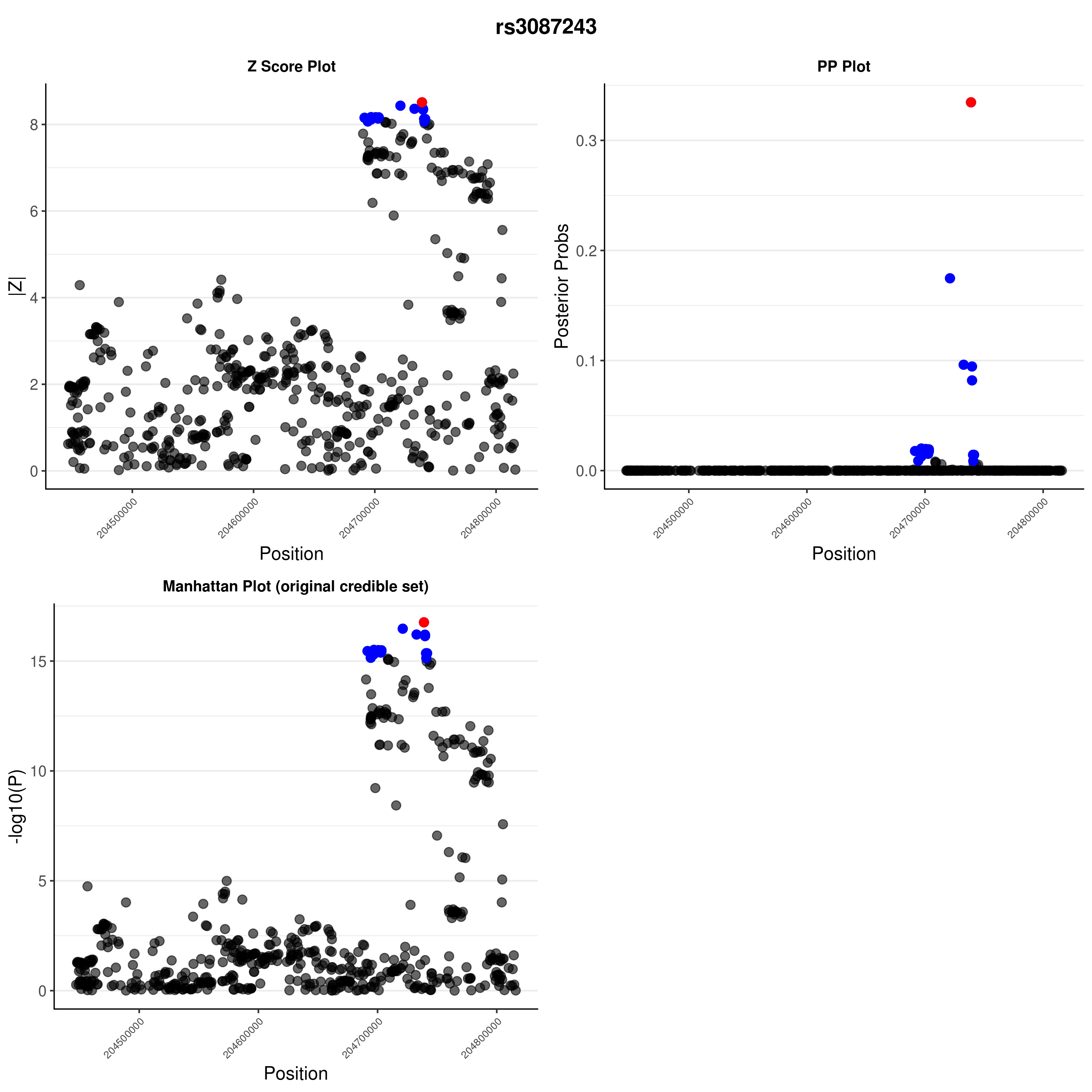

### rs3087243.png

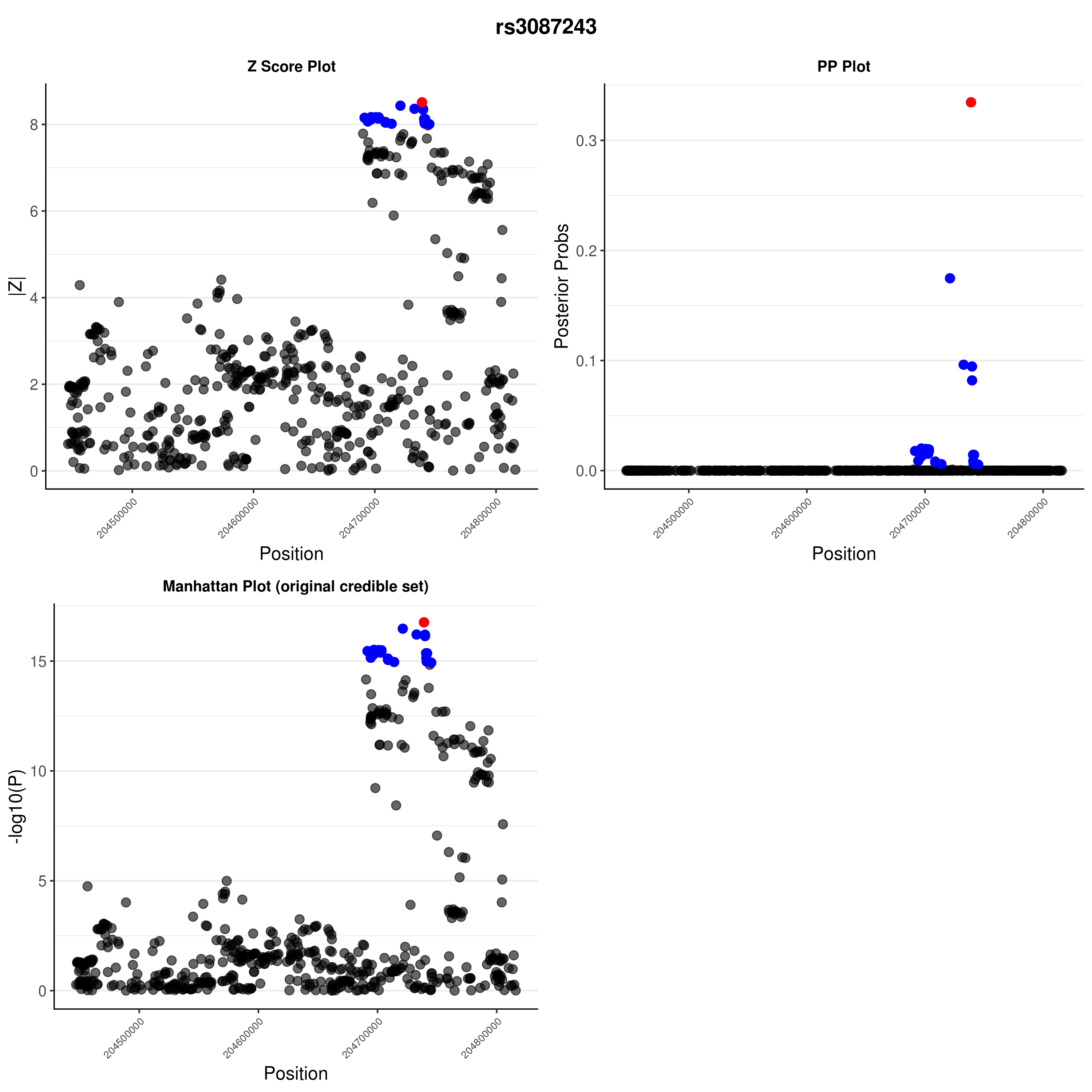

### rs4820830.png

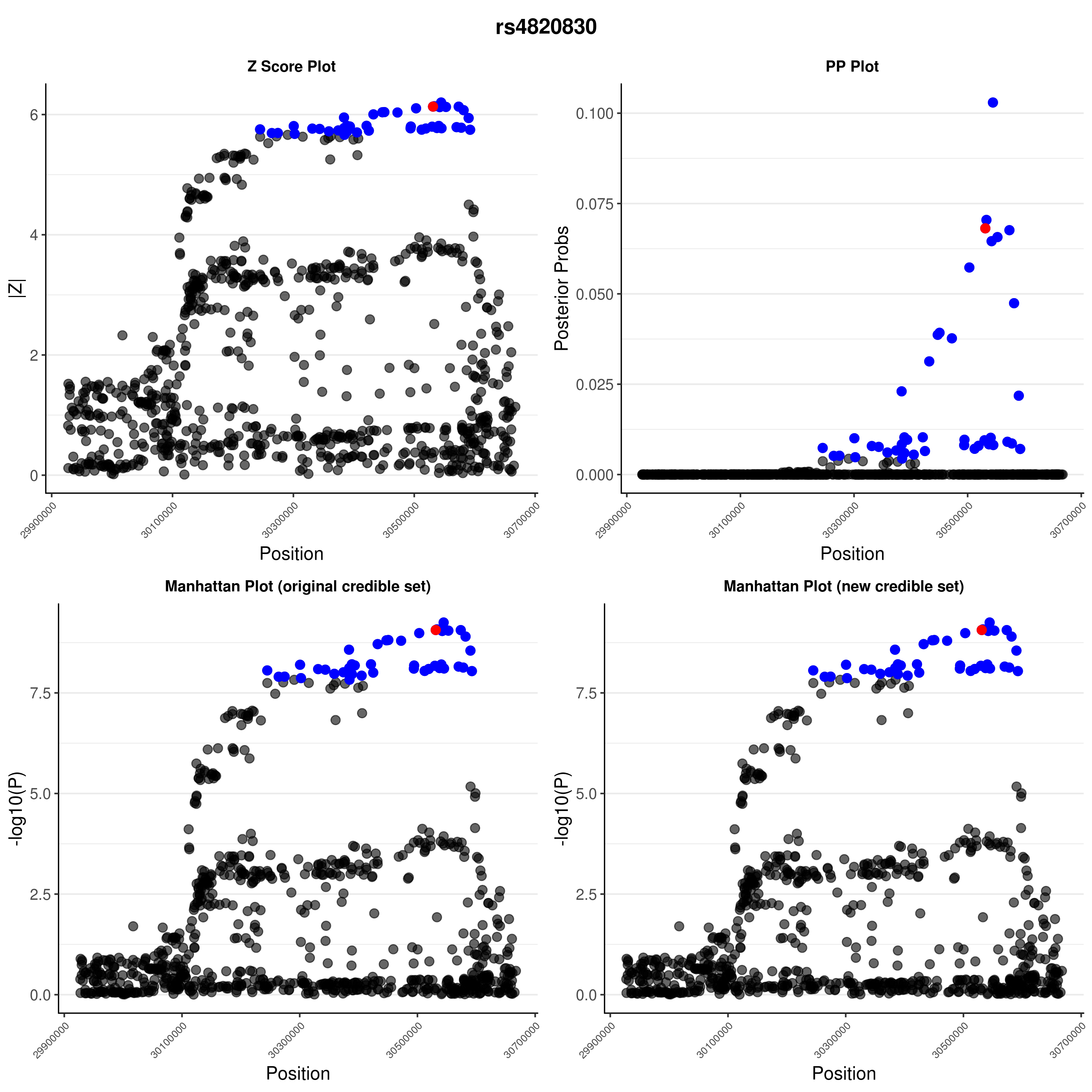

### rs4820830.png

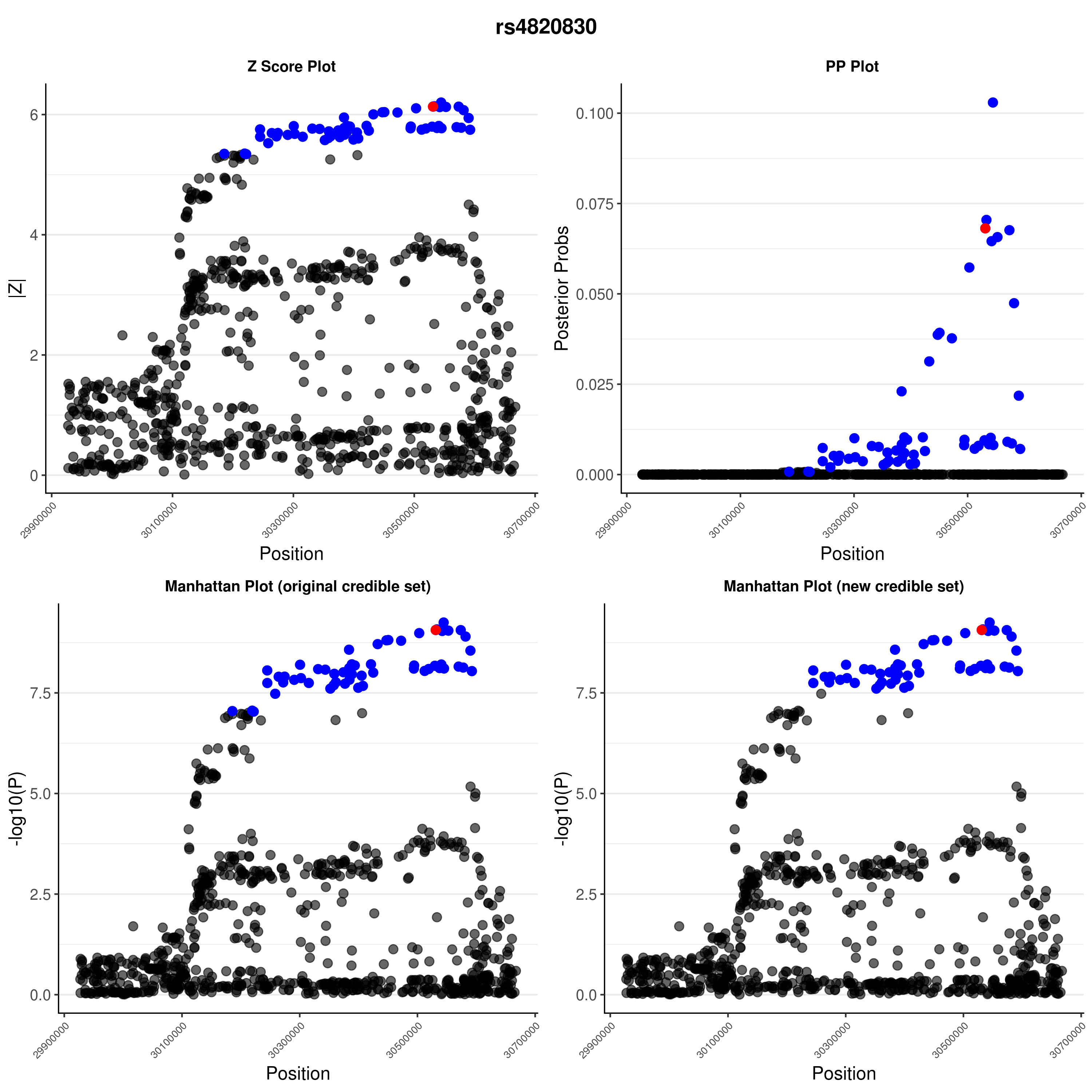

### rs6043409.png

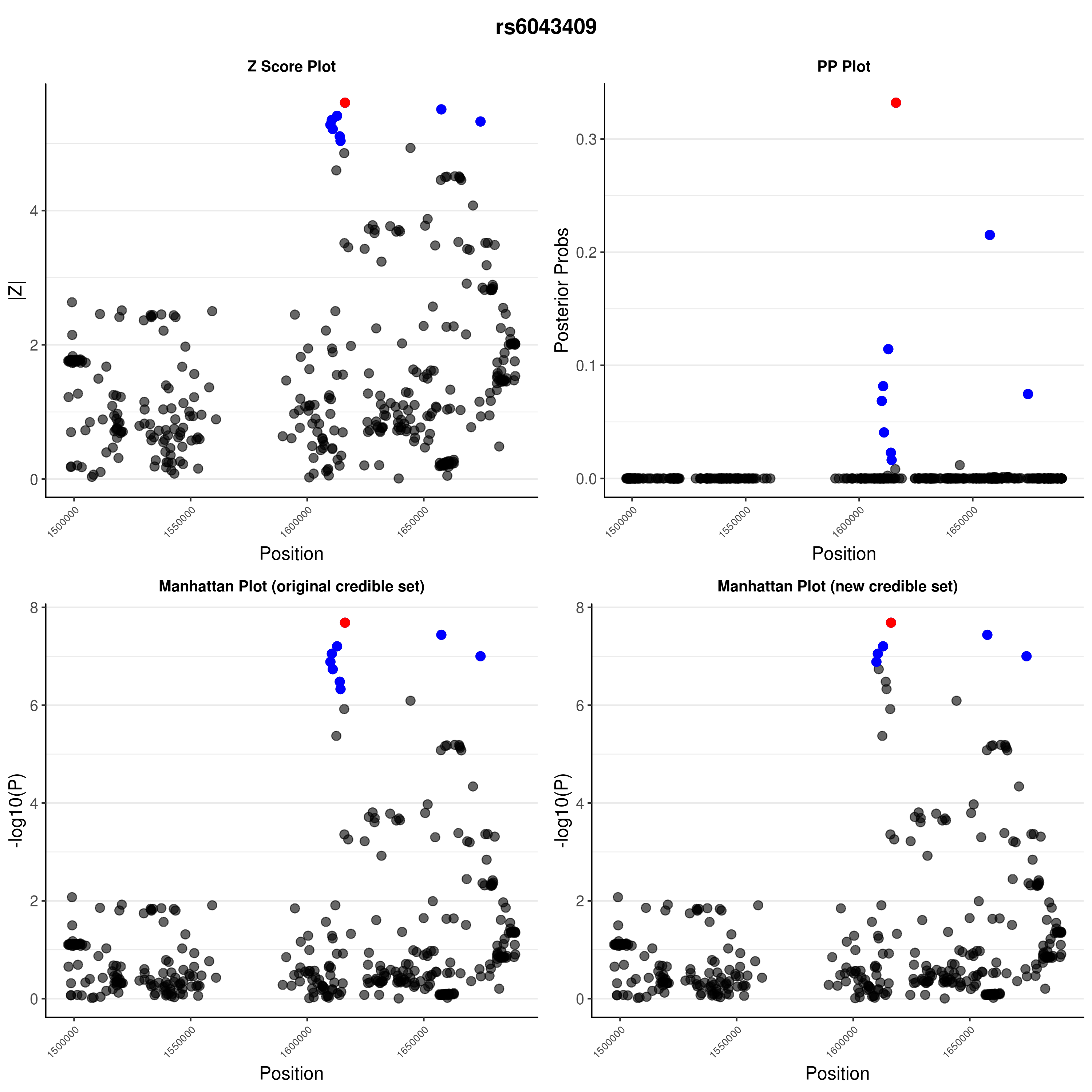

### rs6043409.png

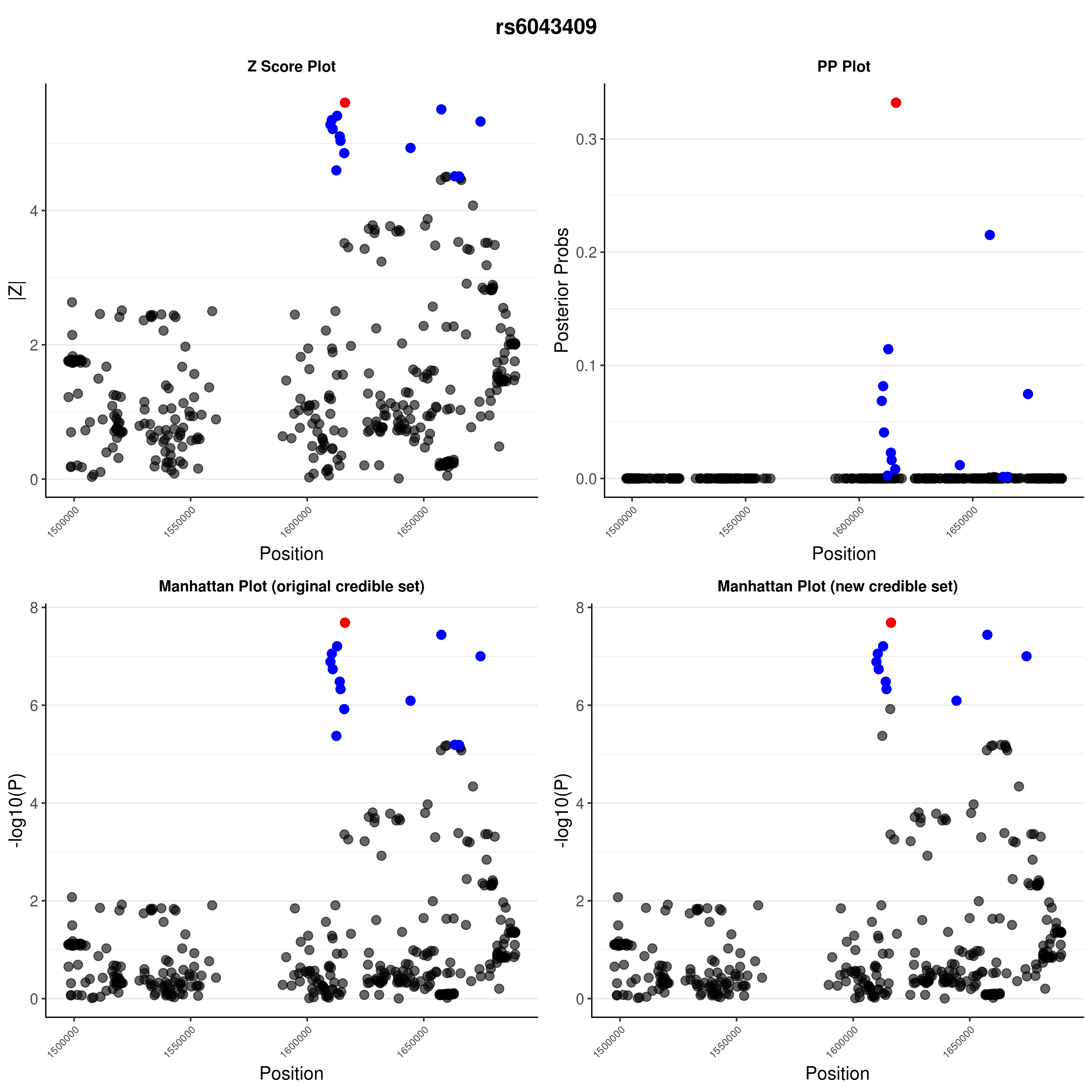

### rs6476839.png

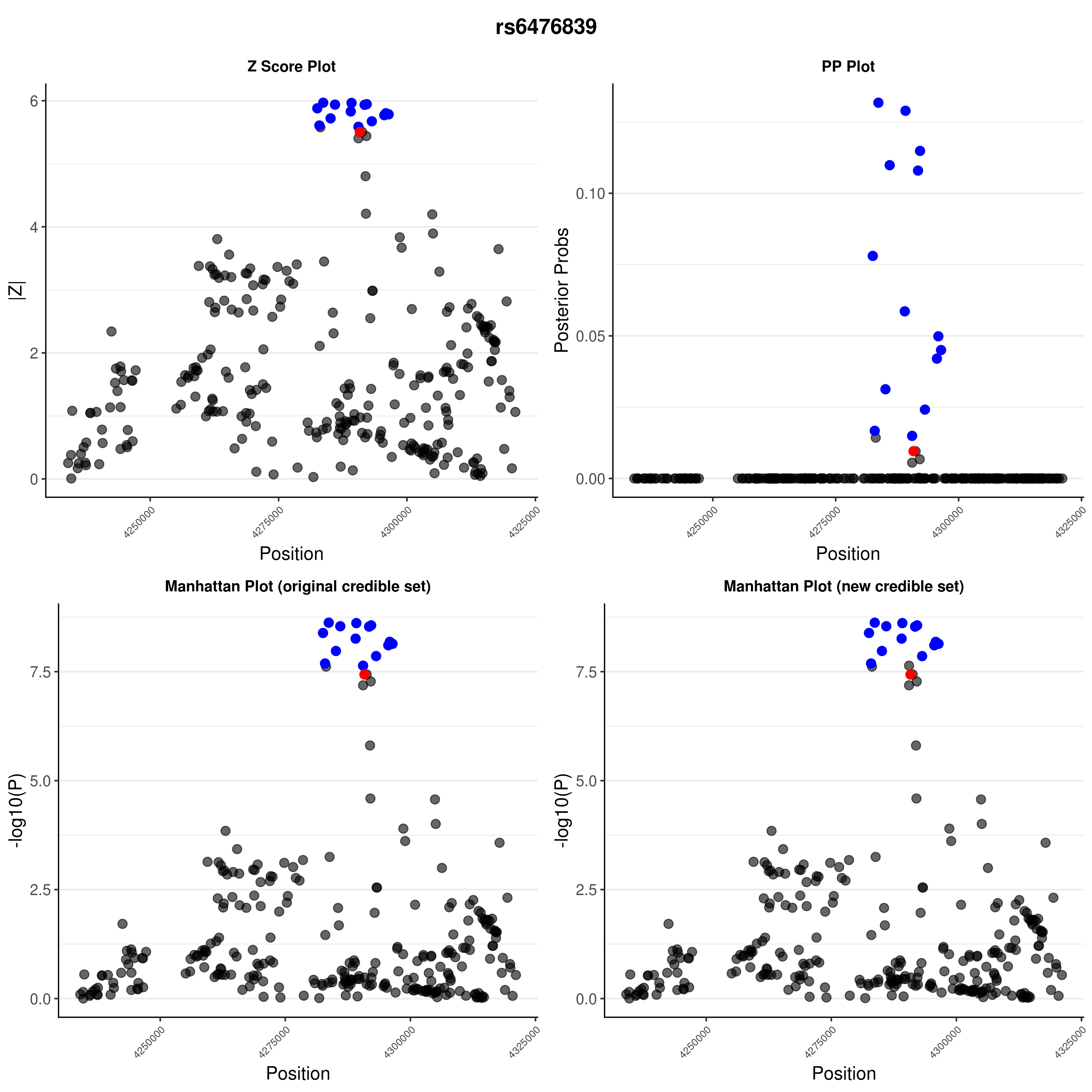

### rs6476839.png

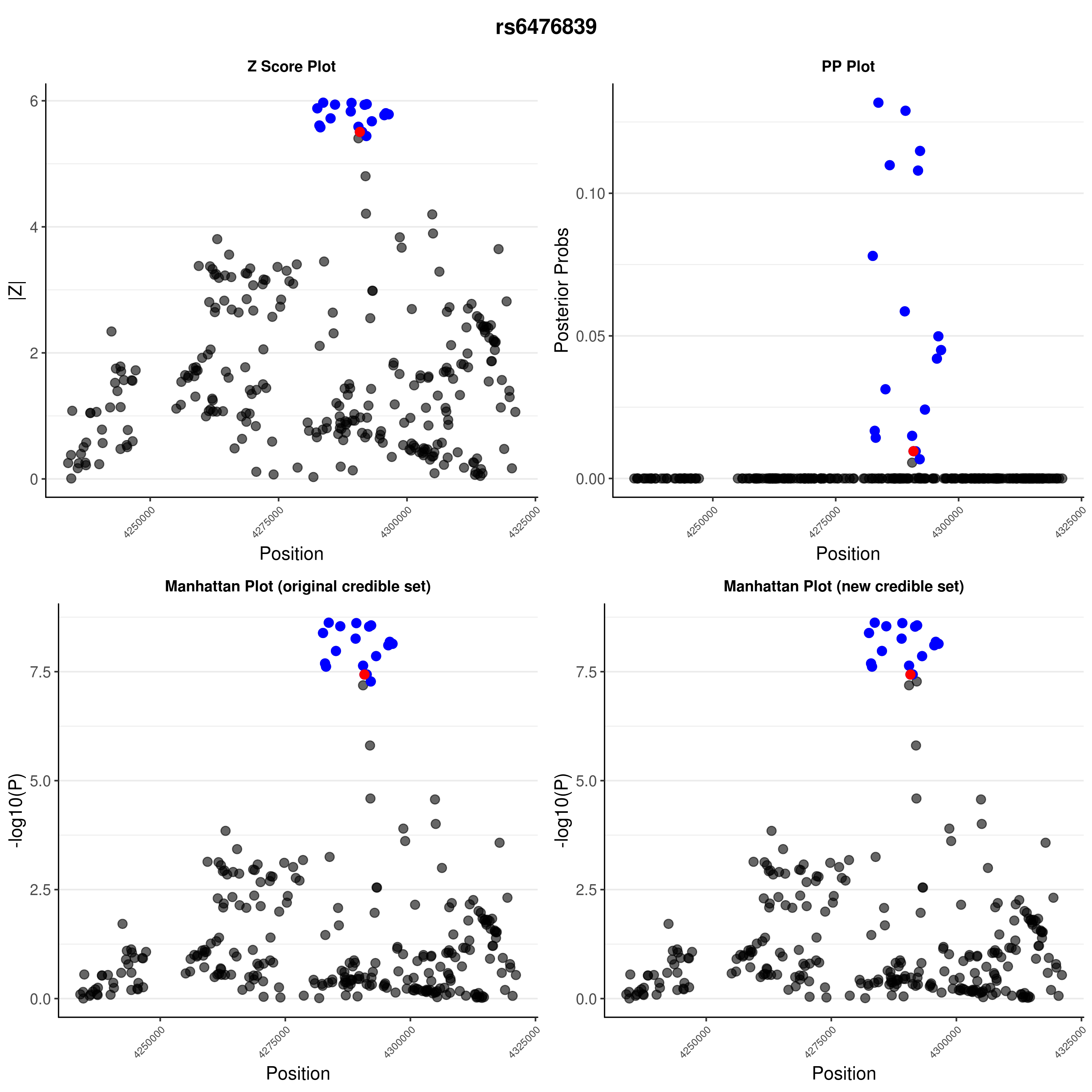

### rs6518350.png

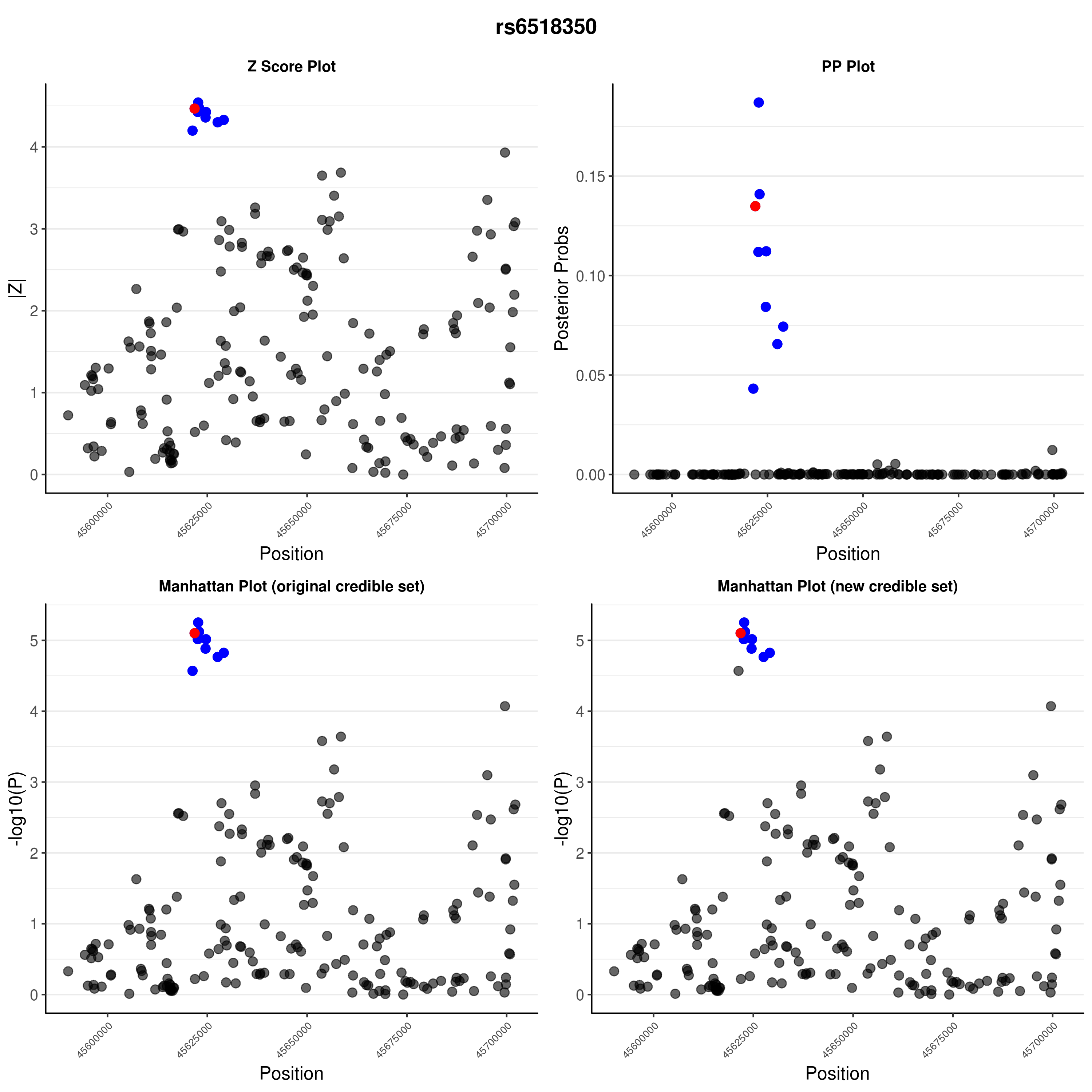

### rs6518350.png

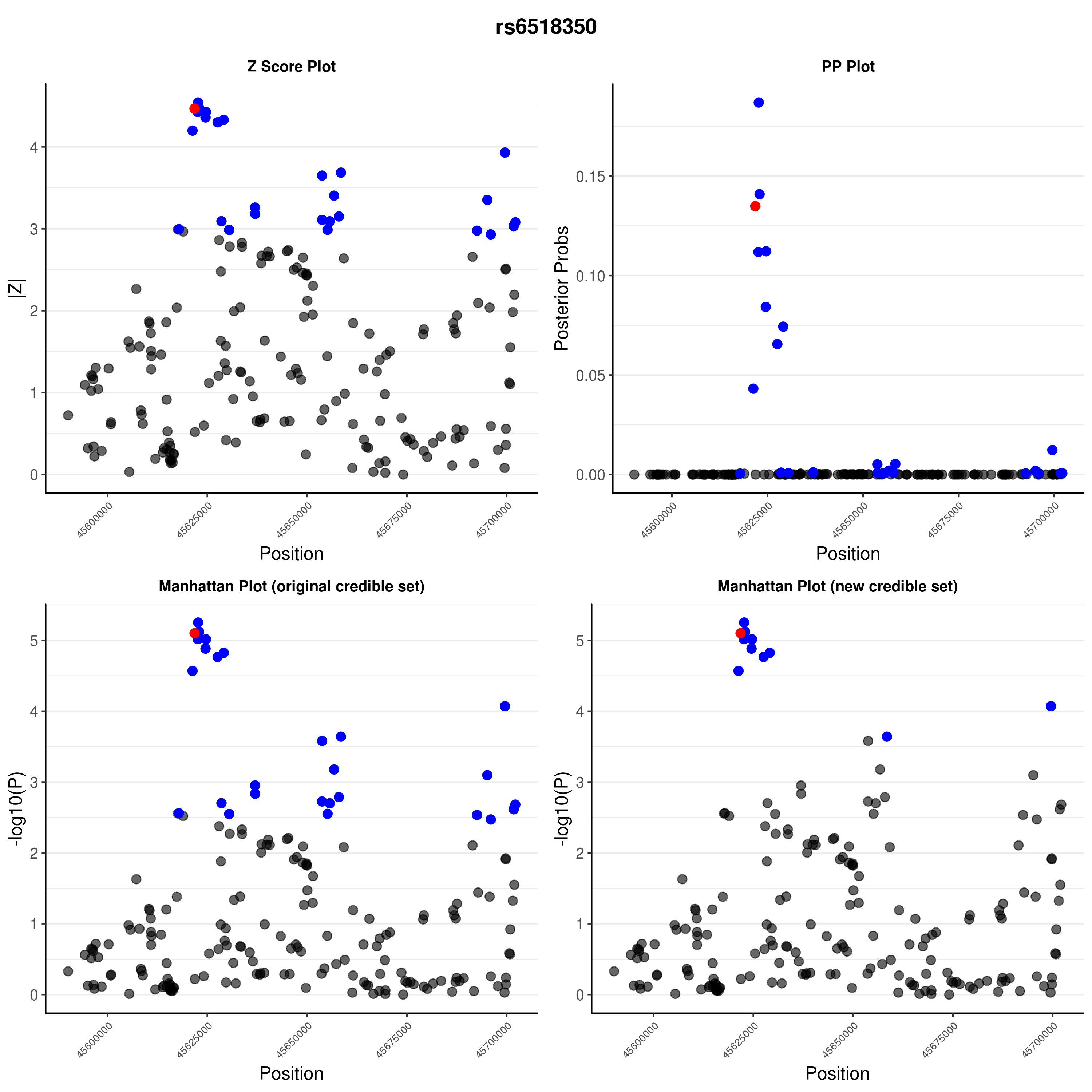

### rs8056814.png

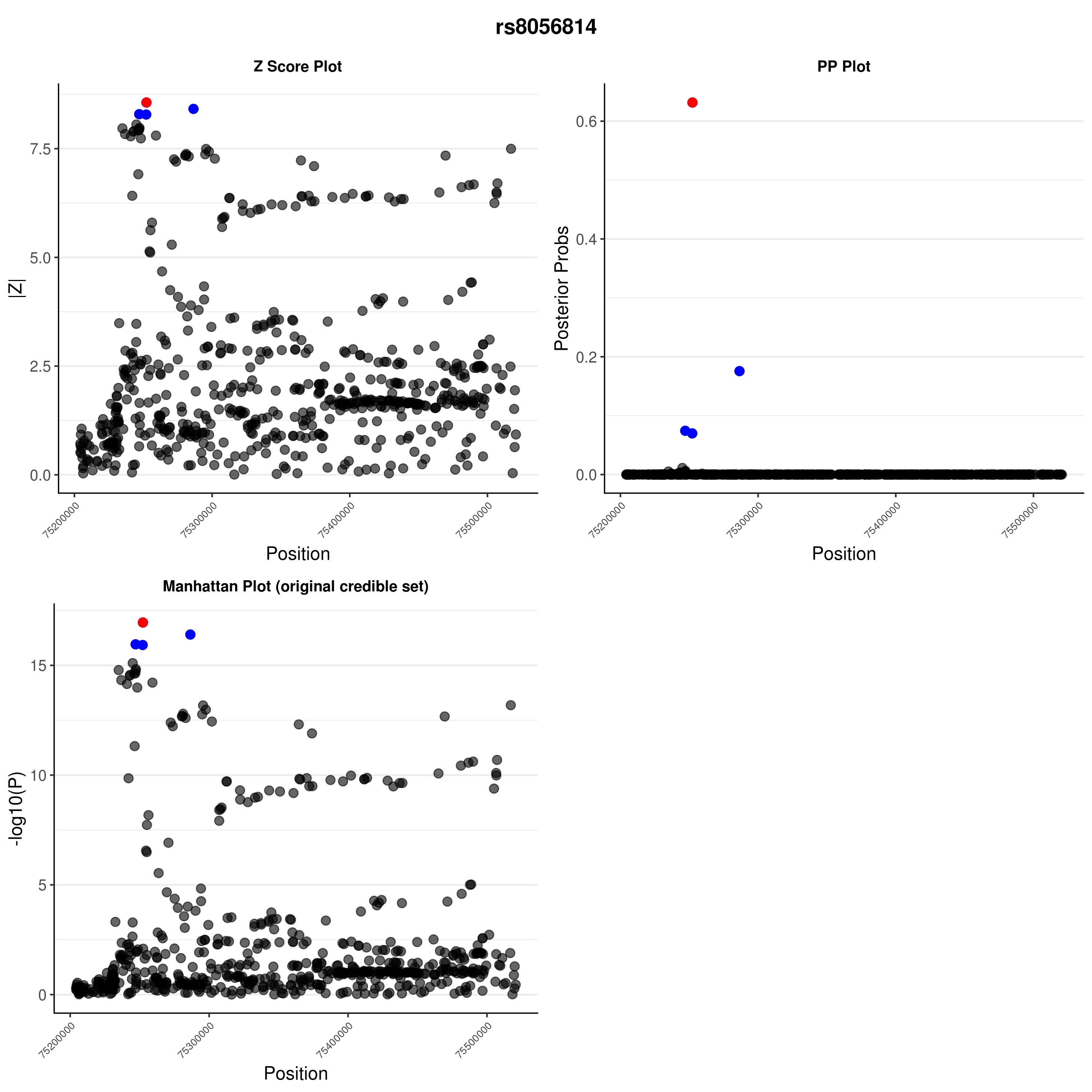

### rs8056814.png

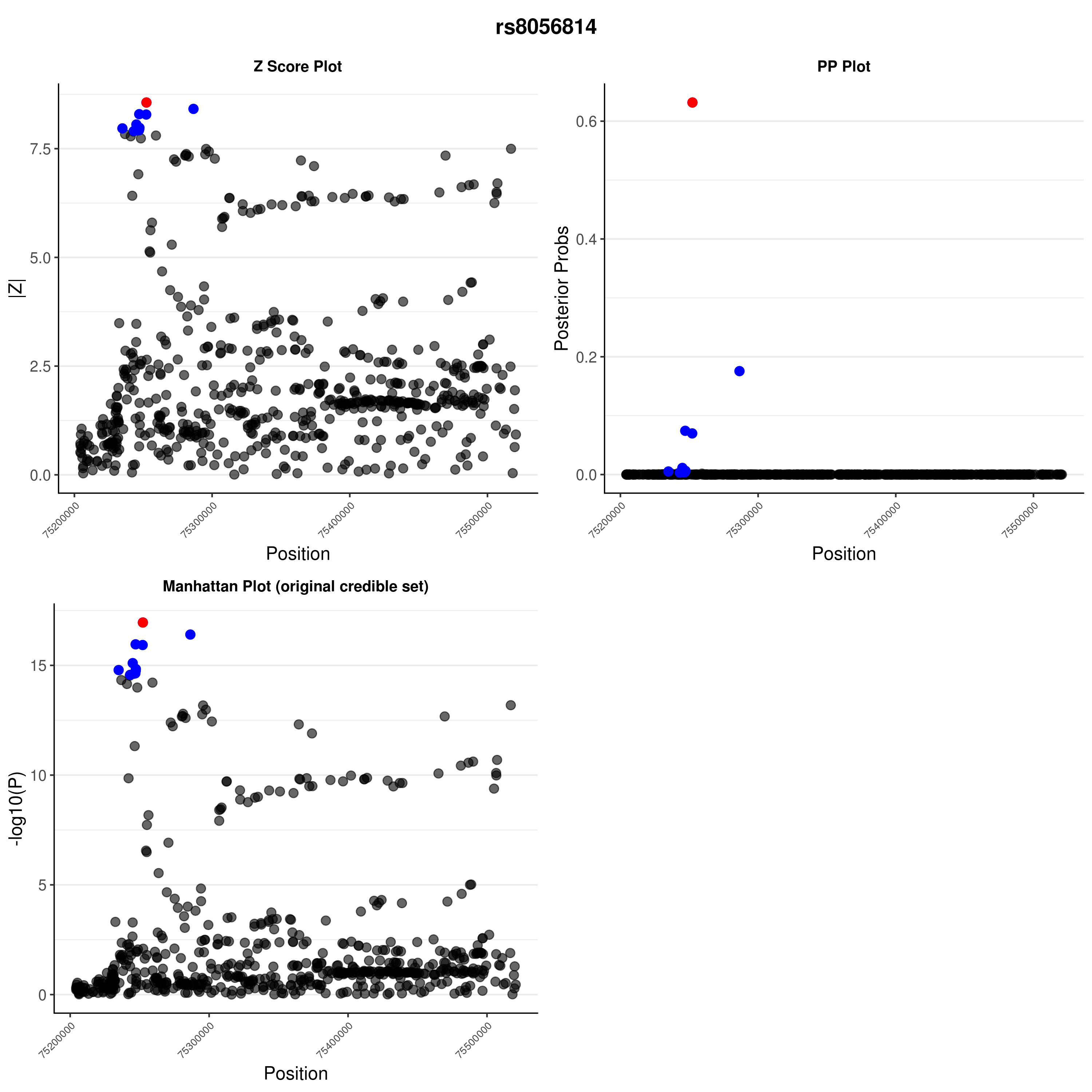

### rs9585056.png

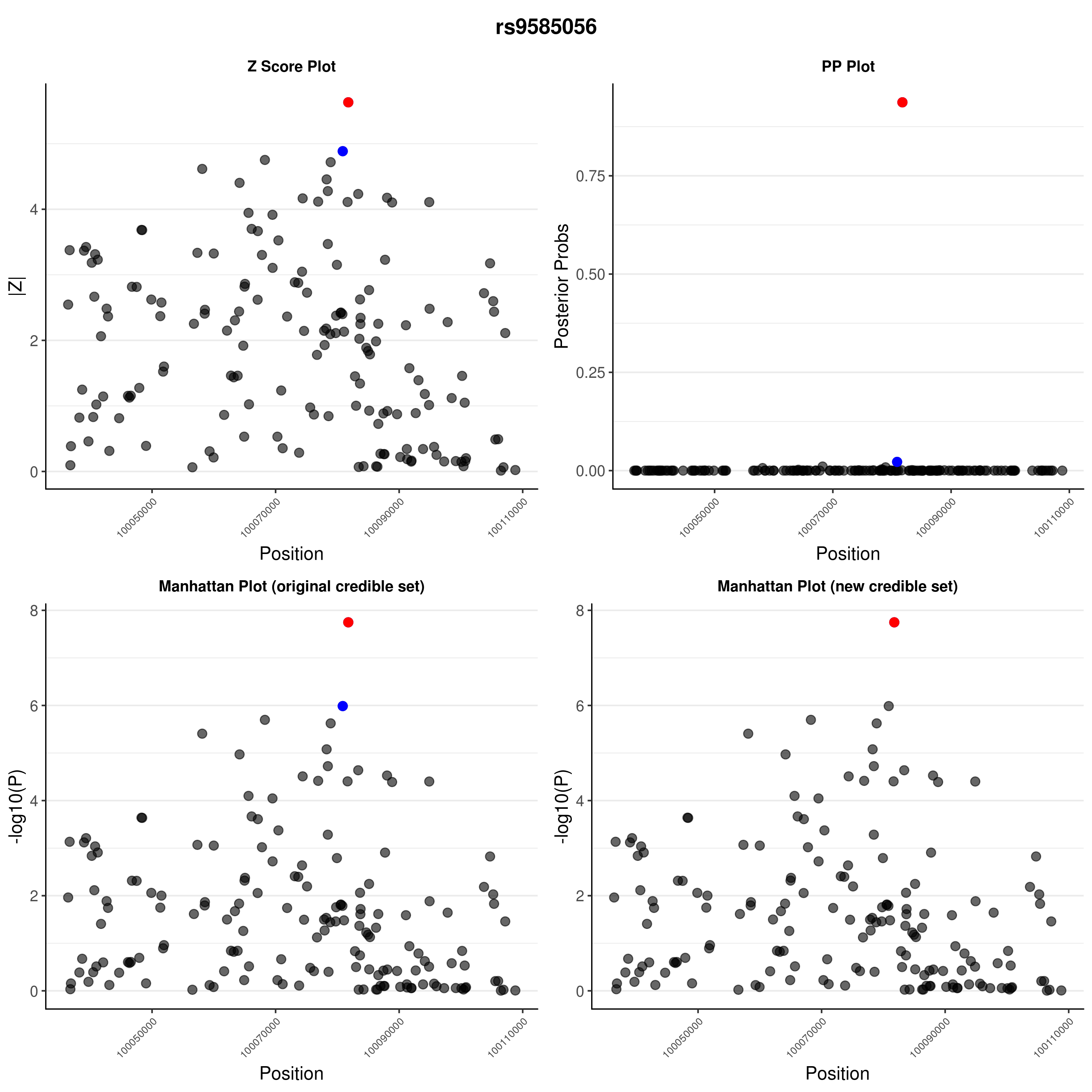

### rs9585056.png

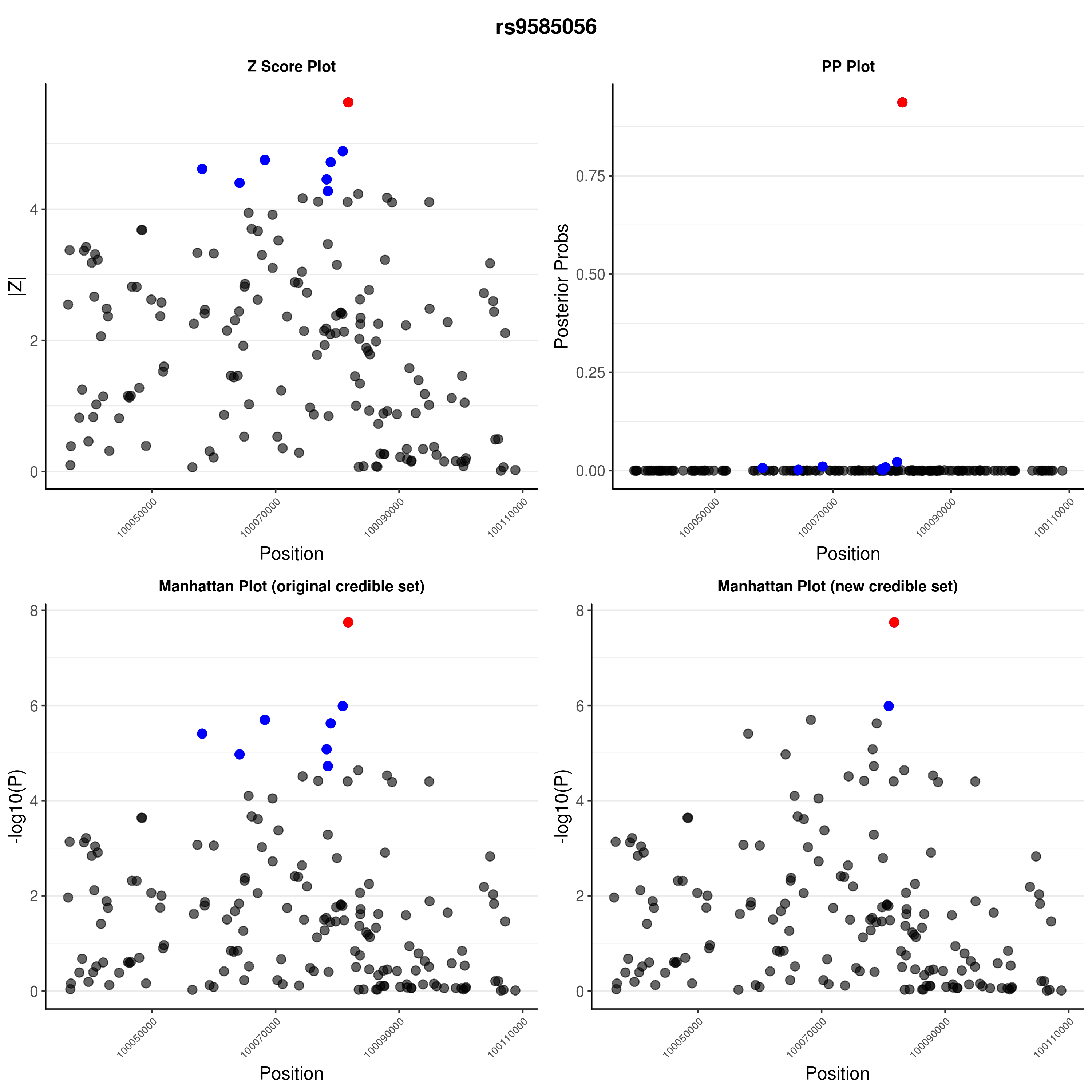

### rs10277986.png

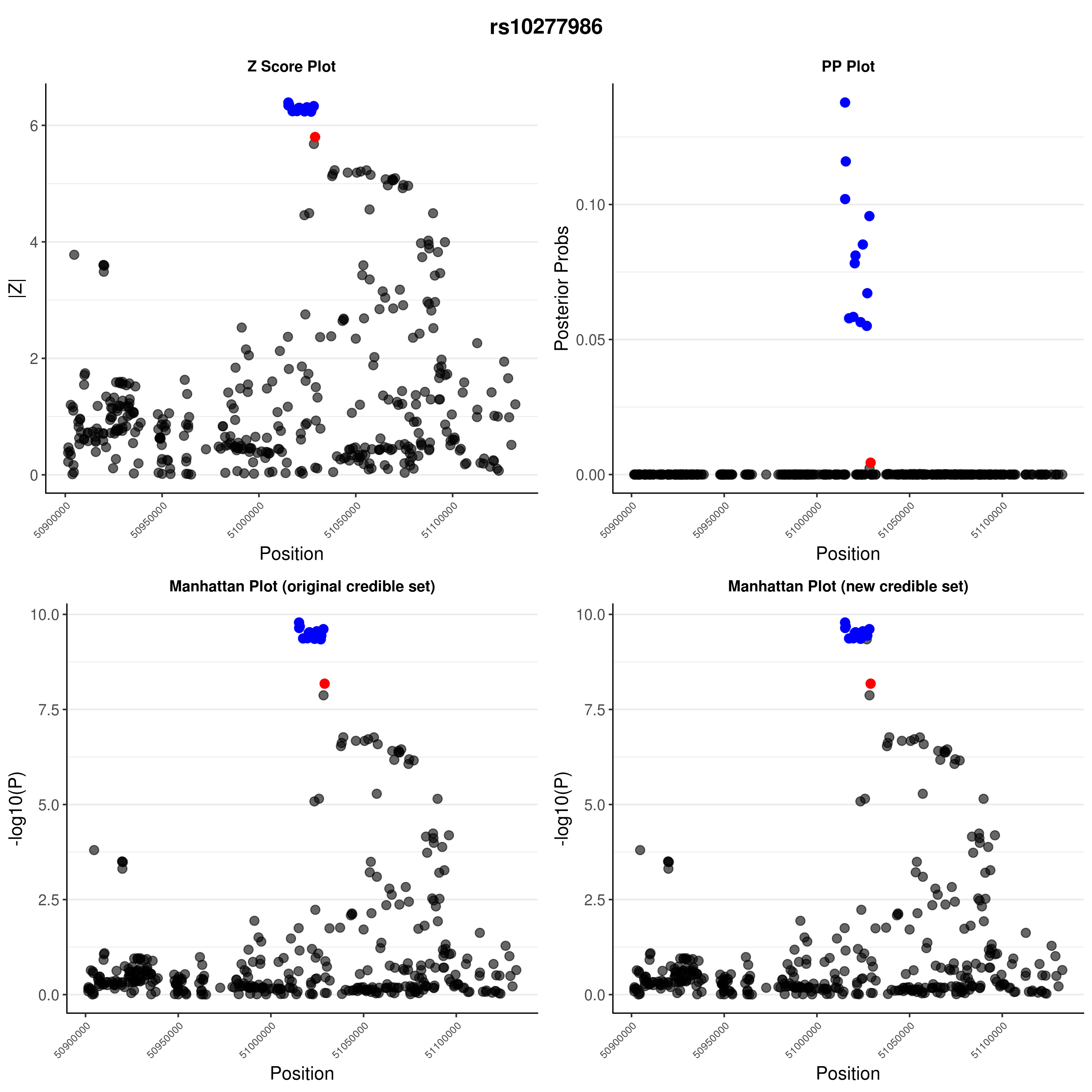

### rs10277986.png

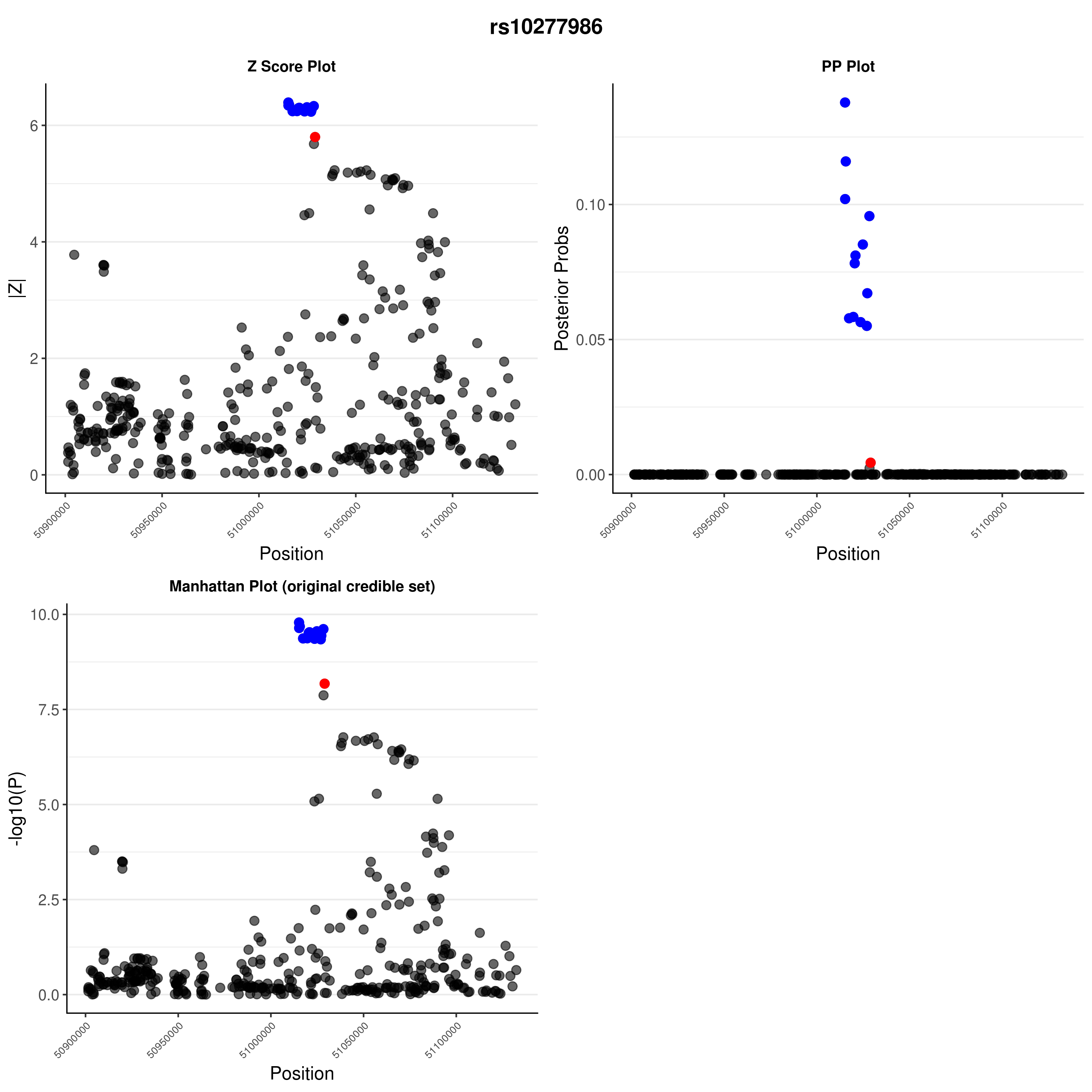

### rs11203202.png

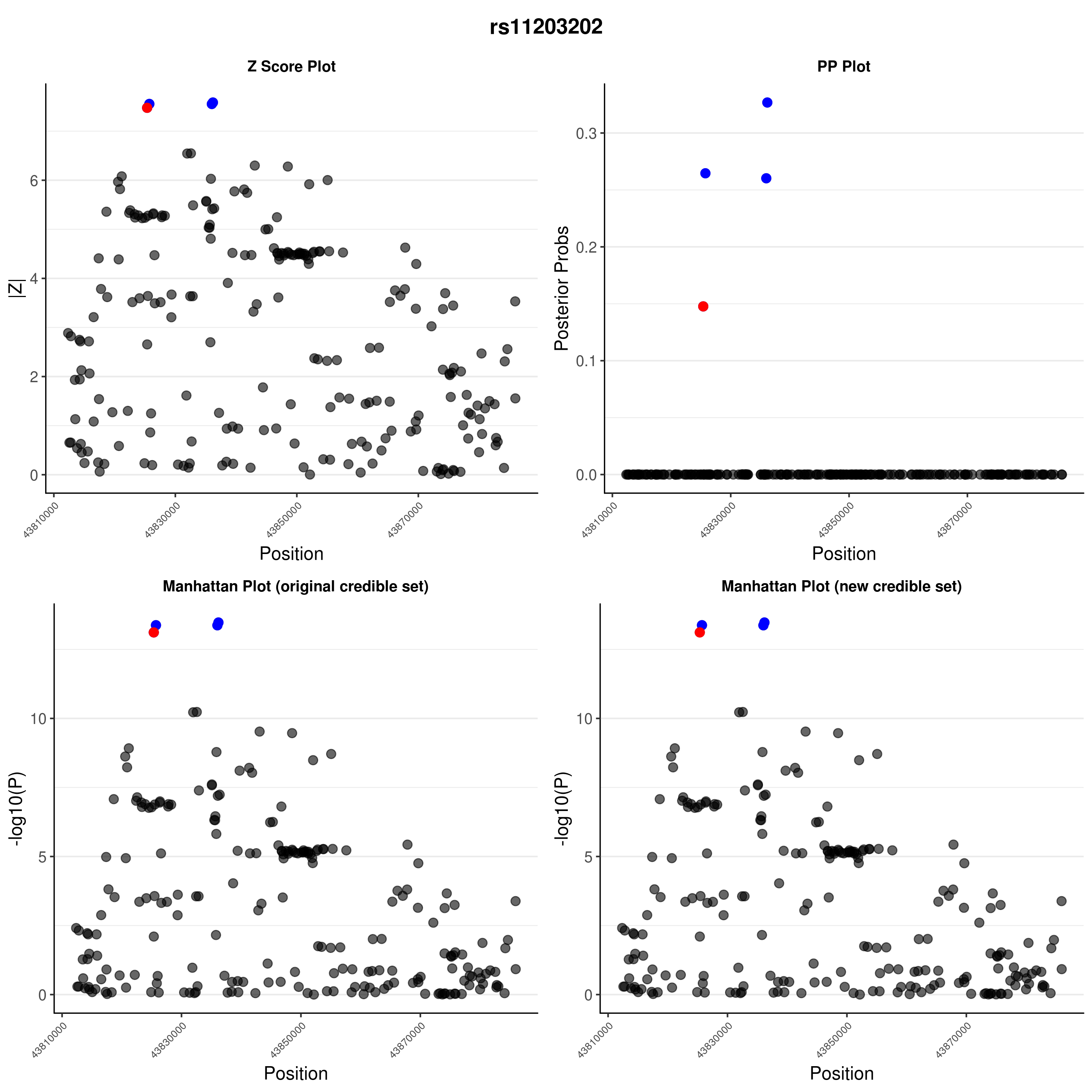

### rs11203202.png

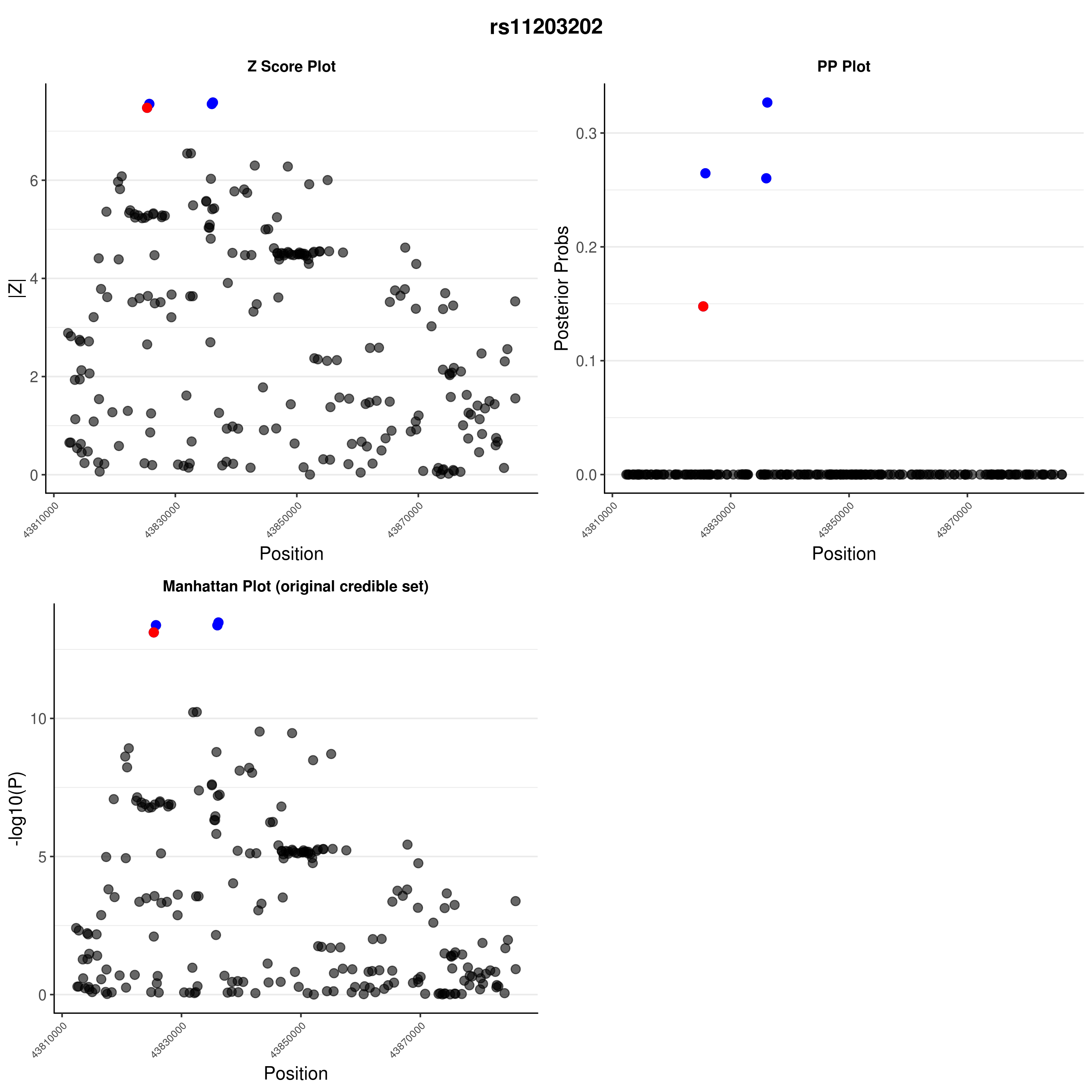

### rs11954020.png

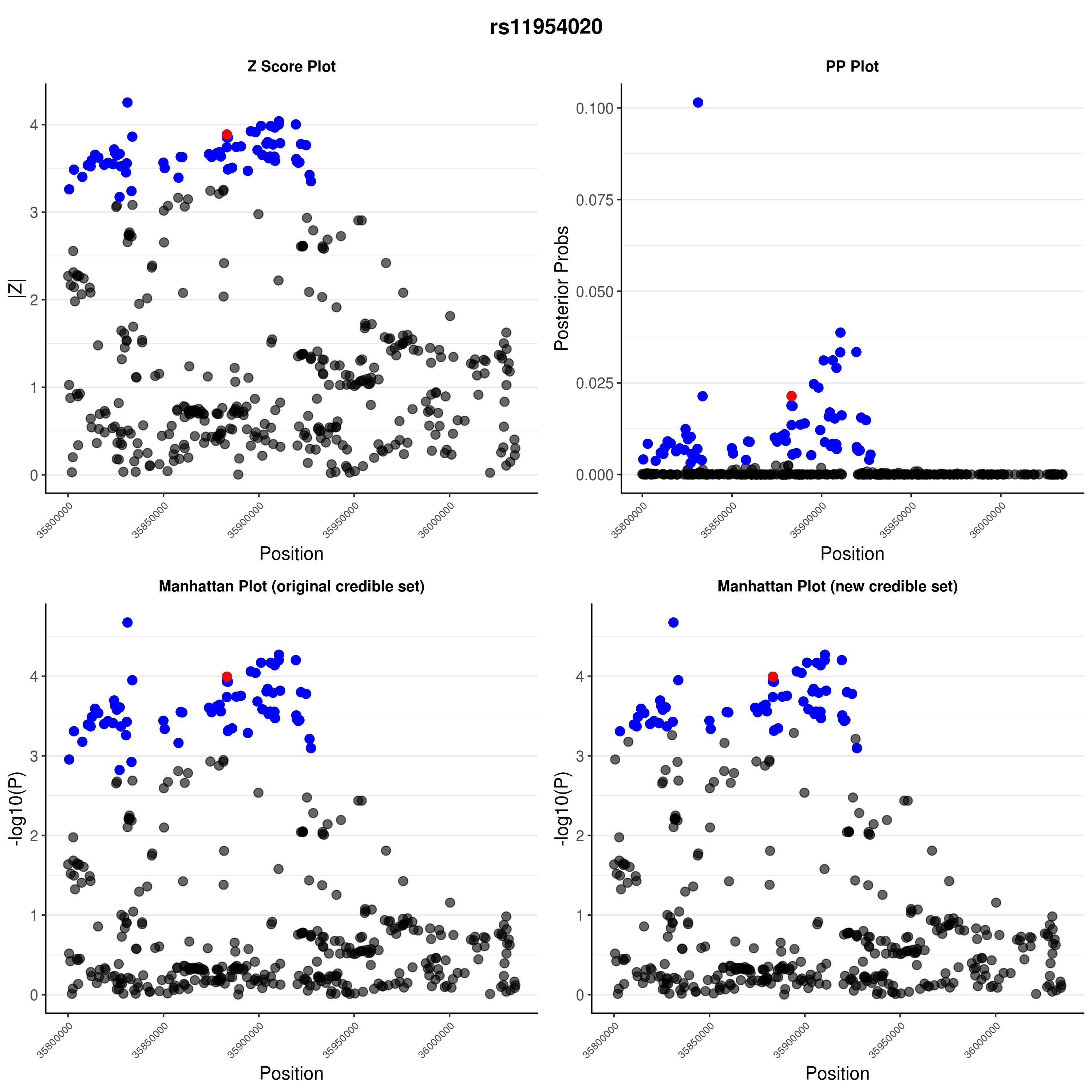

### rs11954020.png

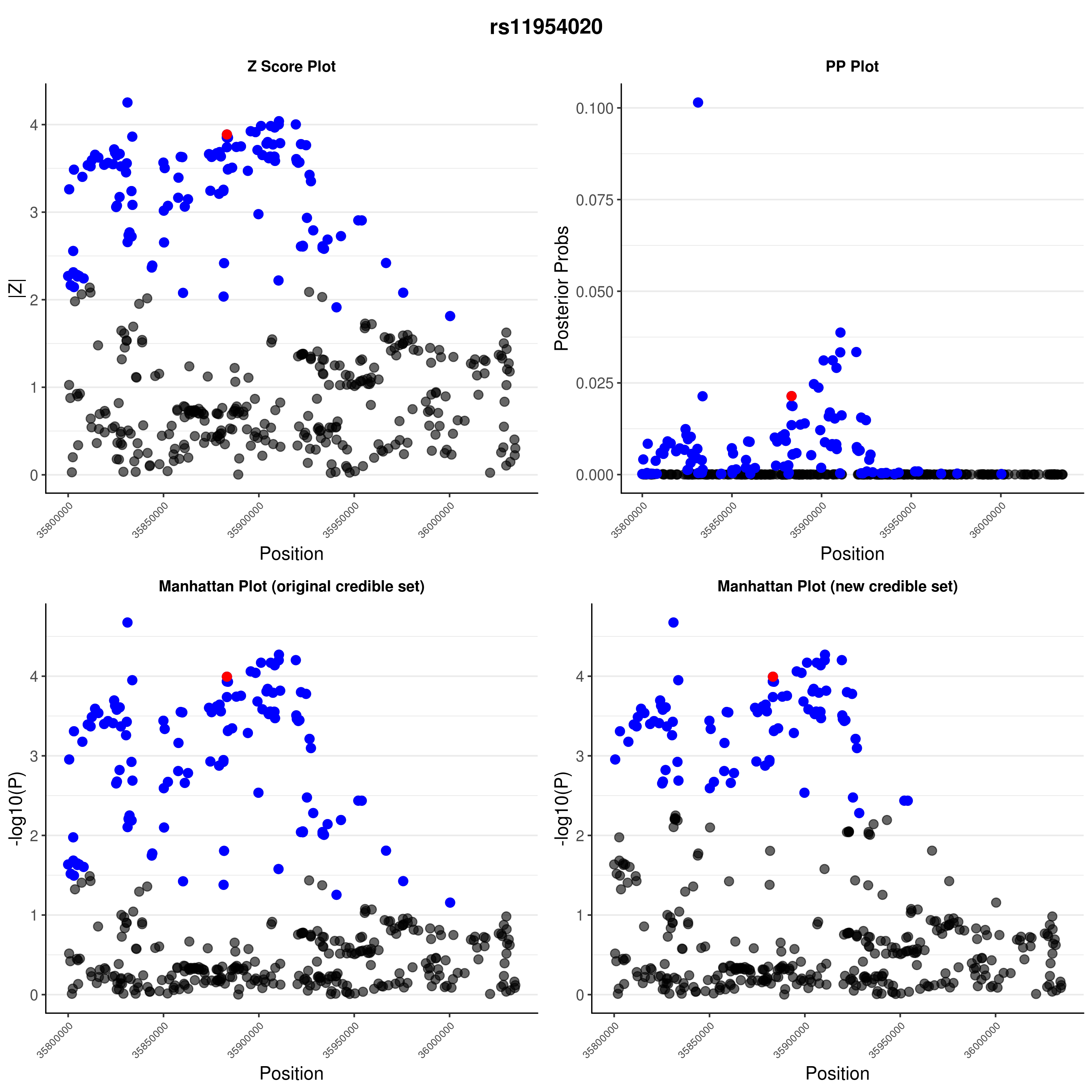

### rs12416116.png

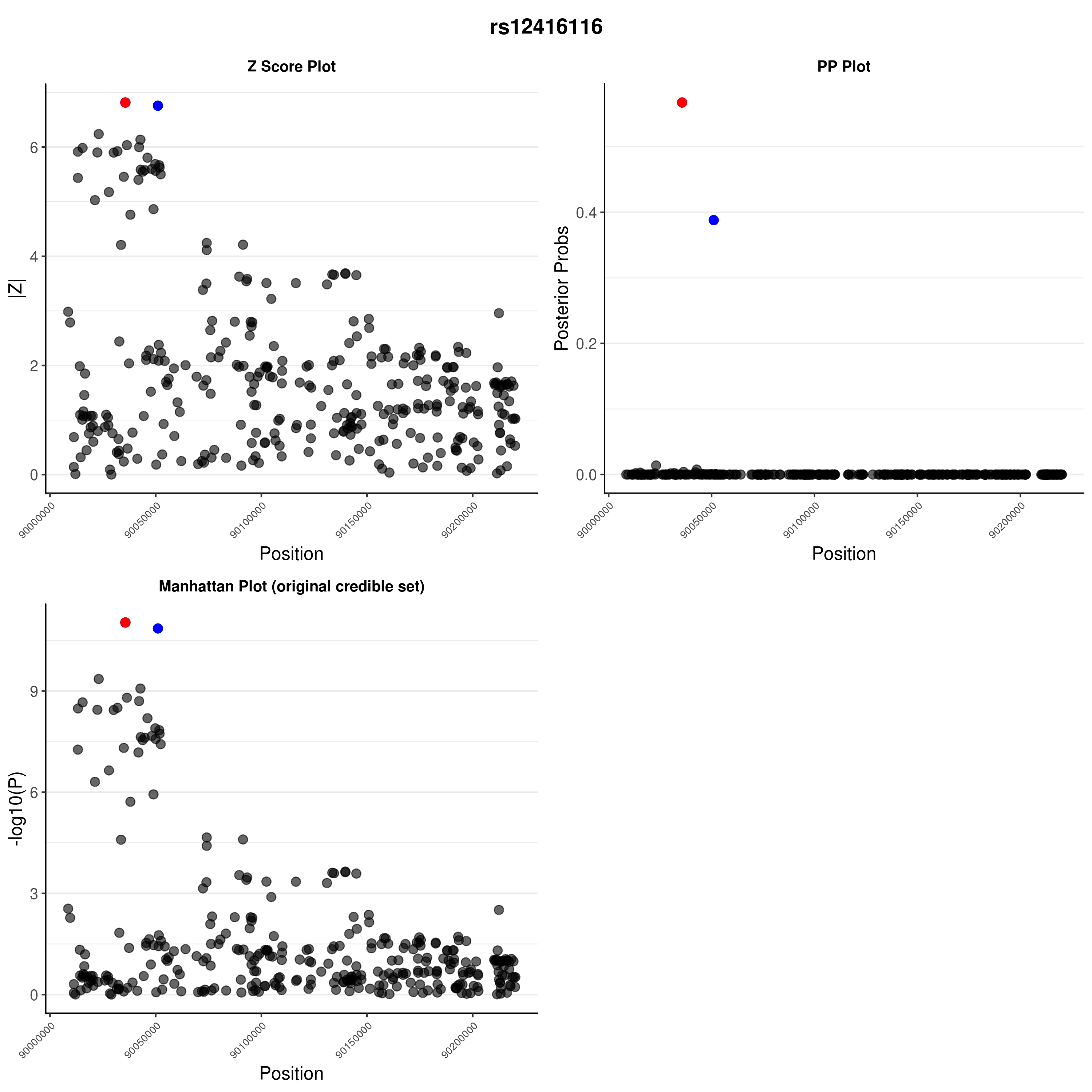

### rs12416116.png

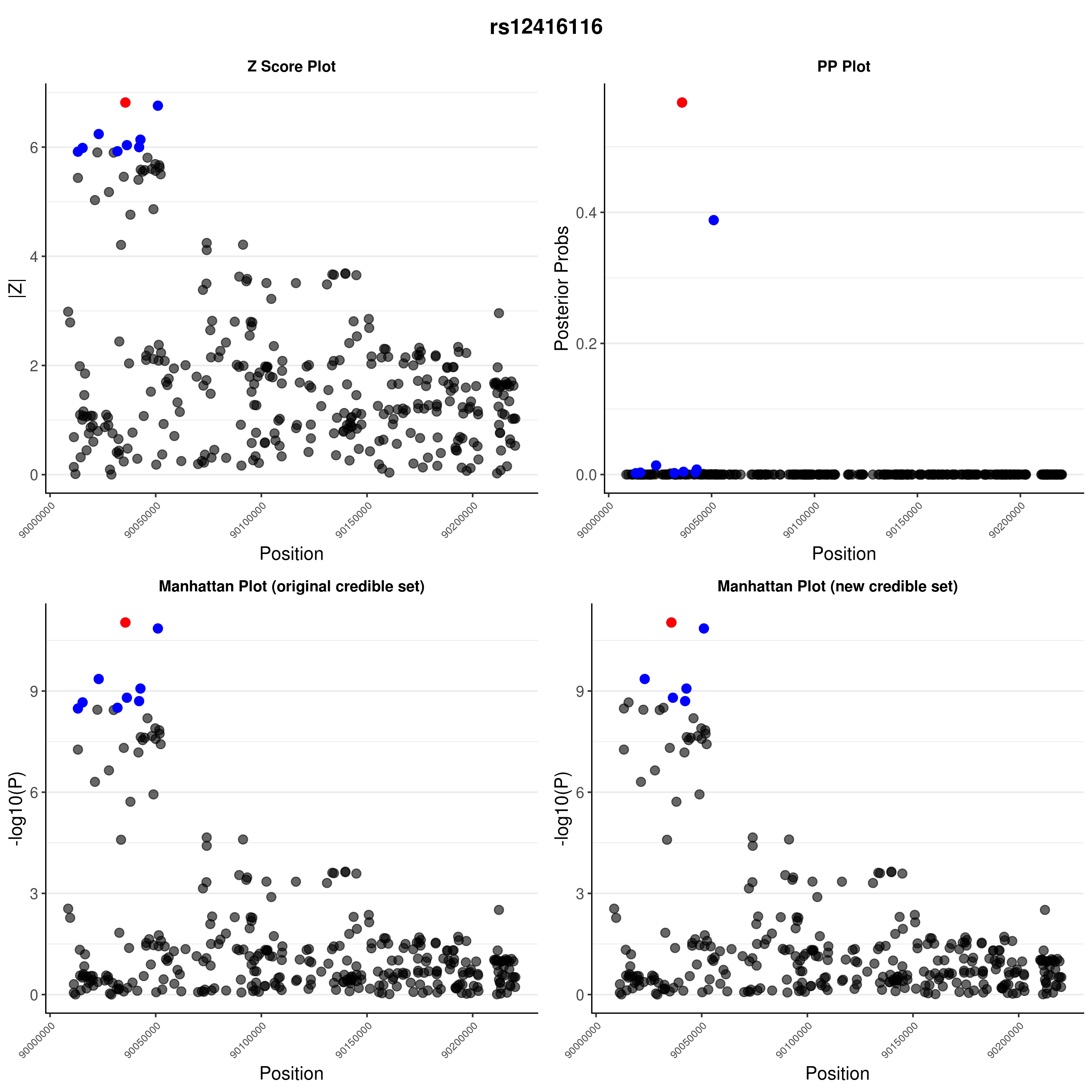

### rs12453507.png

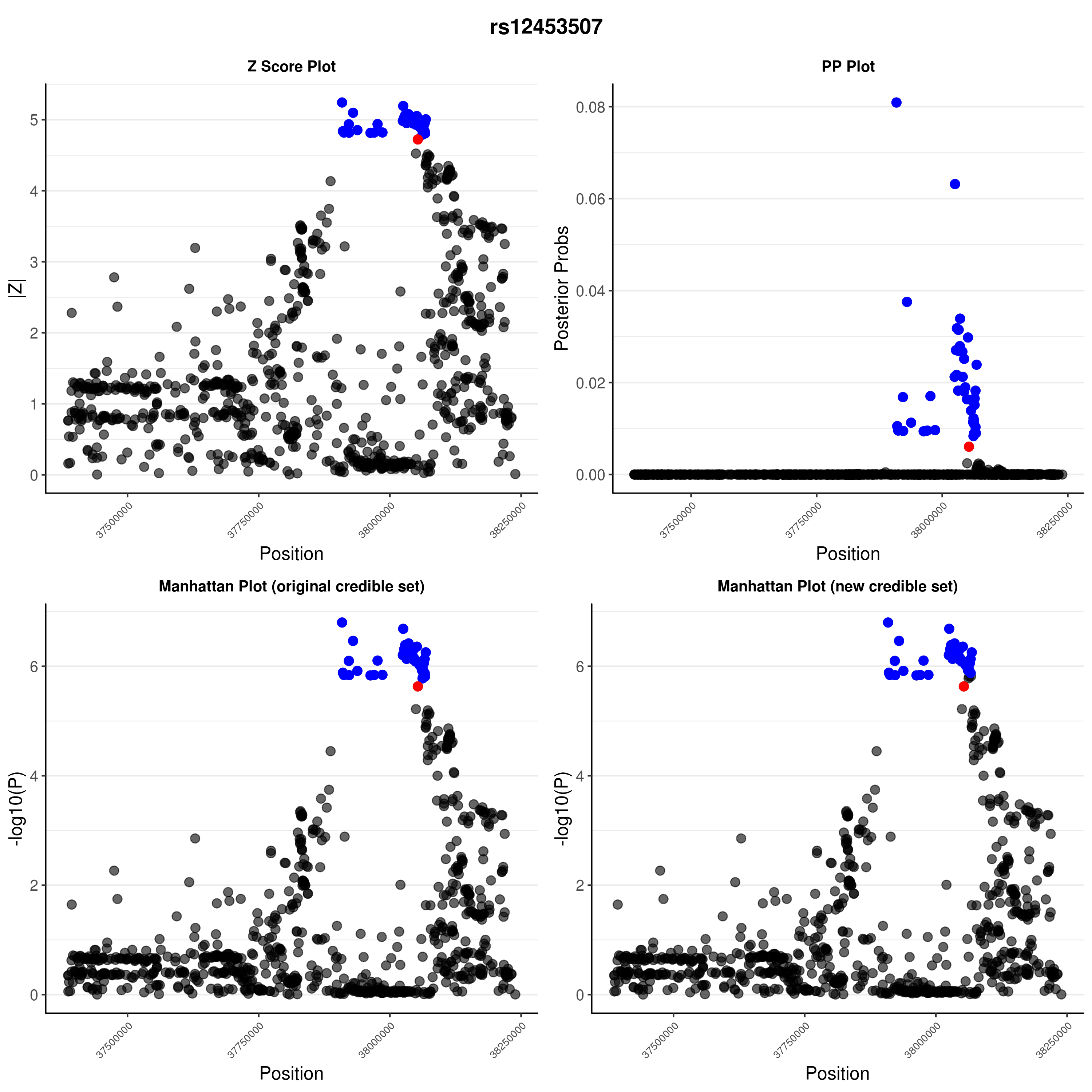

### rs12453507.png

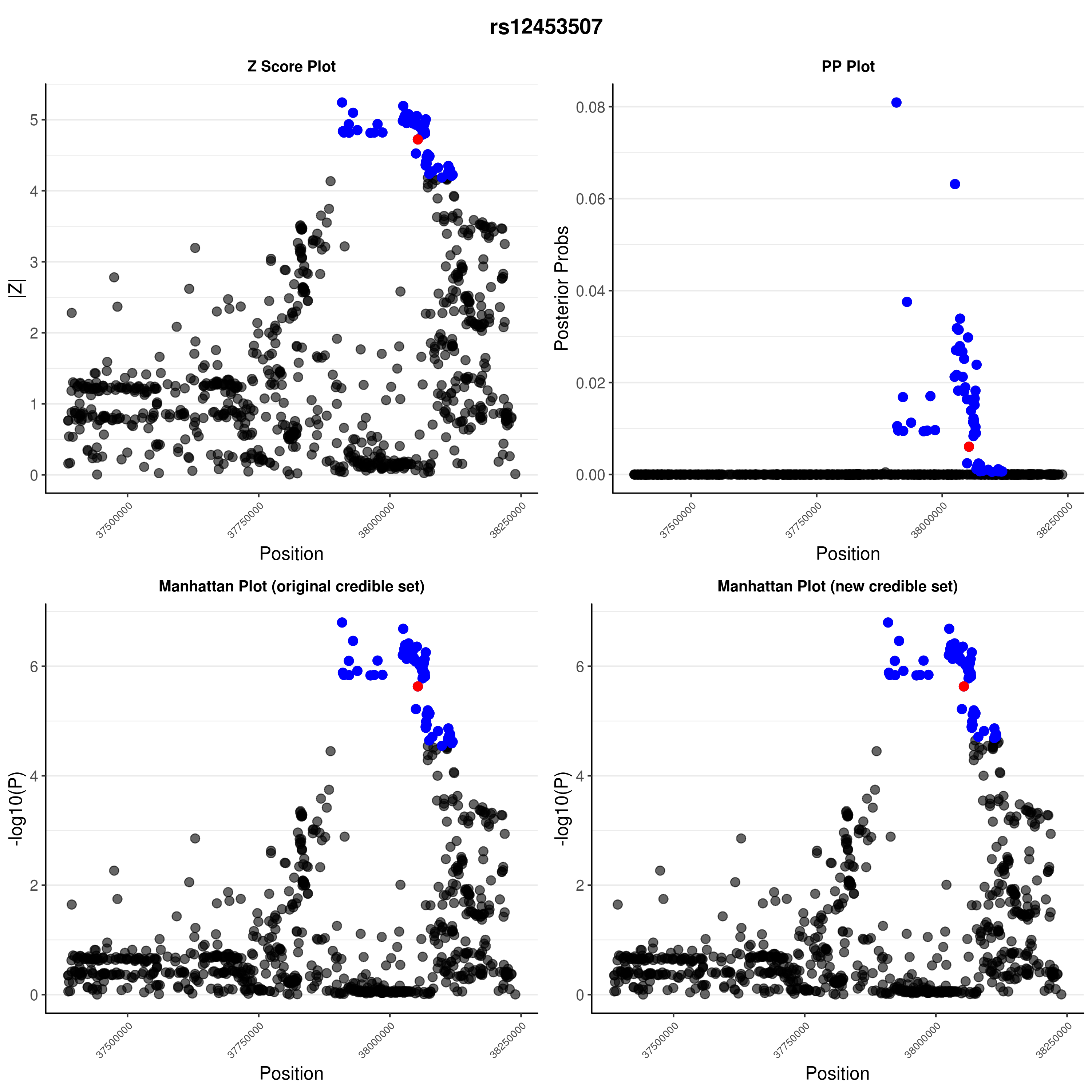

### rs12927355.png

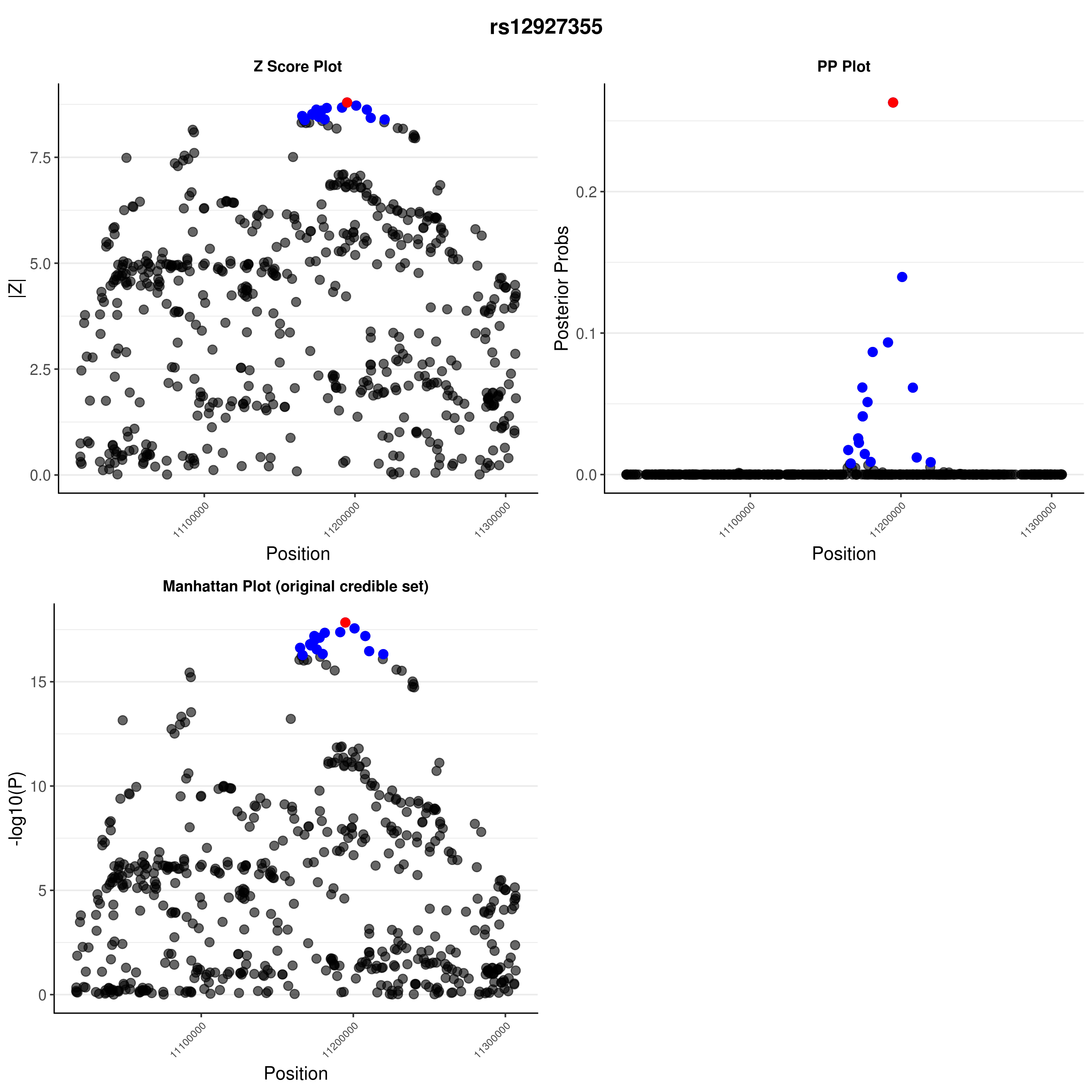

### rs12927355.png

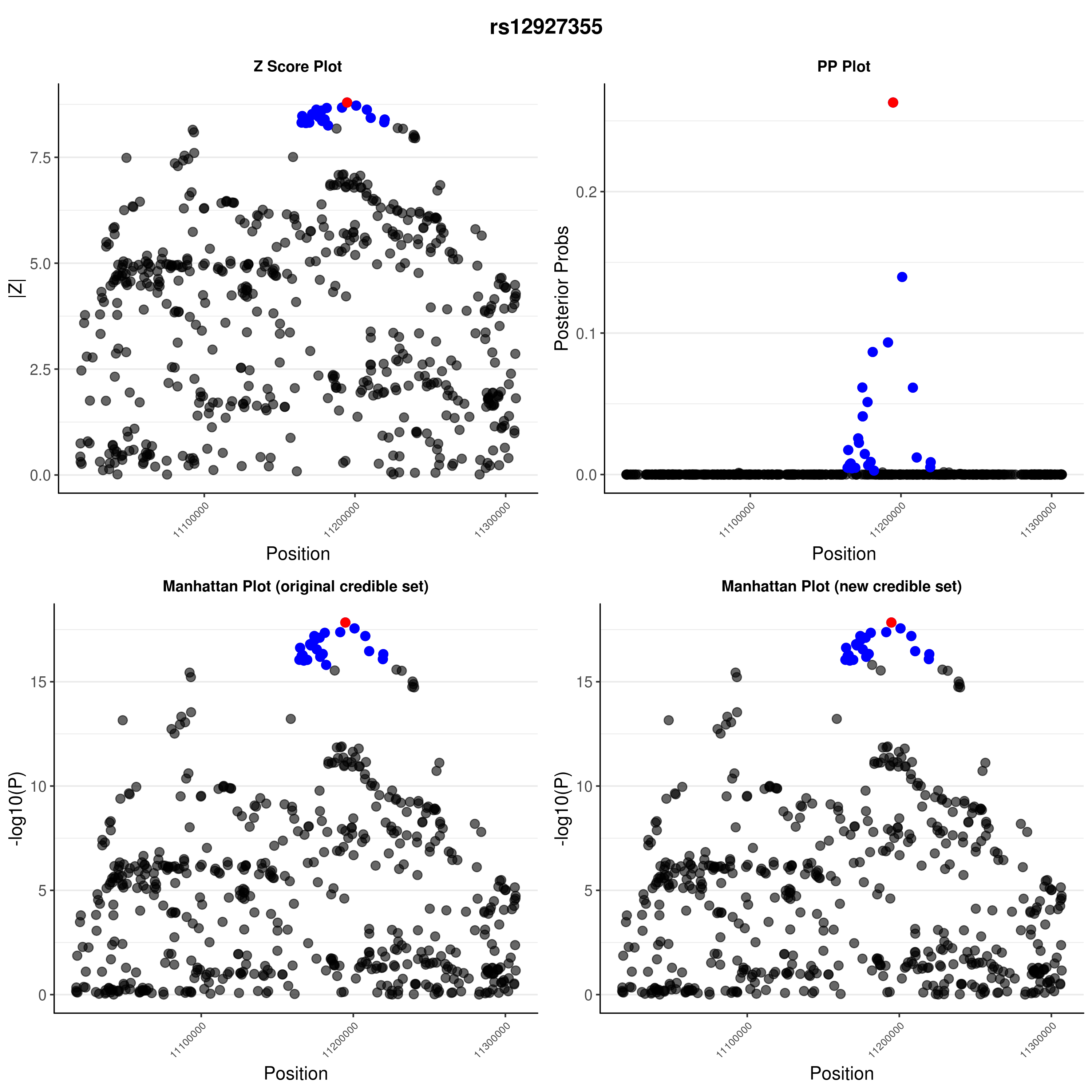

### rs13415583.png

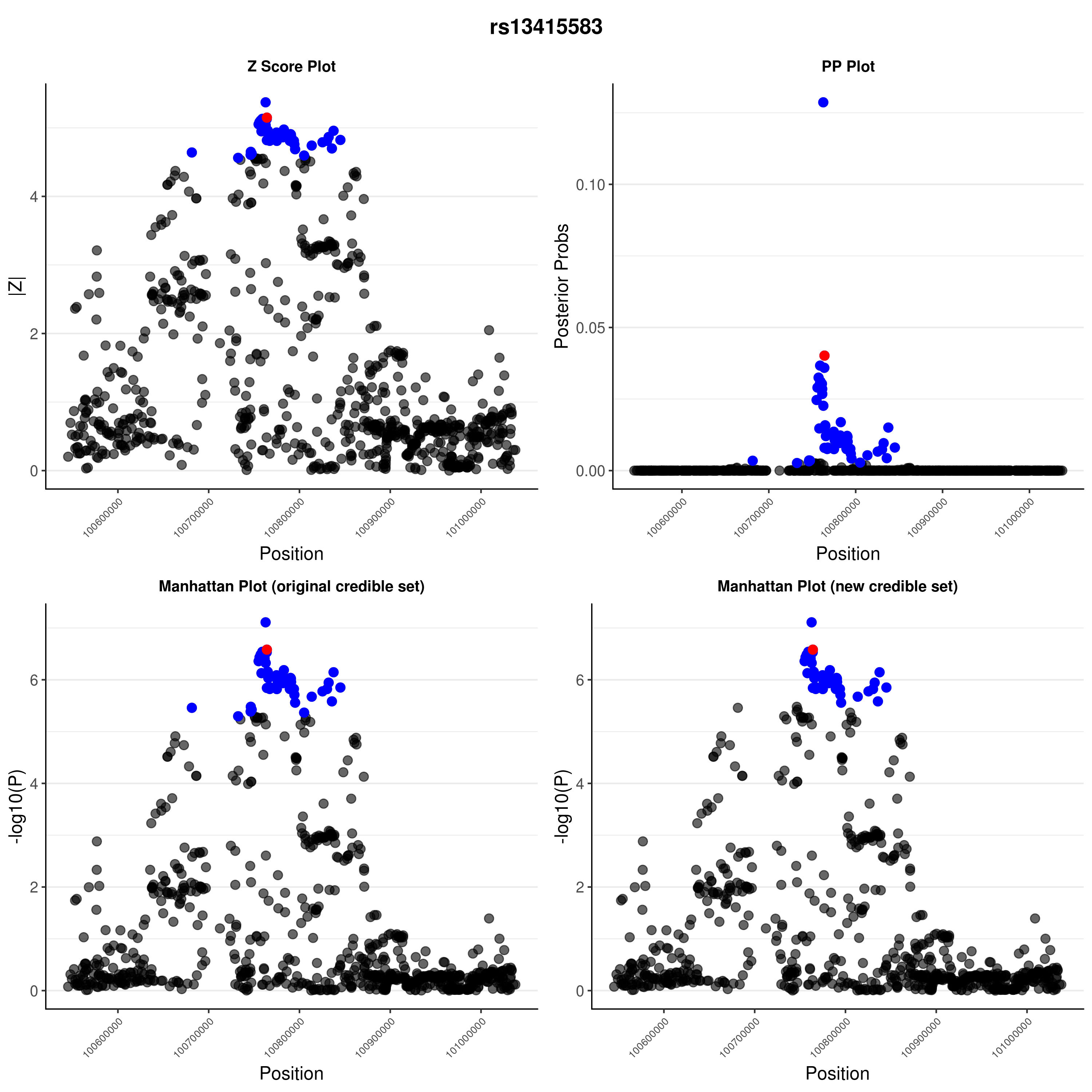

### rs13415583.png

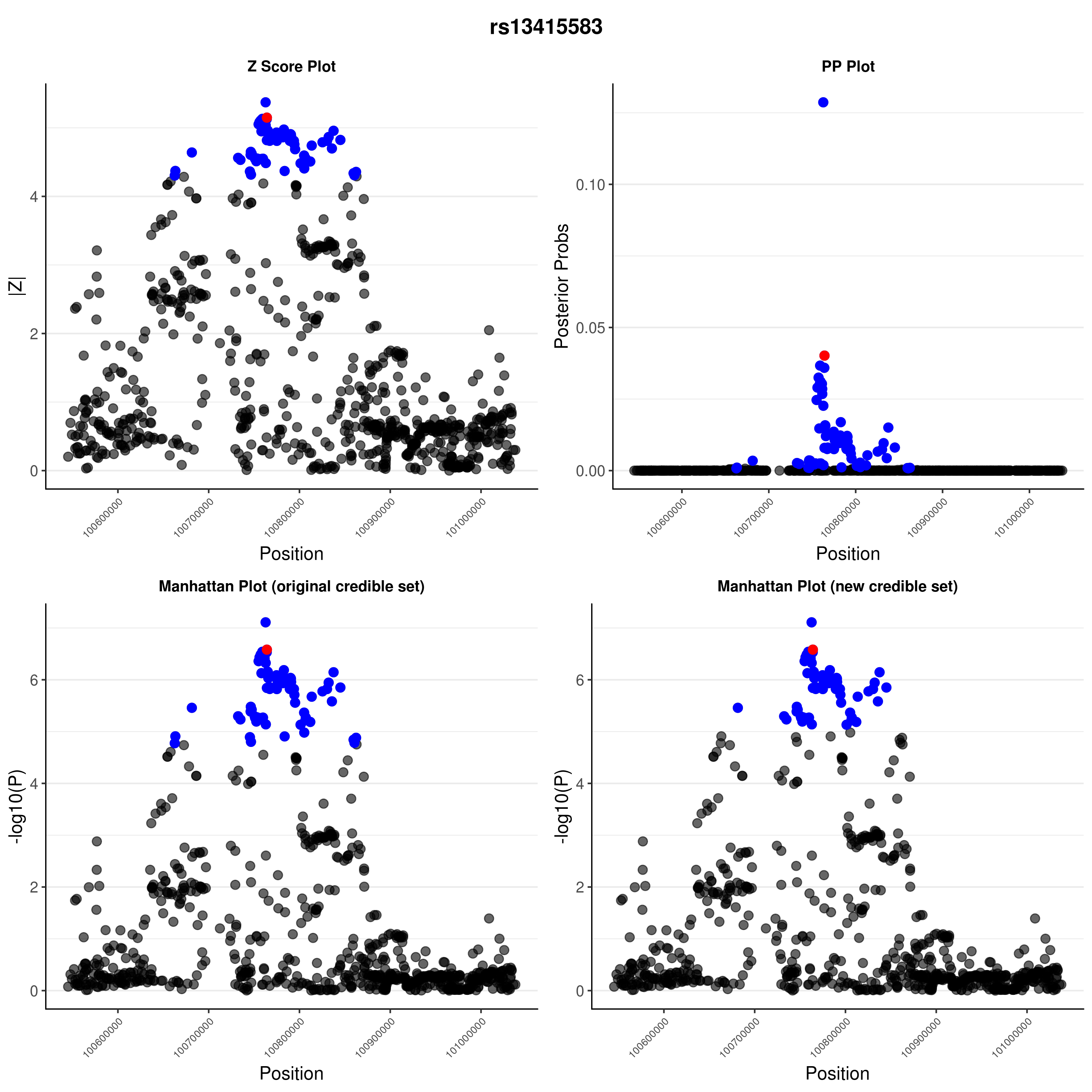

### rs34536443.png

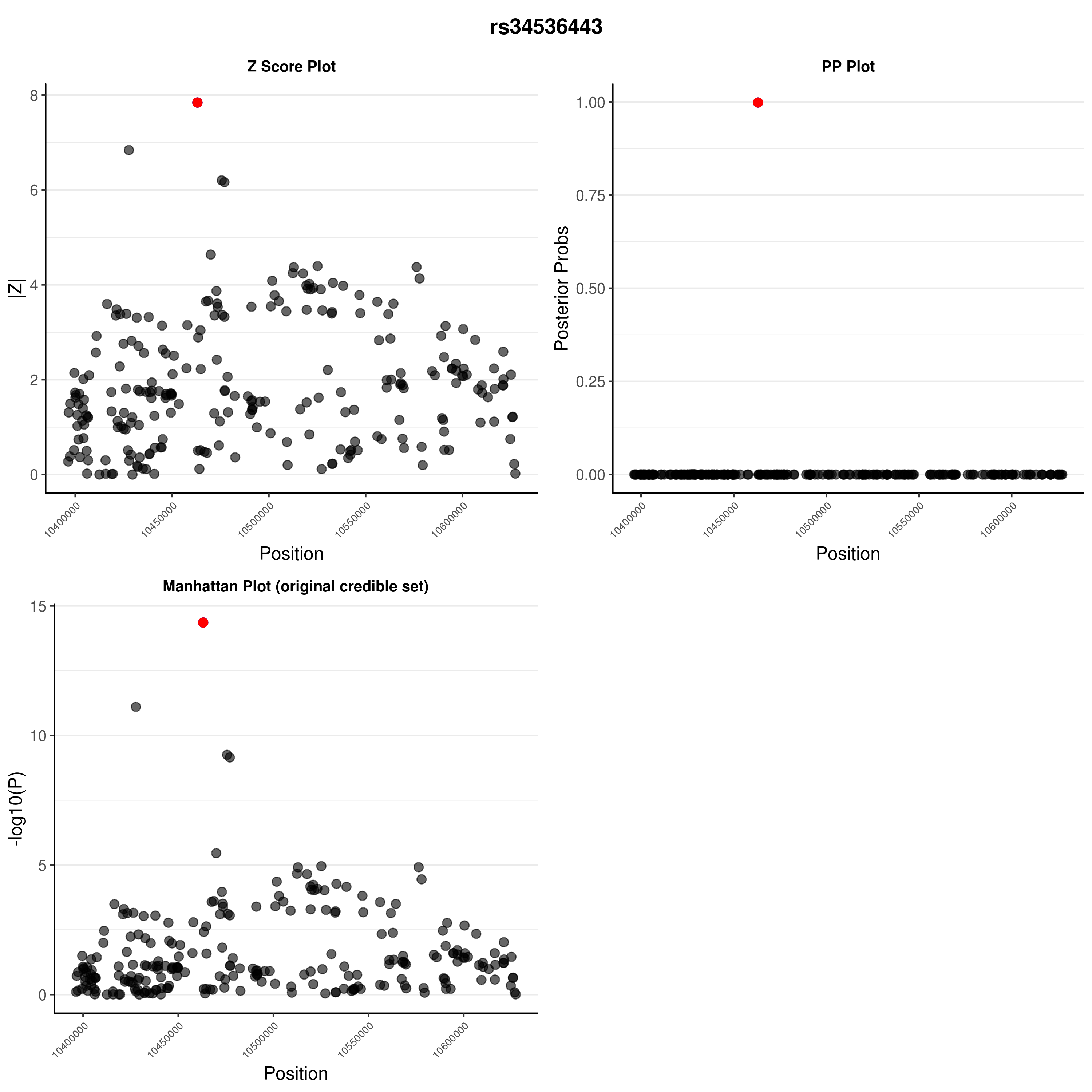

### rs34593439.png

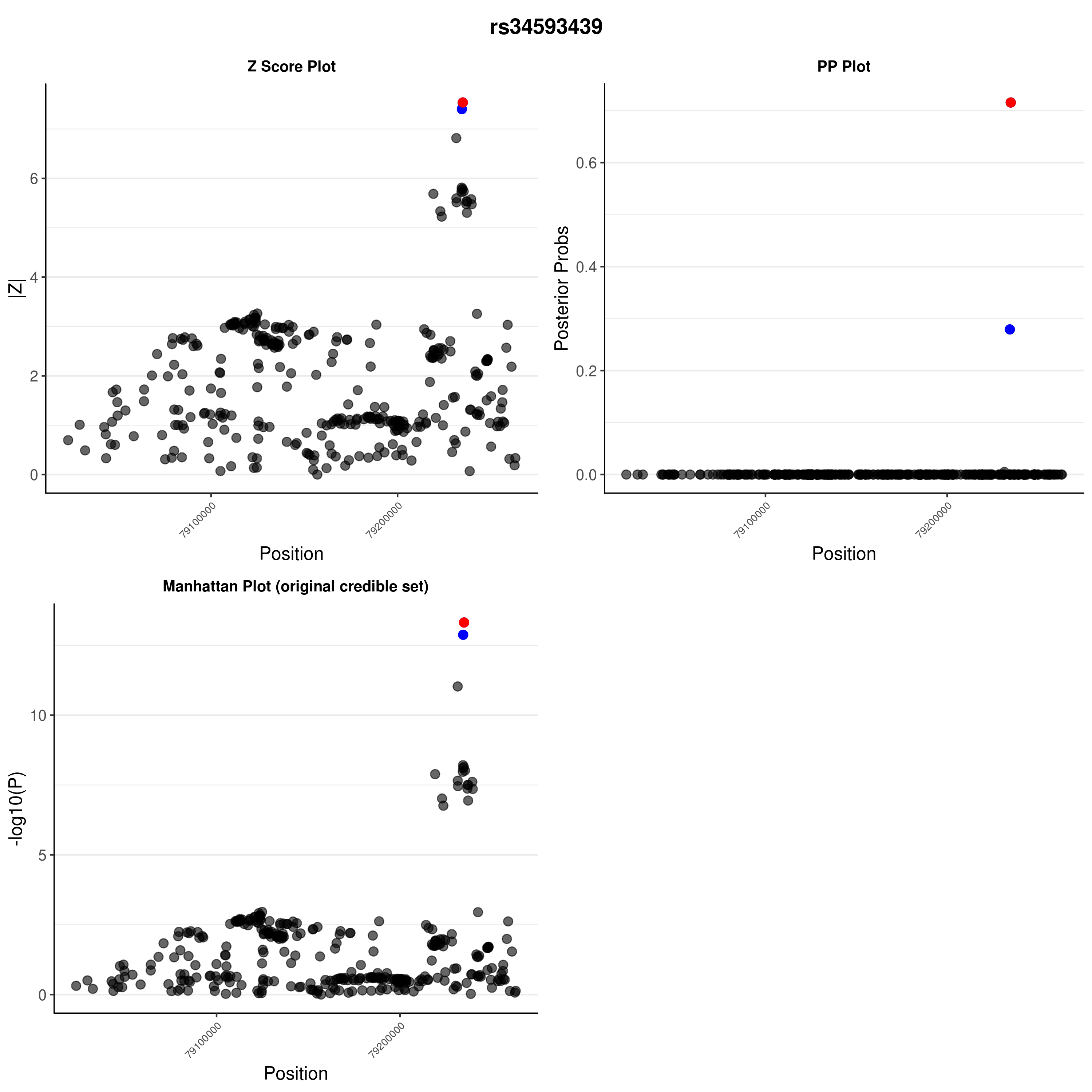

### rs56994090.png

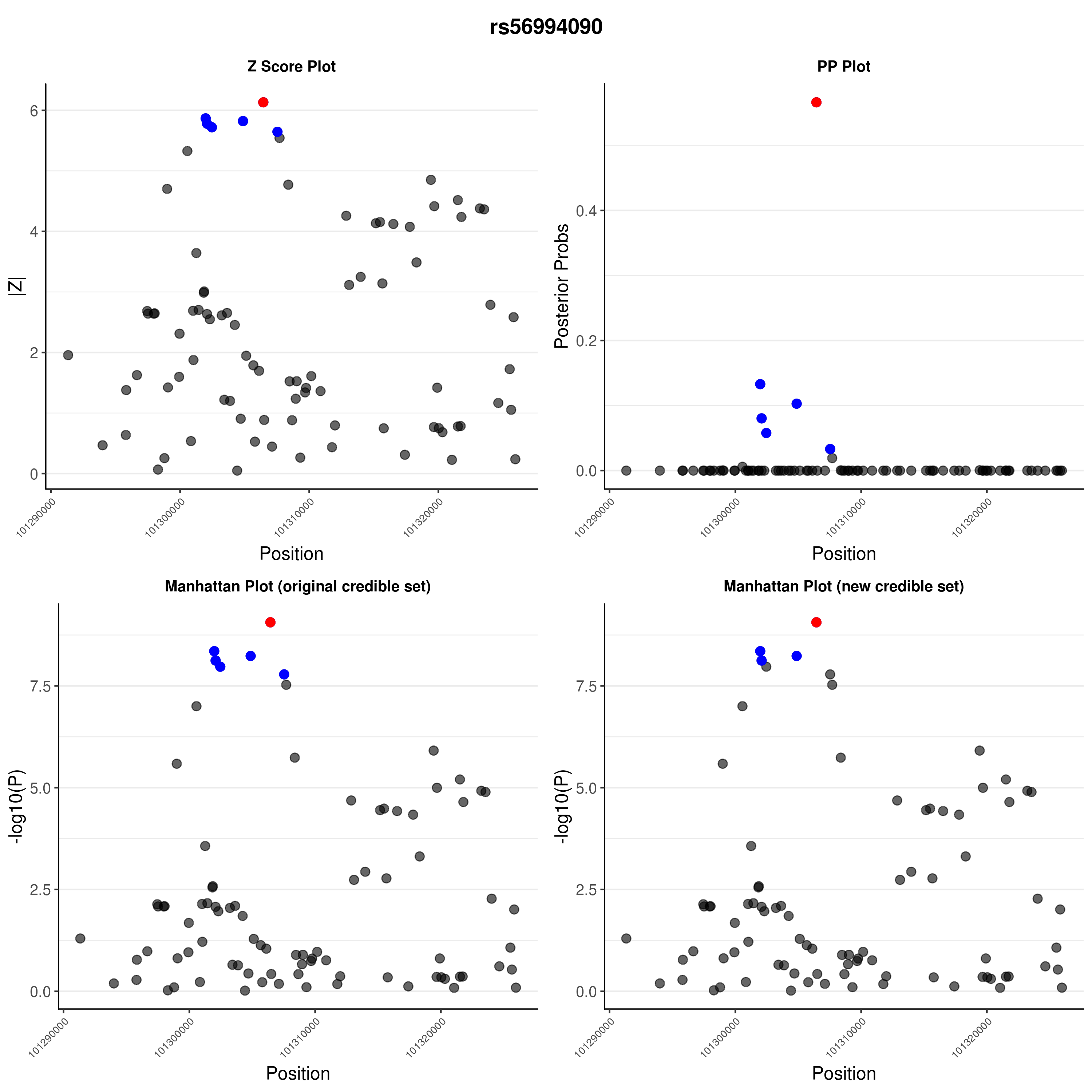

### rs56994090.png

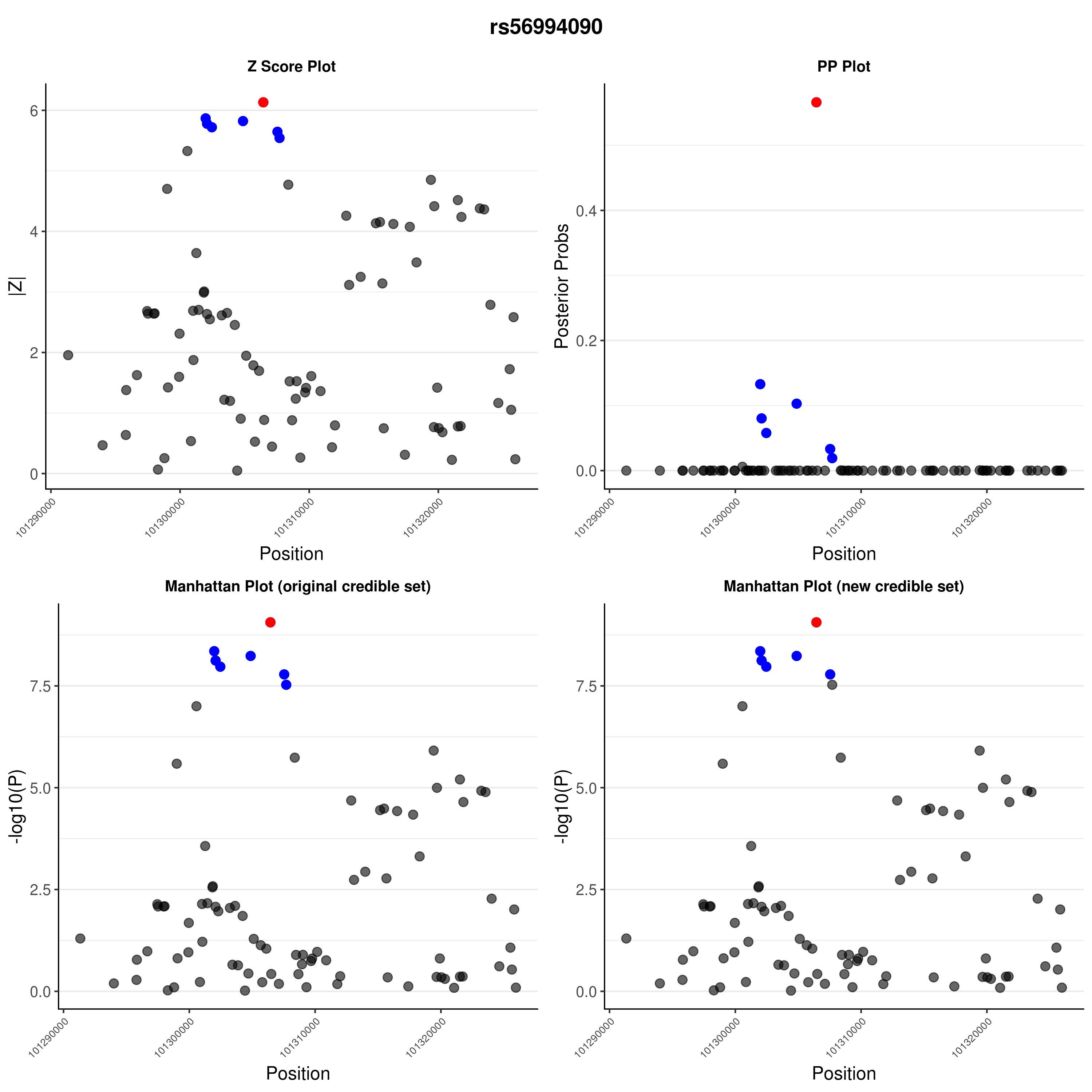

### rs61839660.png

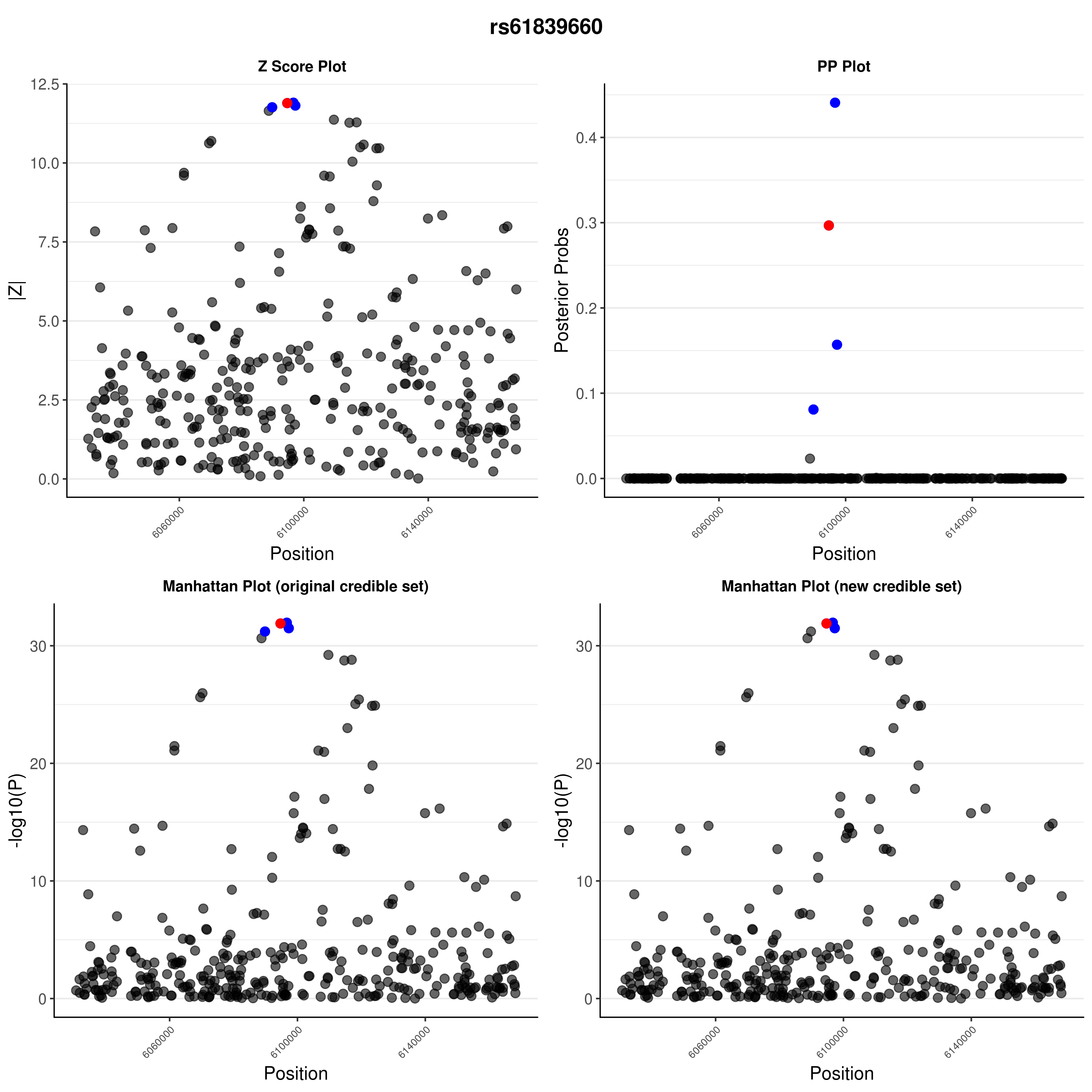

### rs61839660.png

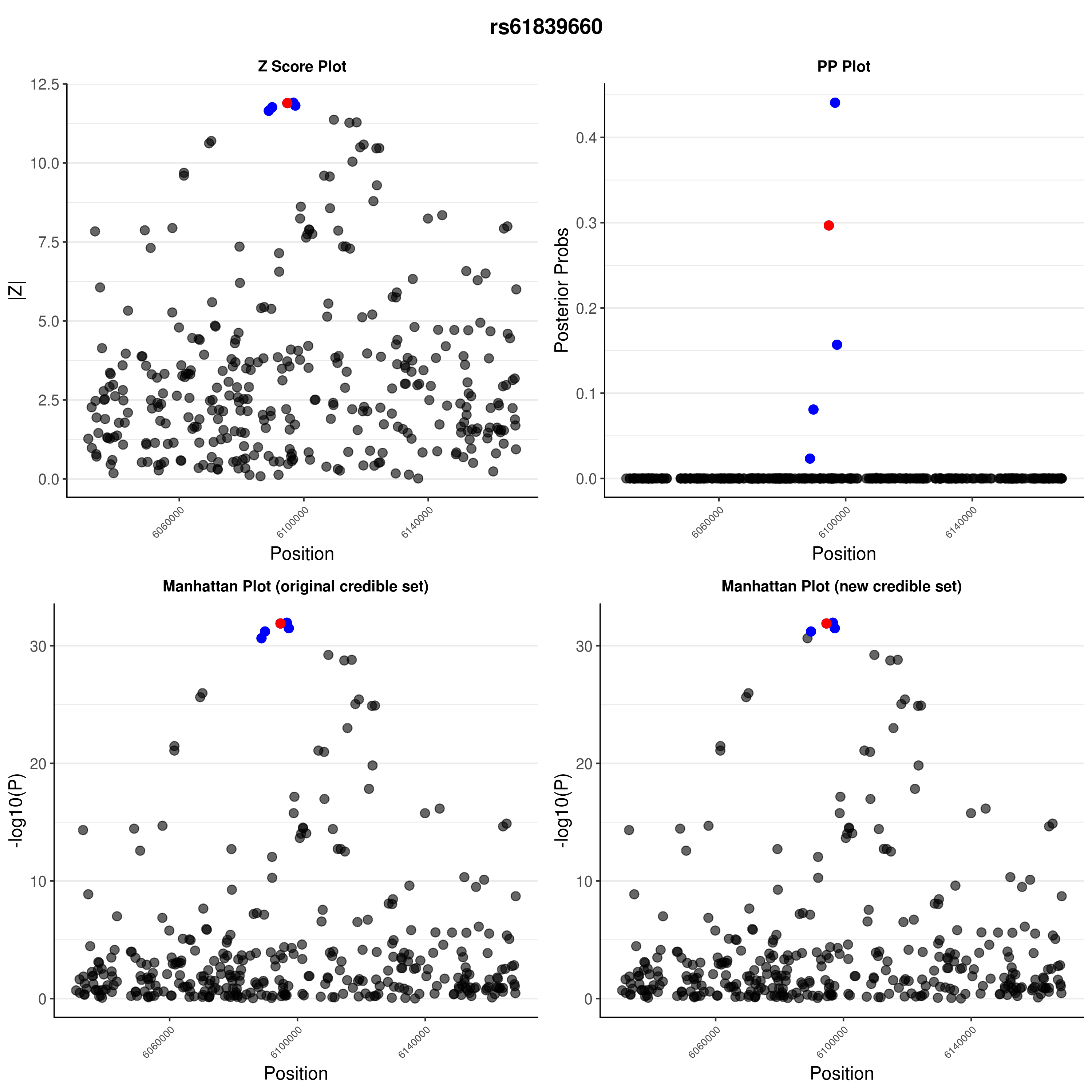

### rs62447205.png

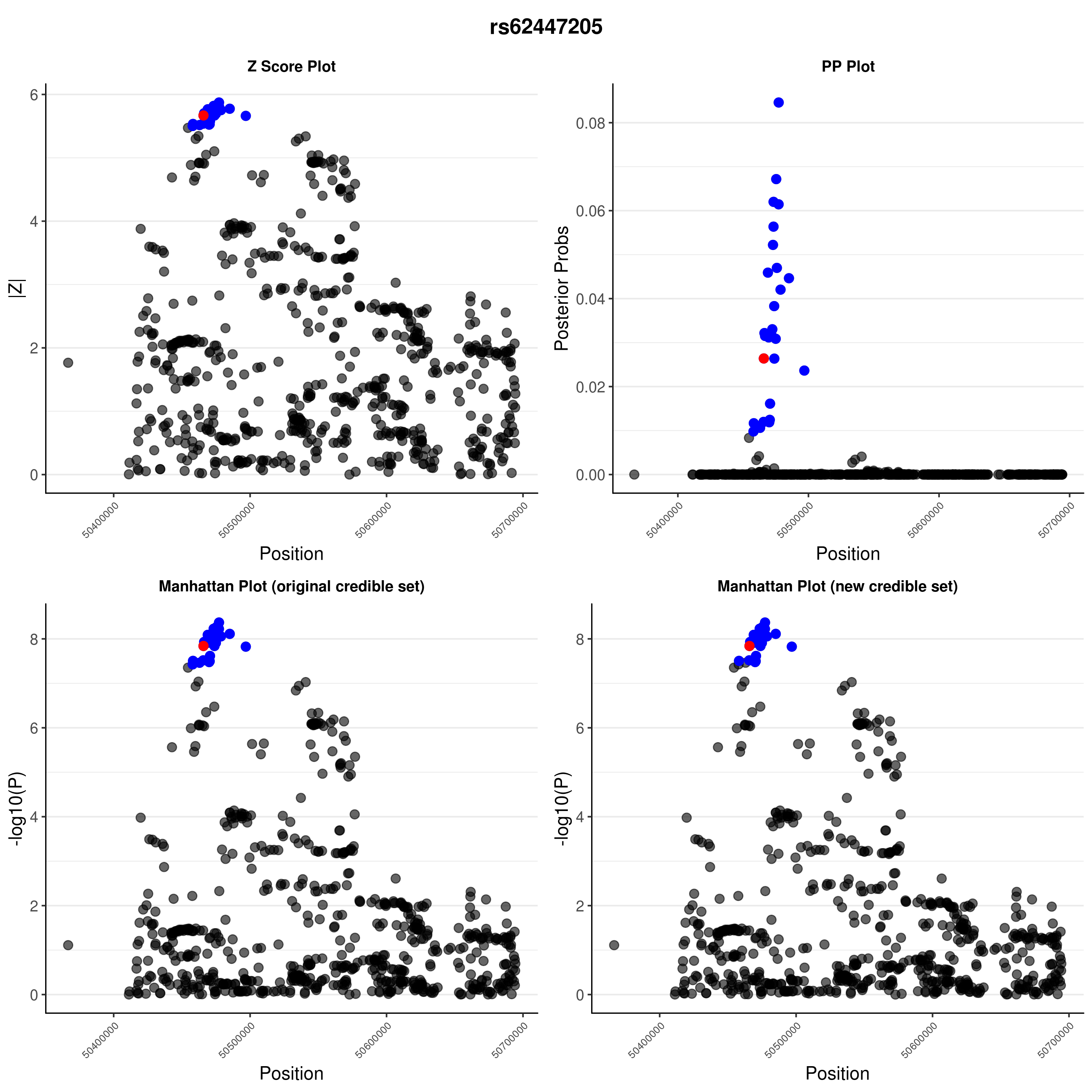

### rs62447205.png

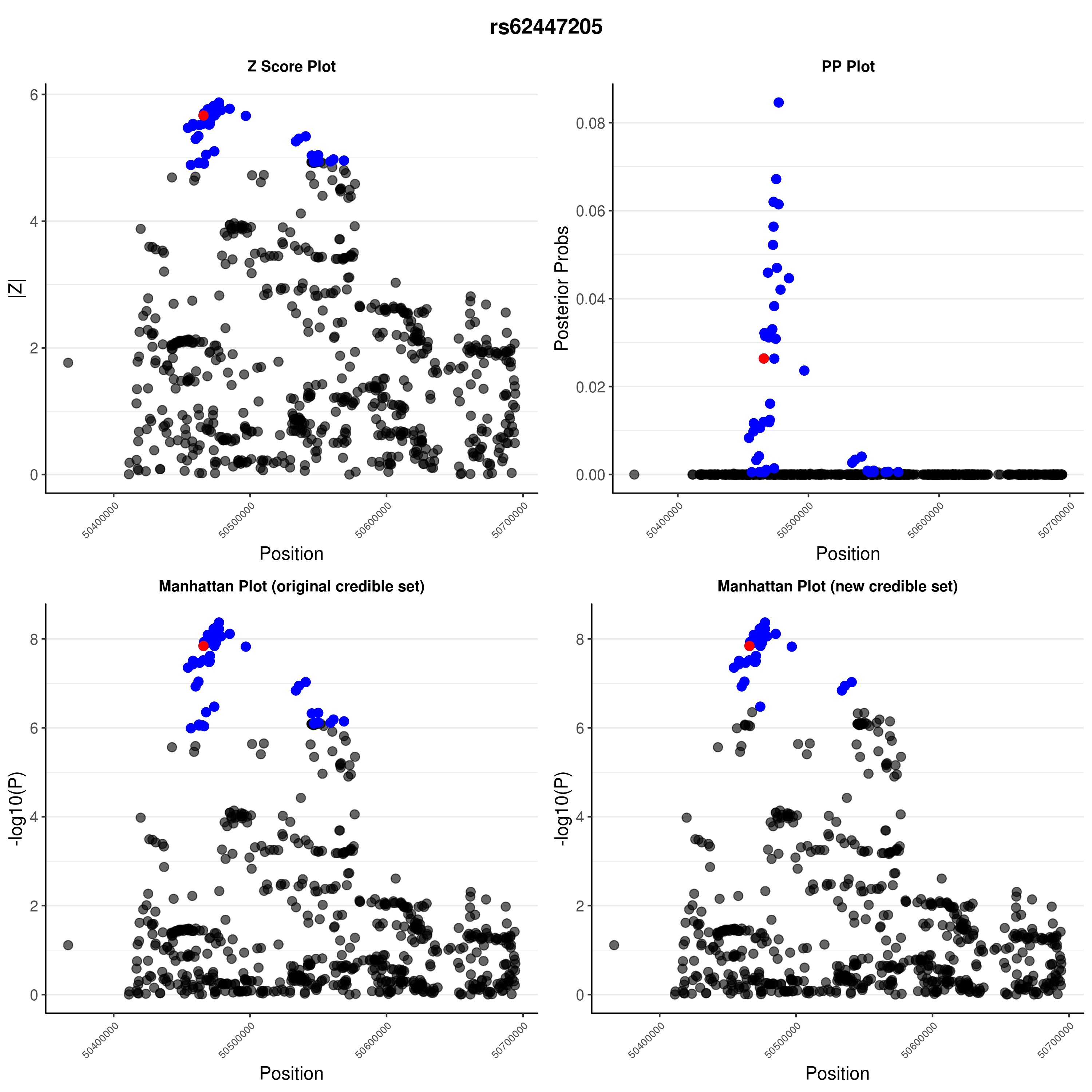

### rs72727394.png

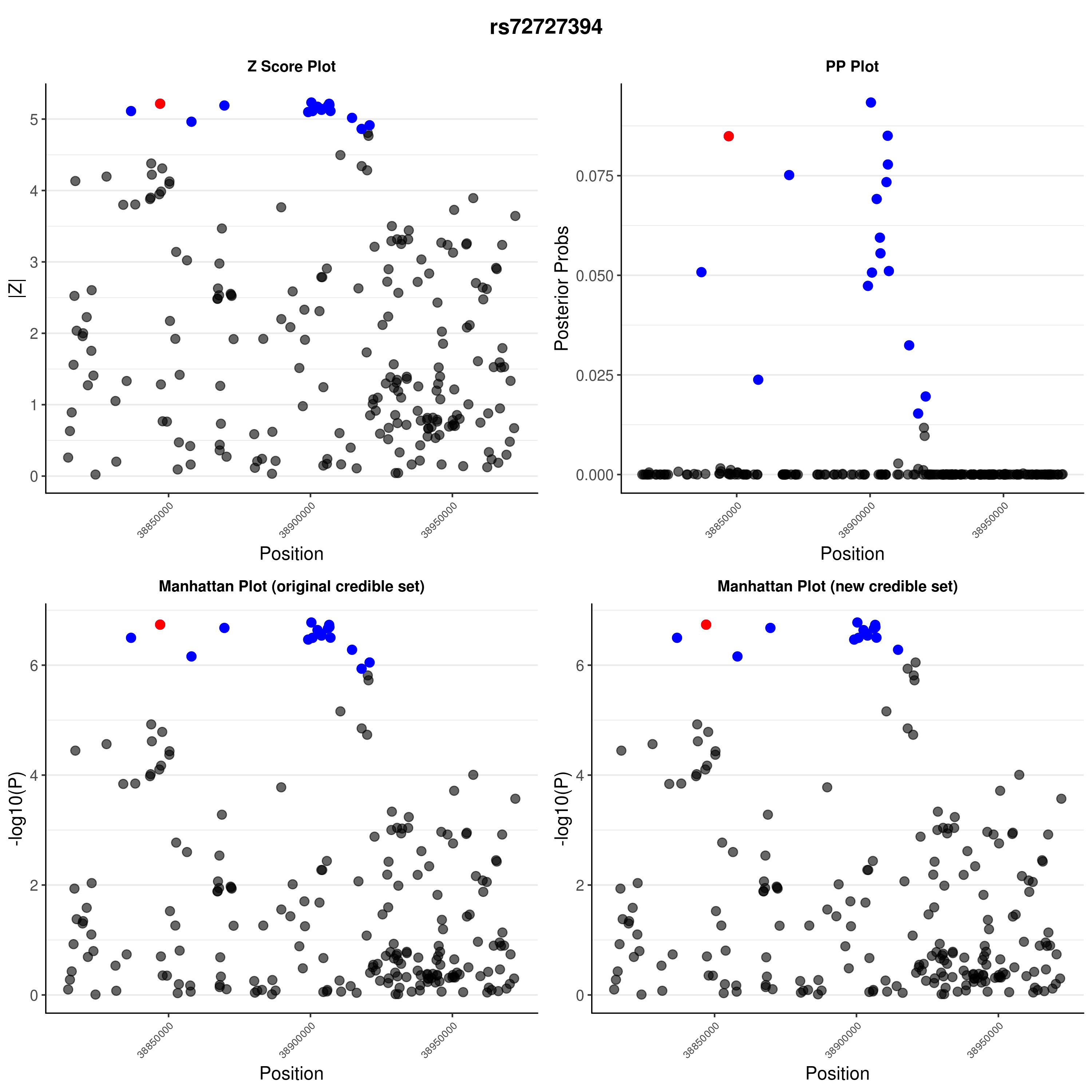

### rs72727394.png

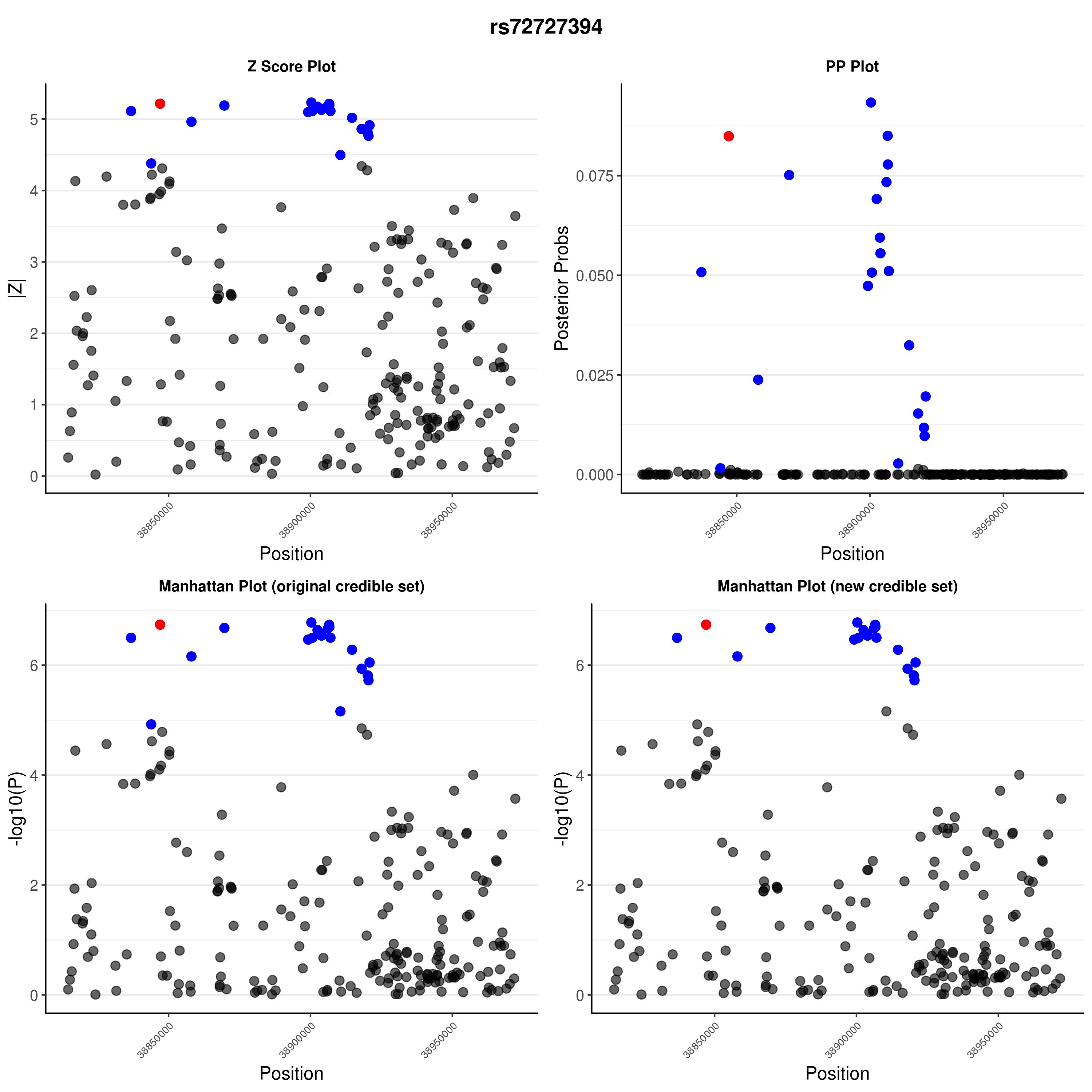

### rs72928038.png

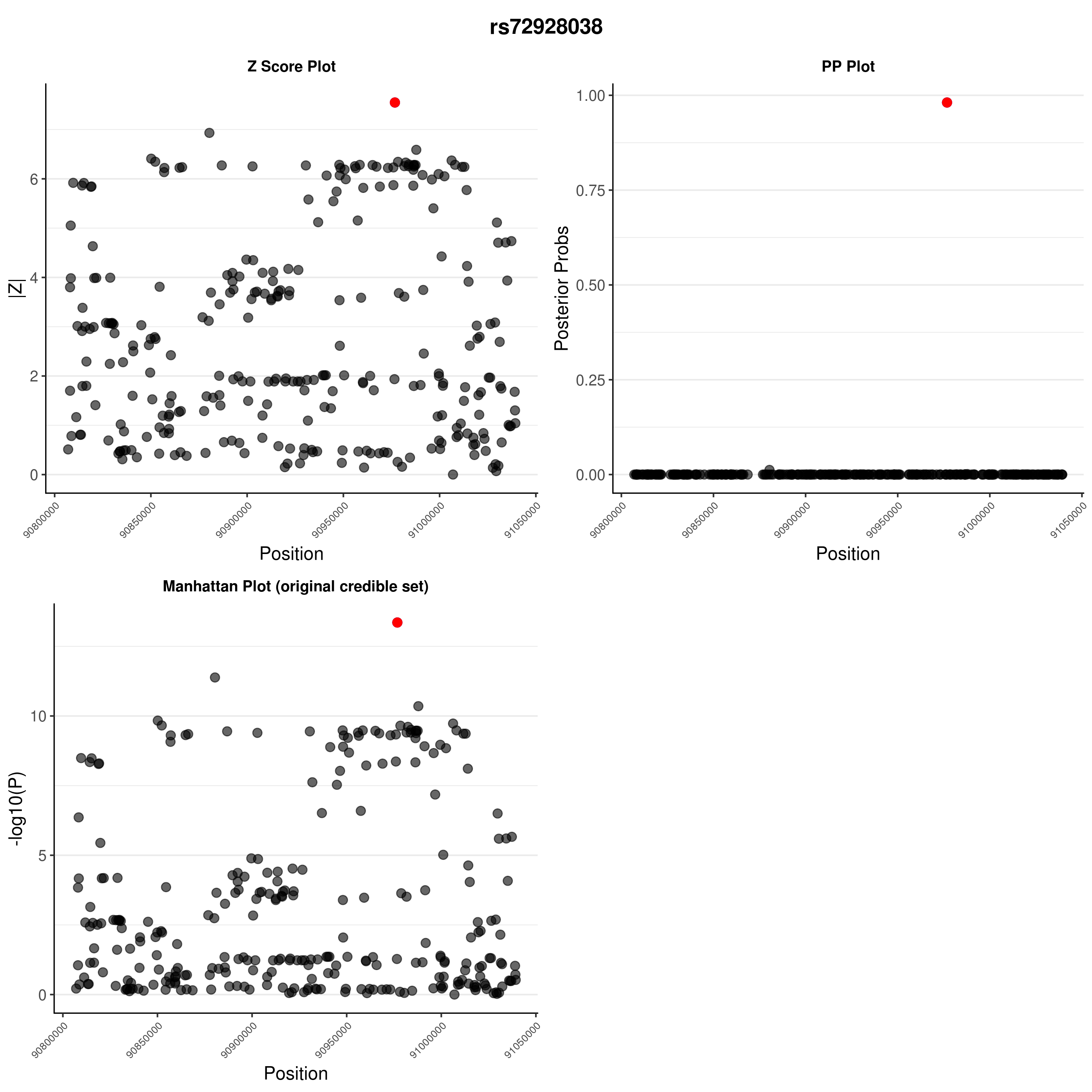

### rs72928038.png

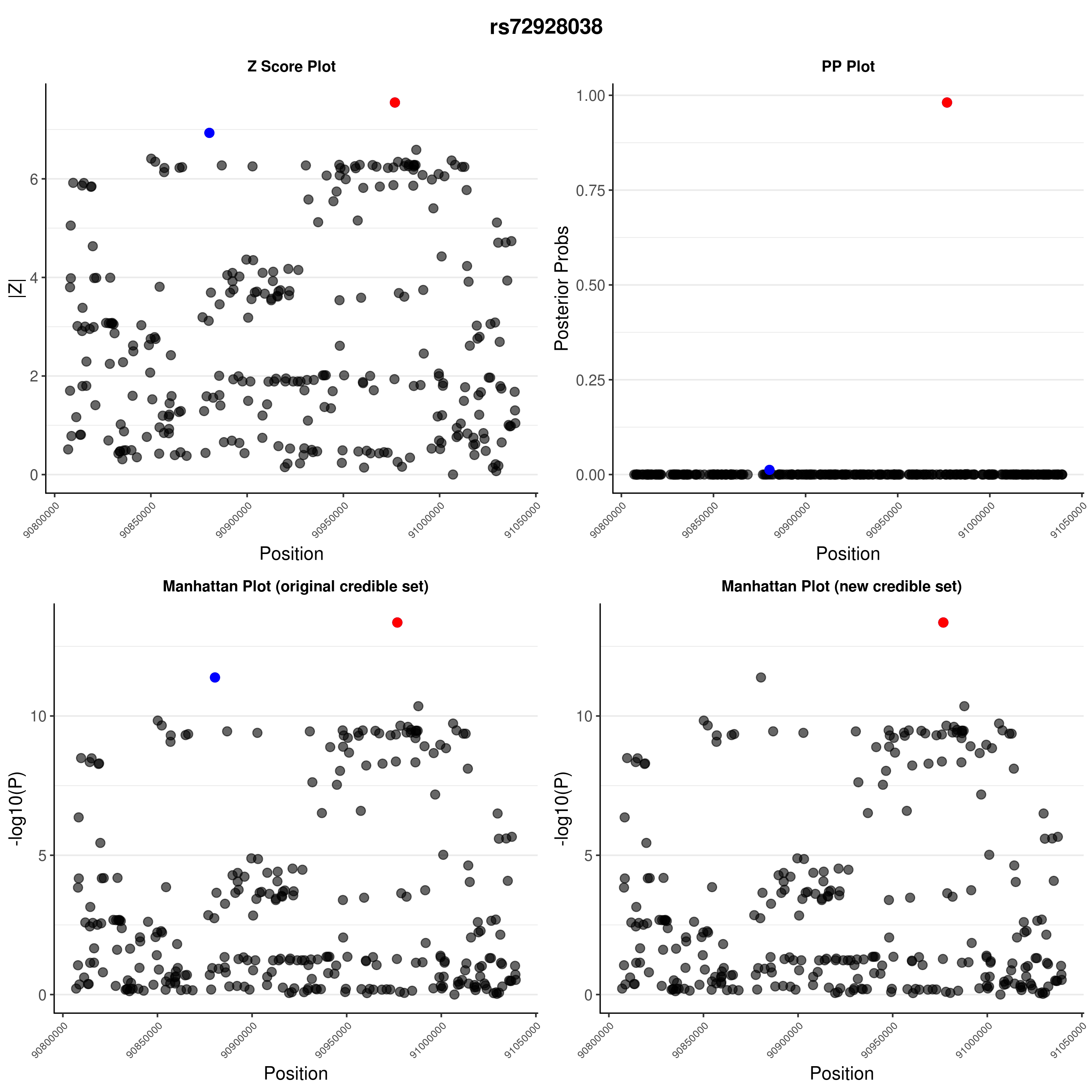

### rs75793288.png

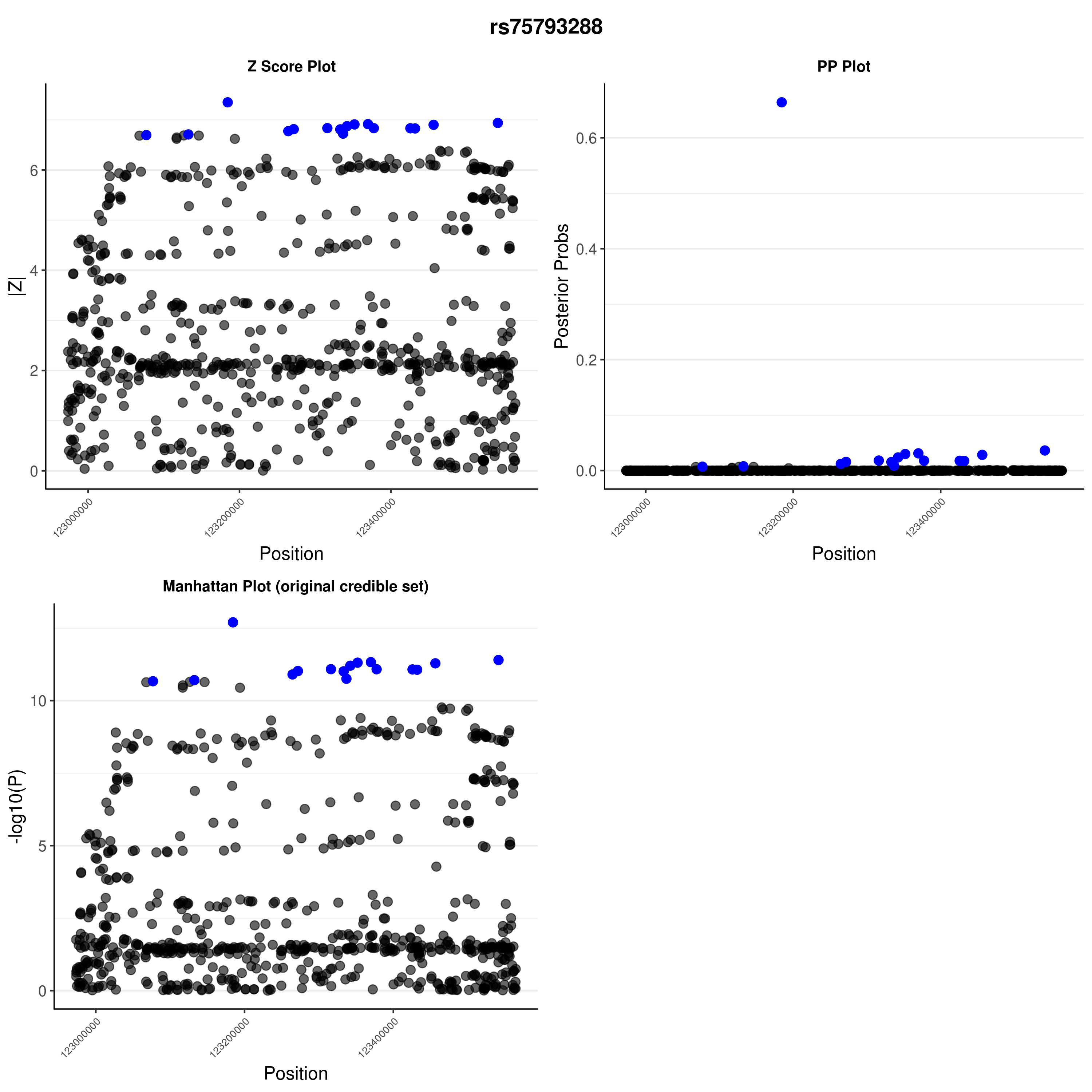

### rs75793288.png

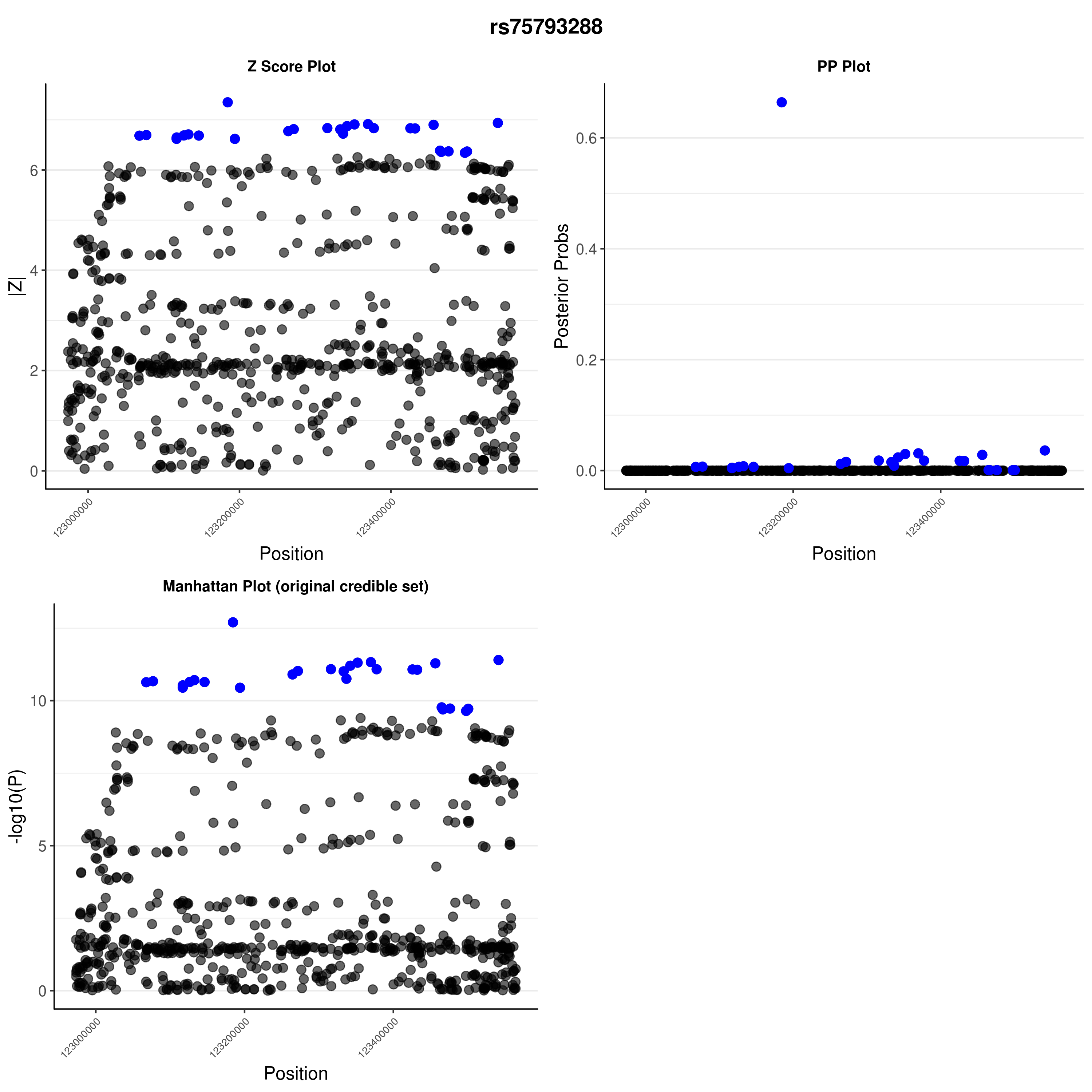

### rs113010081.png

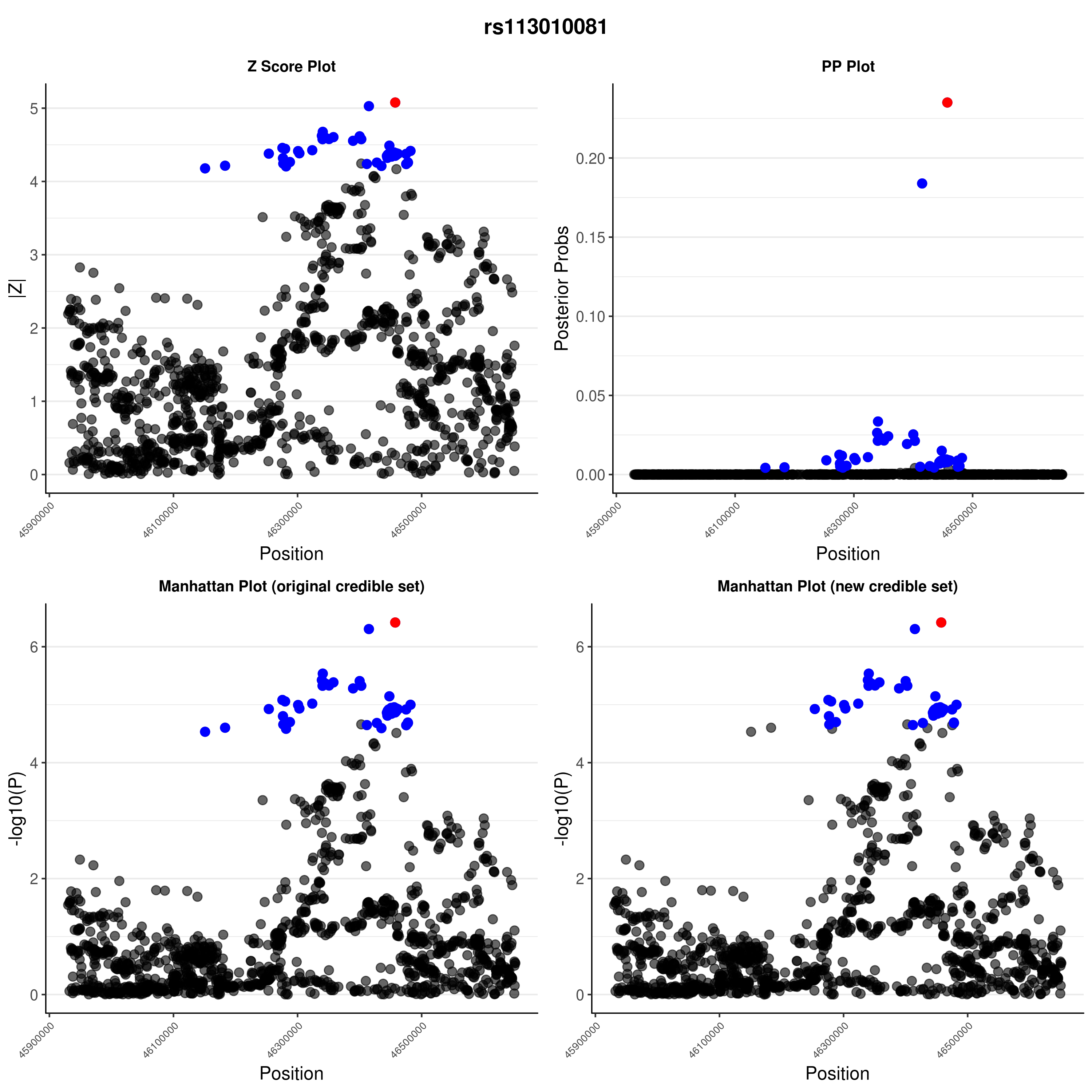

### rs113010081.png

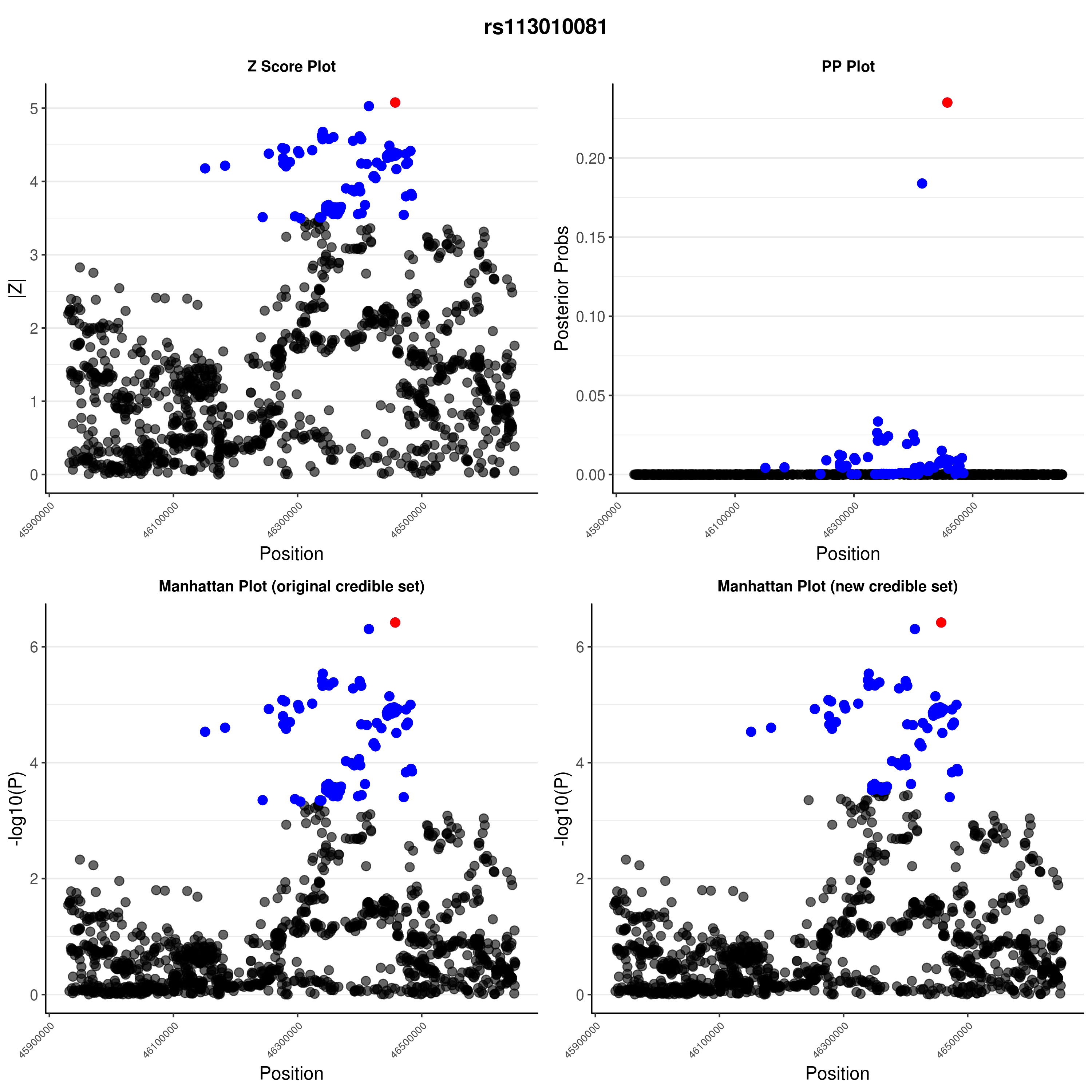

### S1 Fig

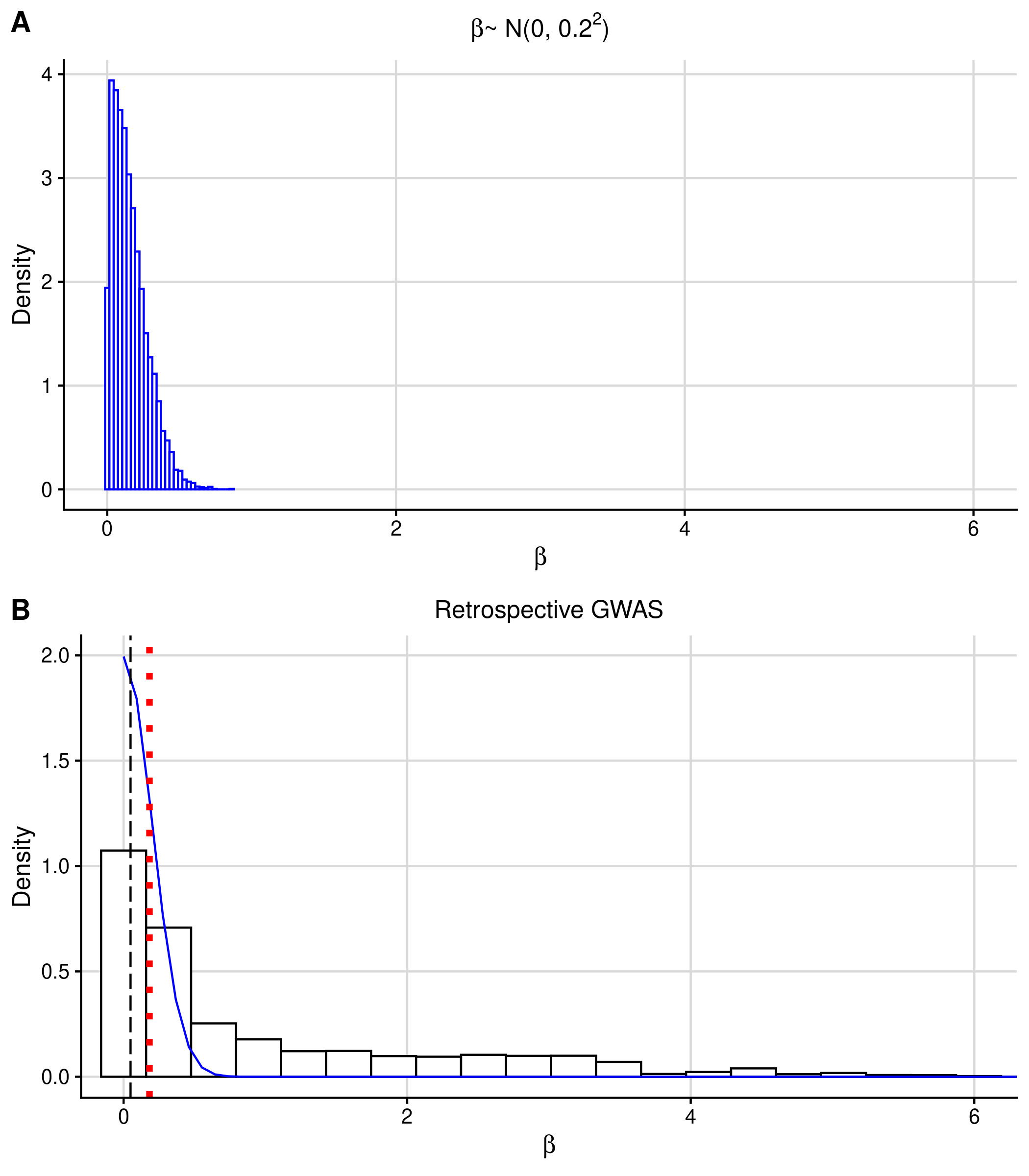

### S2 Fig

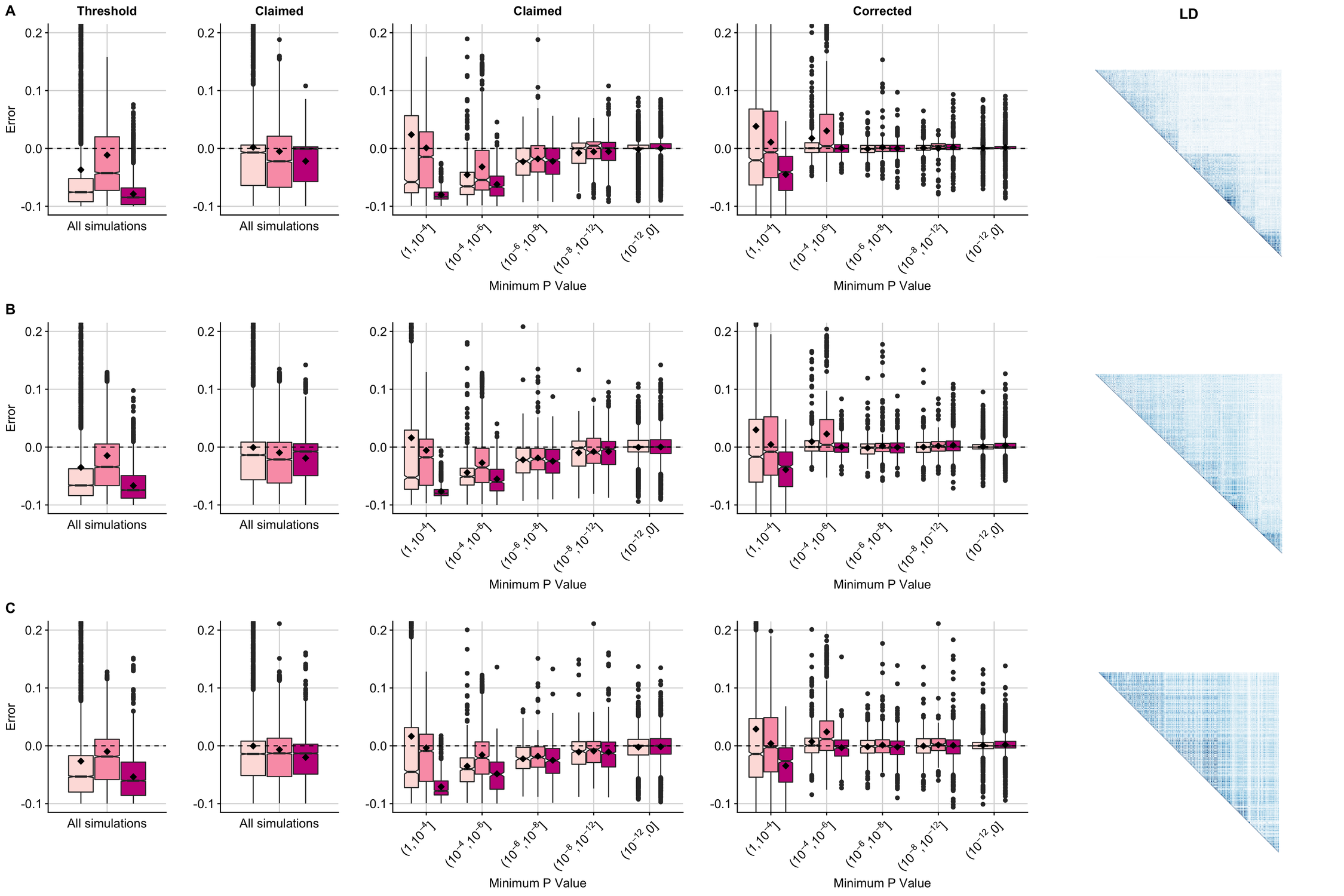

### S3 Fig

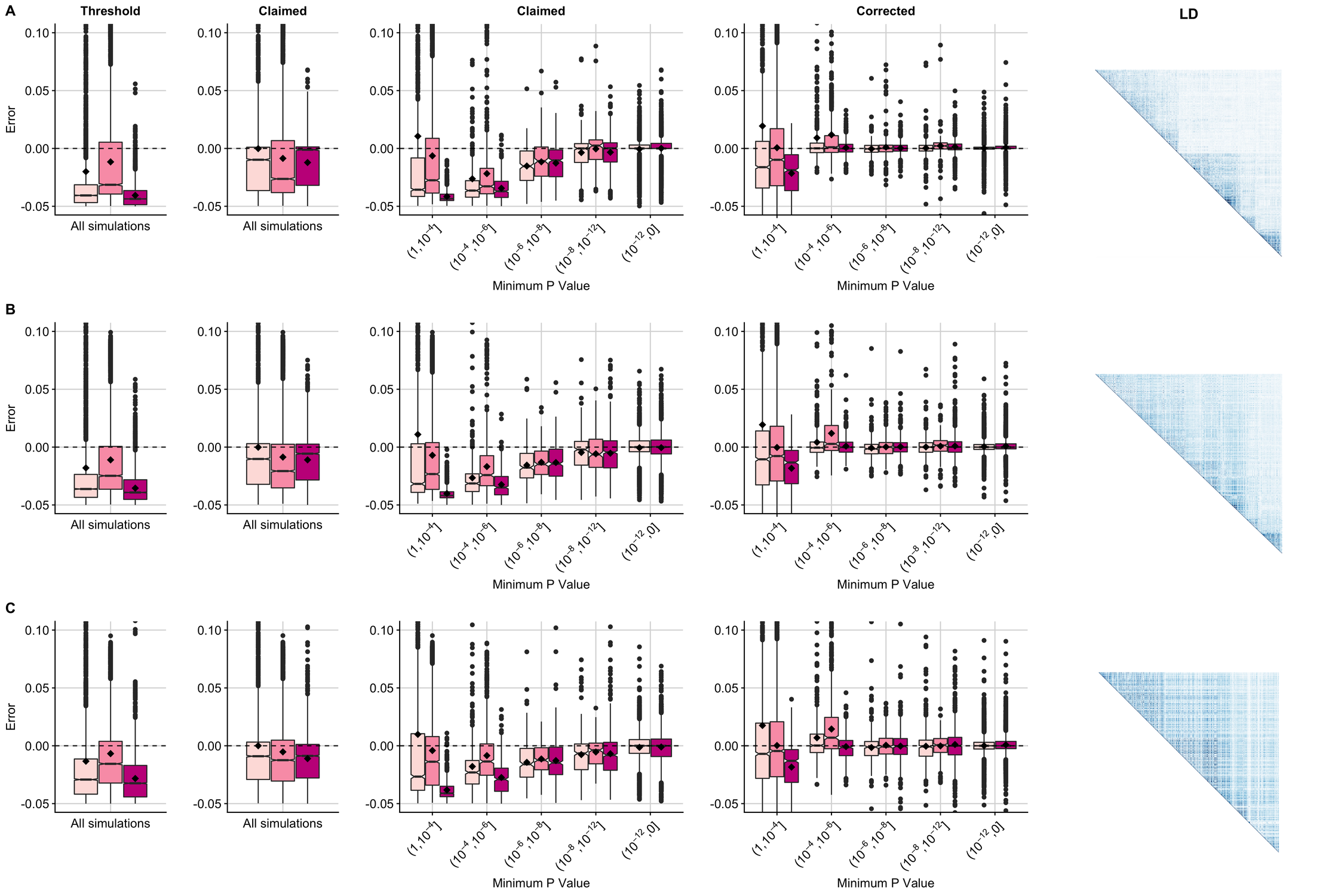

### S4 Fig

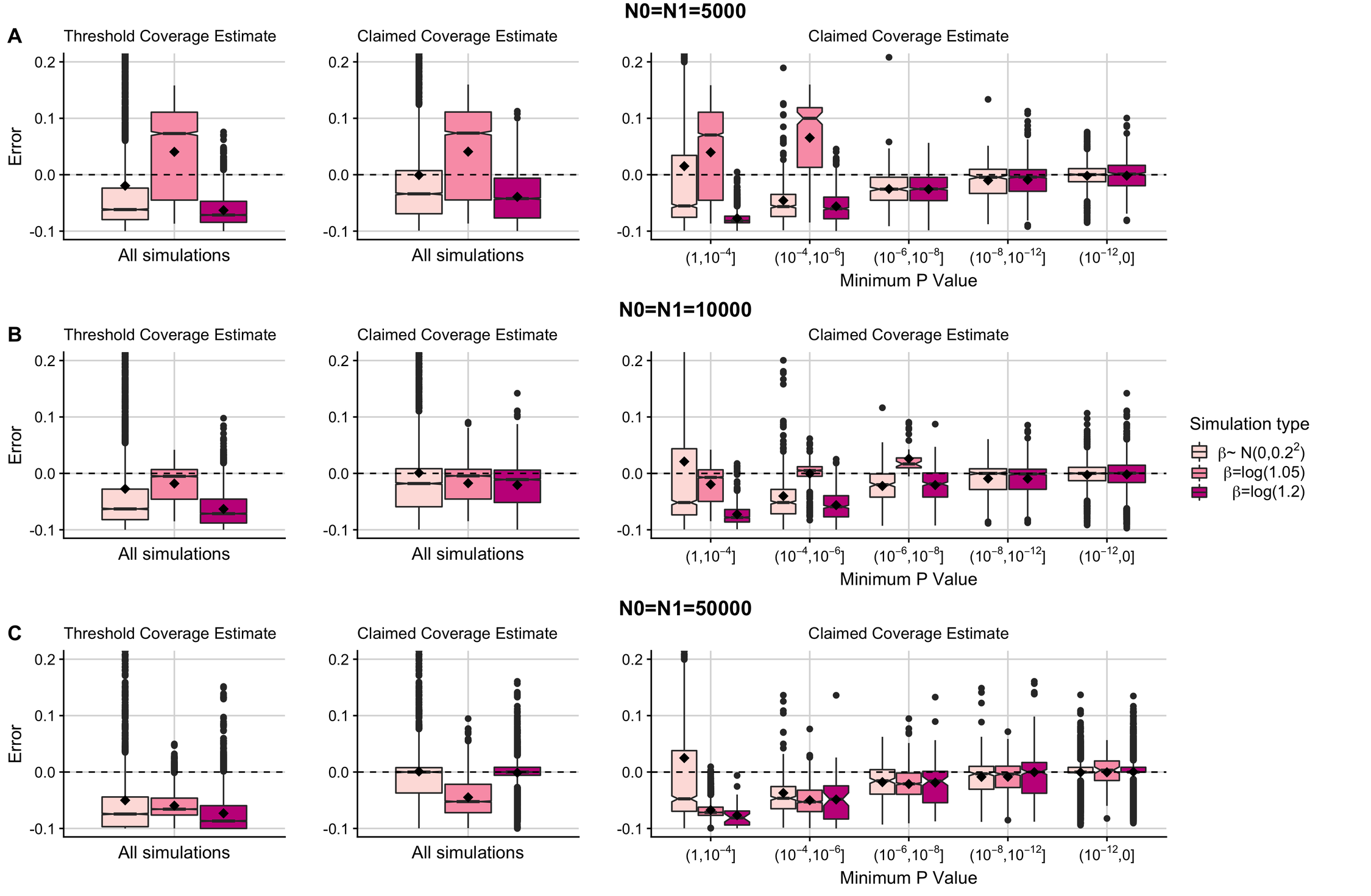

### S5 Fig

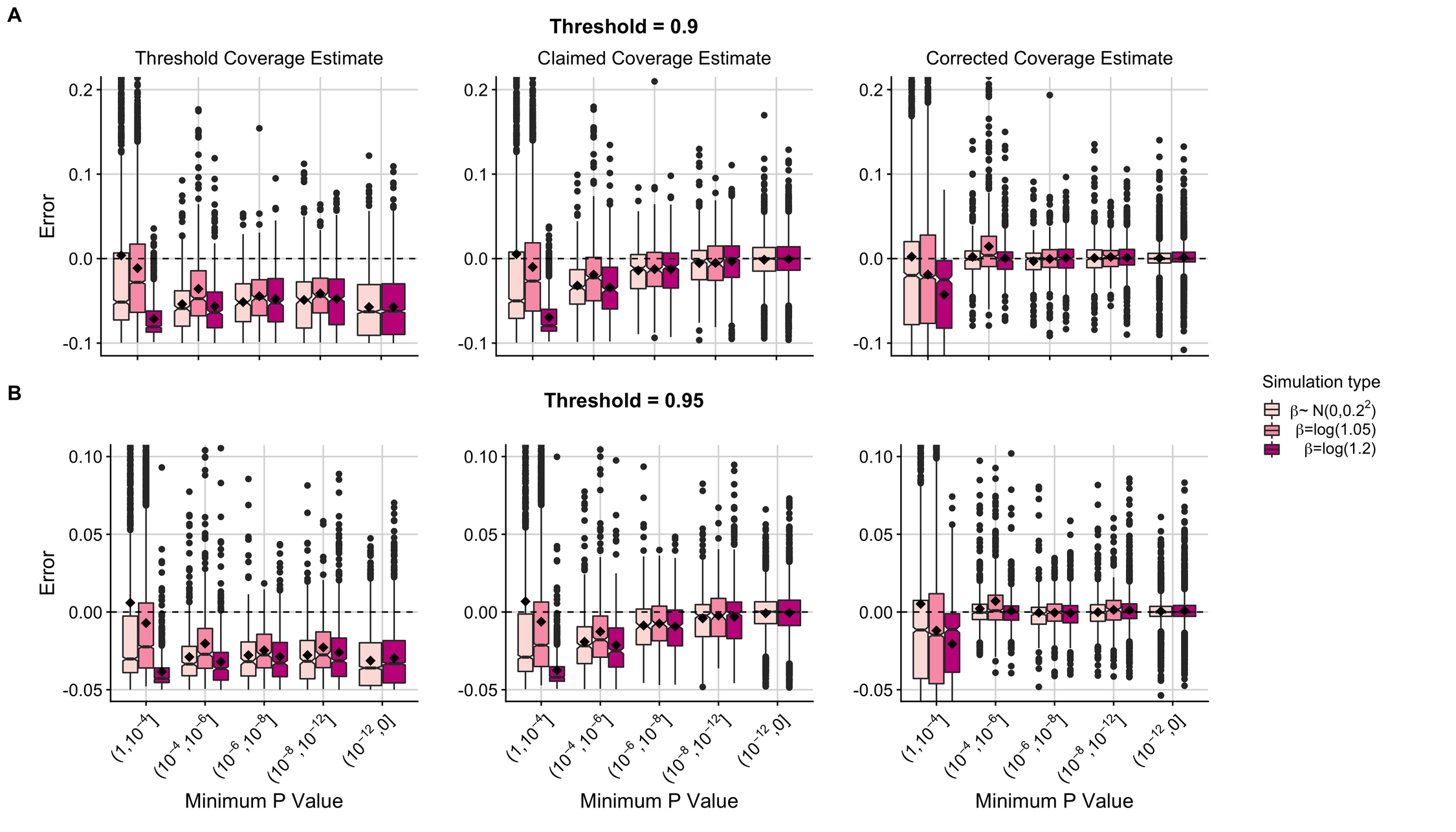

### S6 Fig

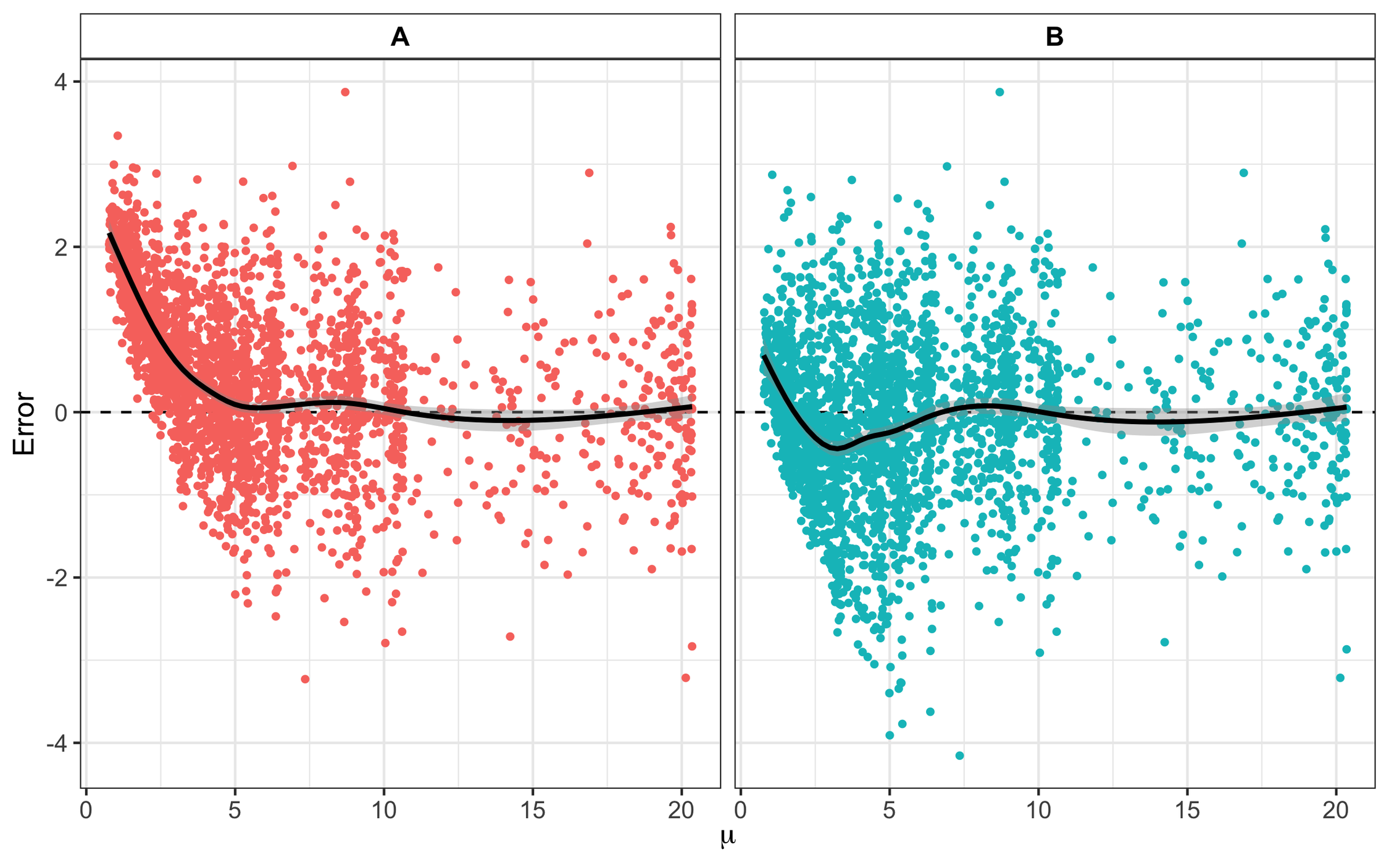

### S7 Fig

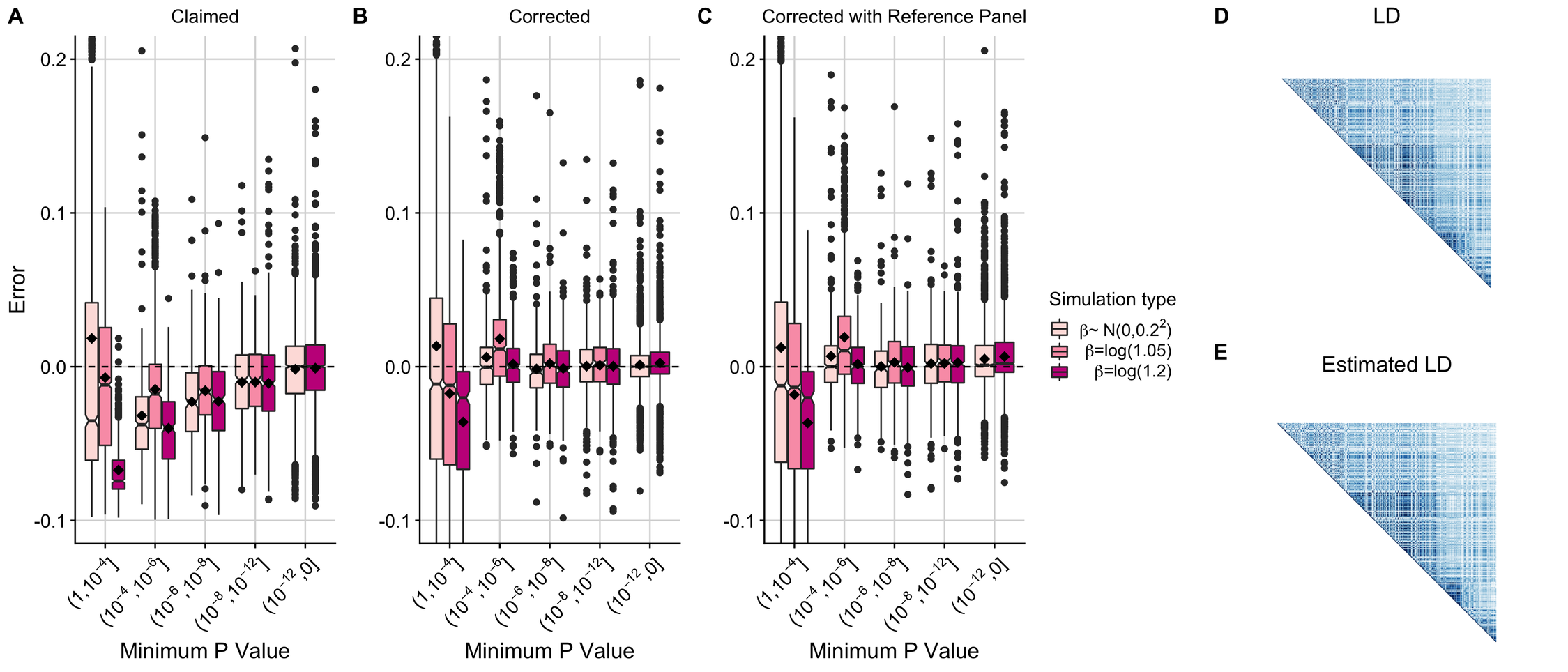

### S8 Fig

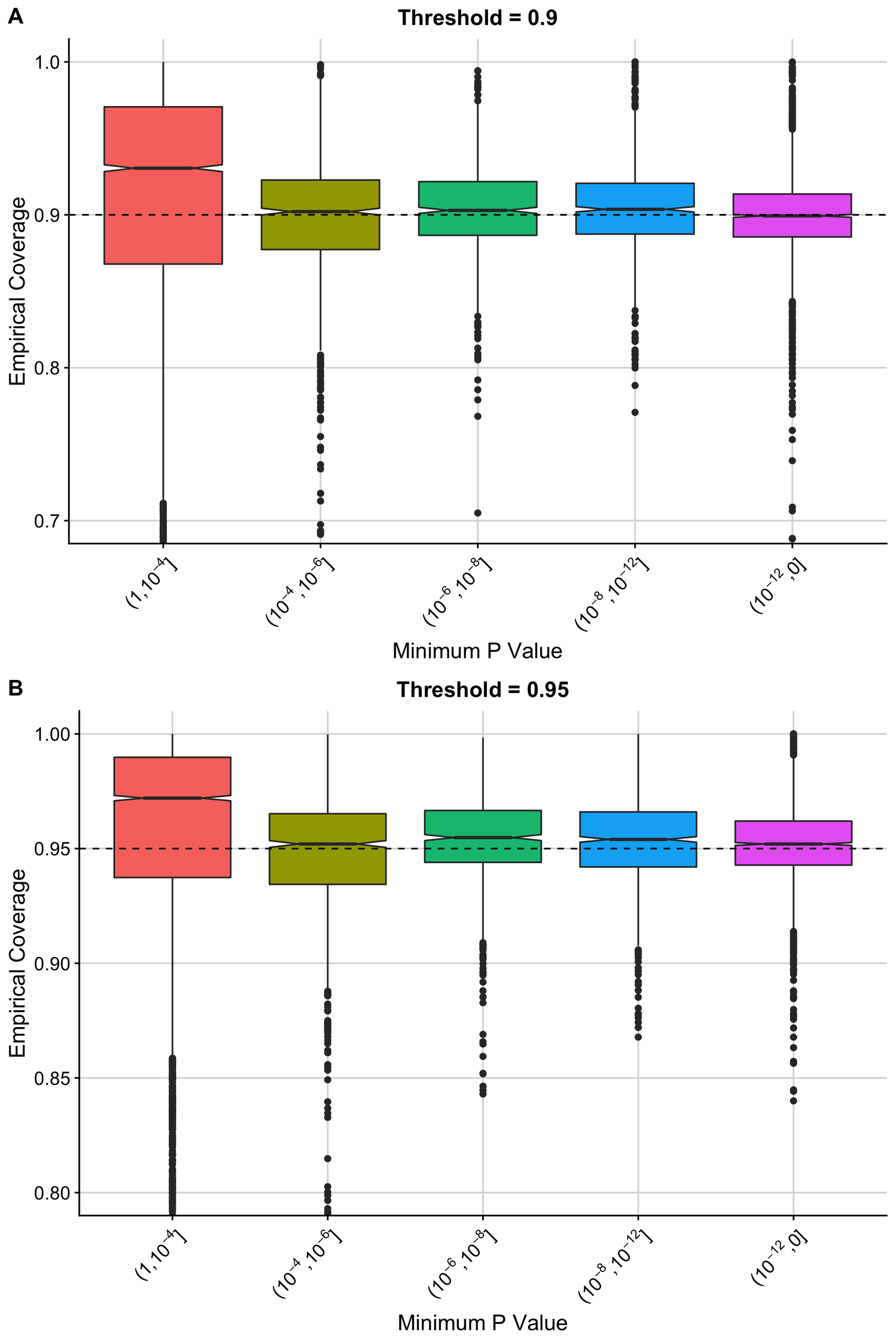

### S9 Fig

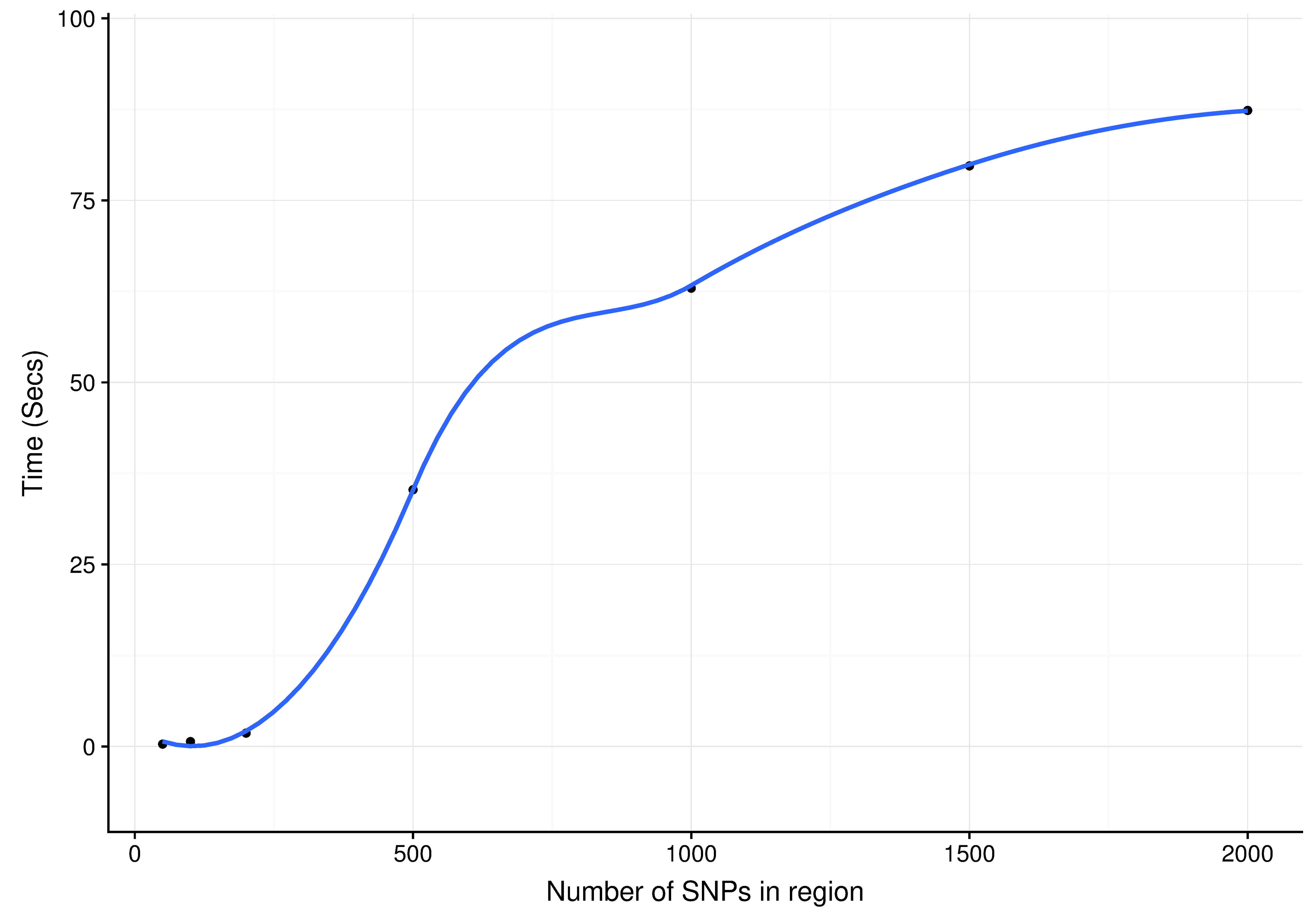

### S10 Fig

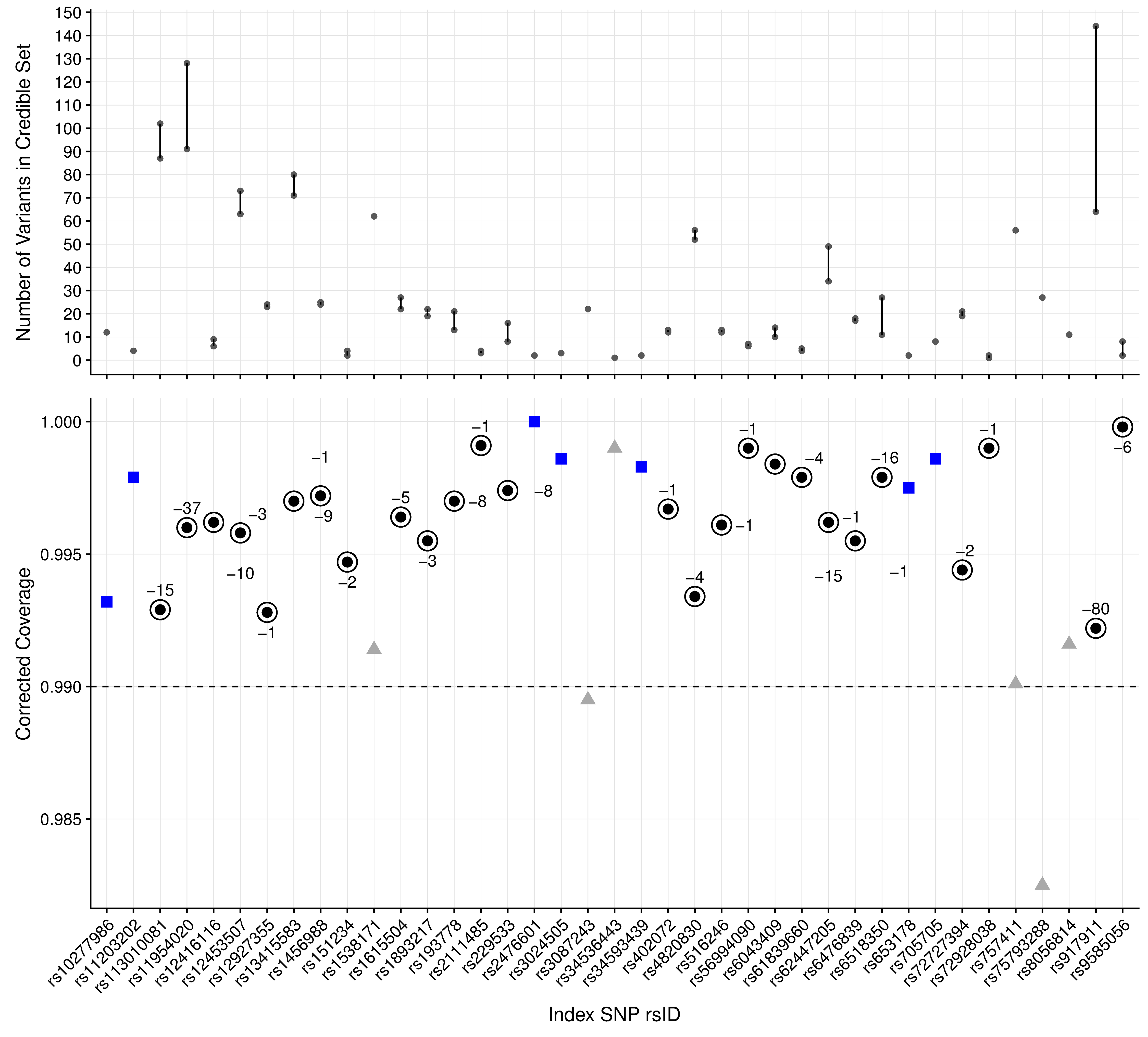

### S11 Fig

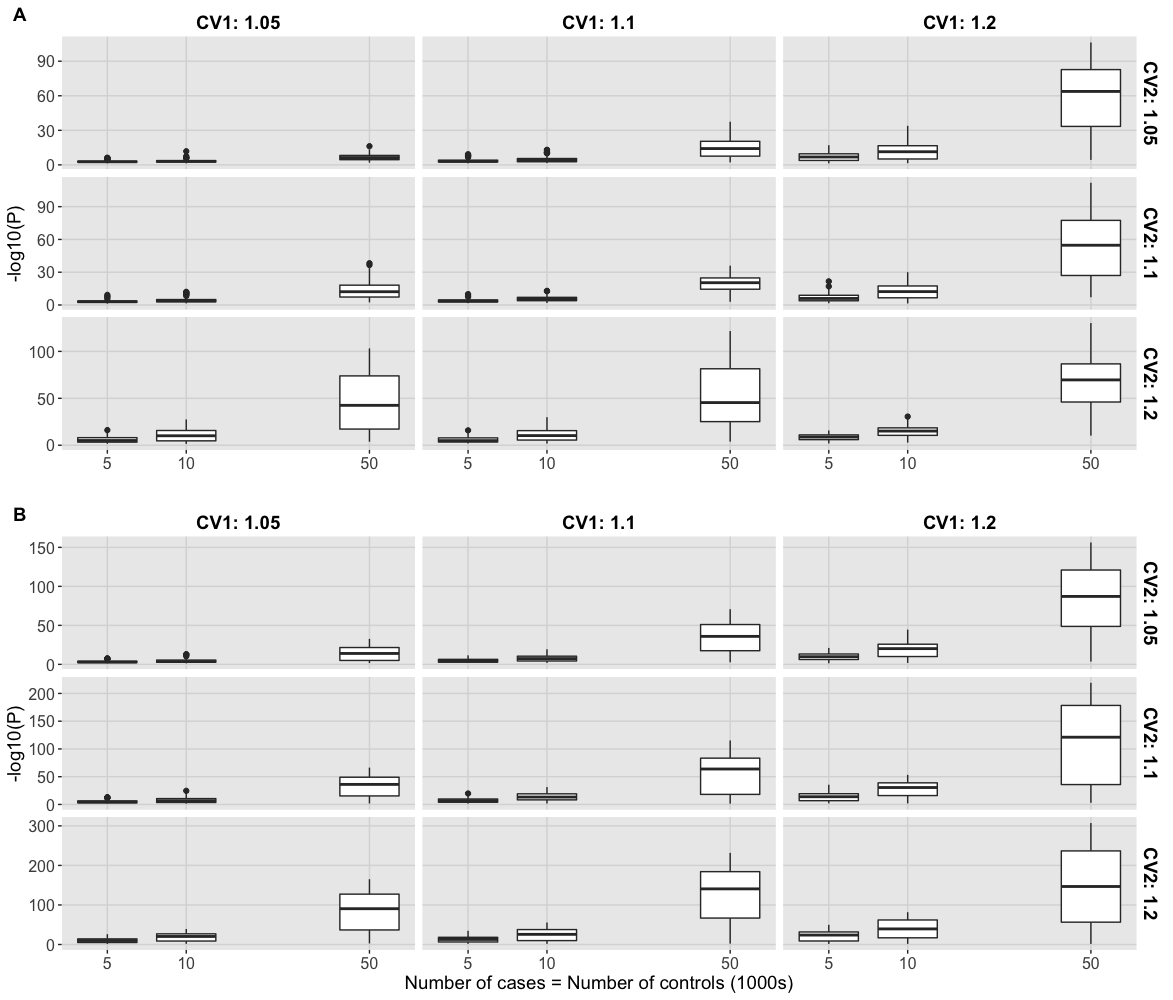
